# Deep-learning-driven tumor microenvironment profiling improves immunotherapy response prediction

**DOI:** 10.64898/2026.10.06.757143

**Authors:** Ruo Han Wang, Brooks A. Benard, Ilayda Ilerten, Sabrina Zdravkovic, Reem Al-Humadi, Sahar Nasr, Brooke Howitt, Andrew J. Gentles

## Abstract

Immunotherapy has transformed the management of many cancers, such as clear cell renal cell carcinoma (ccRCC), yet durable clinical benefit is achieved in only a subset of patients. Immunotherapy outcomes are governed by the multicellular tumor microenvironment (TME), through the structured and directional interplay of regulatory programs and ligand-receptor communications. Yet most existing methods for immunotherapy response prediction rely on independent signatures or cell types and fail to model the structured and directional signaling network within the TME. To address this gap, we introduce EcoNet, a computational framework that first models ecosystems consisting of co-occurring cell states, then constructs intercellular and intracellular signaling networks based on the ecosystems, and applies a graph attention network to derive embeddings from the signaling network for immunotherapy response prediction. Pretrained on 2,532 pan-cancer immunotherapy samples across multiple cohorts, EcoNet outperformed existing prediction methods by 7.5-19.9% and marker-based signatures by 4.4-21.2% in AUC, and its high-attention genes were enriched among those required for T-cell-mediated tumor killing in an independent CRISPR screen; applied to ccRCC with disease-specific fine-tuning, it stratified an independent ccRCC cohort by clinical outcome (*P* =0.024). Biologically, high-attention genes capture an interferon-responsive program that enhances antigen presentation and immune infiltration. To validate these findings, we curated a ccRCC clinical cohort profiled with CosMx single-cell spatial transcriptomics (46 samples, ~450,000 cells) and found the response-associated TME configuration identified by EcoNet localized to immune-enriched niches, supporting the response stratification. These results establish EcoNet as a generalizable framework for TME-informed therapeutic response prediction, readily extensible to other cancer types and therapeutic contexts.

## Introduction

Immunotherapy by immune checkpoint blockade (ICB), such as PD-1 and CTLA-4 inhibitors, has led to clinical benefit across diverse malignancies. However, durable responses are achieved in only a subset of patients, demonstrating substantial interpatient variability in immunotherapy outcome. For example, clear cell renal cell carcinoma (ccRCC), accounting for ~75% of renal malignancies and the majority of kidney cancer-related deaths (1), has shown promising responses to immunotherapy (2), yet durable benefit is achieved in only about one-third of patients (3, 4). Therefore, accurate prediction of immunotherapy outcomes is critical to guide treatment selection, avoid unnecessary toxicity, and maximize clinical benefit.

Gene expression biomarkers have demonstrated clinical utility in predicting immunotherapy response across multiple tumor types, enabling stratification based on immune activation signatures and checkpoint expression (5). However, most existing approaches focus narrowly on isolated molecular or cellular features, such as cytotoxic T-cell infiltration (6), interferon-*γ* signaling (7), or PD-L1 expression (8), without capturing the multicellular organization of the tumor microenvironment (TME). In reality, immunotherapy outcomes are driven by the interplay among diverse populations within the TME. For example, despite abundant CD8+ T cells infiltration, tumors can remain nonresponsive through the effects of M2-like macrophages and T-reg suppression or cancer-associated fibroblast (CAF)-driven TGF-*β*-mediated immune exclusion (5).

Comprehensively profiling the TME remains challenging, as dynamic cell states and intercellular cross-talk exhibit significant interpatient heterogeneity. We previously developed EcoTyper (9), which deconvolves bulk transcriptomes to infer transcriptionally distinct cell states and group their co-occurrence into multicellular ecosystems, termed *ecotypes*, thereby enabling large-scale reconstruction of TME organization.

However, several key biological layers remain unmodeled. First, immunotherapy outcomes are governed not only by expression profiles but by the structure and directionality of underlying regulatory programs and ligand-receptor interactions across the TME. For example, effective PD-1 blockade often requires IFN-*γ* from CD8+ T cells to activate JAK-STAT signaling and antigen presentation in tumor cells (10). Second, ecotype marker genes largely reflect co-abundance rather than regulatory control; a gene may appear enriched due to co-expression with proliferative states, limiting its value for therapeutic targeting. Third, a single ecotype assignment fails to capture the intratumoral heterogeneity that modulates immunotherapy response. We therefore need a predictive model to link ecotype mixtures, regulatory programs, and intercellular communications, to clinical response.

To this end, we developed EcoNet, a graph-based deep-learning framework that aligns transcriptional states to gene-regulatory and ligand-receptor interactions from multi-omic data and predicts immunotherapy response. From context-specific ecotypes, EcoNet reconstructs intercellular and intracellular networks, then applies a graph attention network (GAT) to learn ecotype-informed embeddings, with its attention mechanism highlighting genes that shape the TME. It then maps these embeddings to clinical outcomes with a downstream model. The EcoNet model was pretrained on 2,532 pan-cancer immunotherapy samples across multiple cohorts and outperformed existing prediction methods by 7.5-19.9% and marker-based signatures by 4.4-21.2% in AUC scores. Its attention scores recovered functionally important genes, which were enriched among those required for T-cell-mediated tumor killing in an independent CRISPR screen. EcoNet was subsequently applied to ccRCC with disease-specific fine-tuning. The fine-tuned model stratified an independent ccRCC cohort by progression-free survival (*P*-value= 0.024); high-attention genes identified core programs of immune activation and regulation.

To validate our findings from EcoNet, we curated a clinical cohort of 46 ccRCC samples from patients treated at Stanford Hospital and generated CosMx single-cell spatial transcriptomics. From these spatial maps, tumor programs highlighted by EcoNet localized to immune-enriched neighborhoods in situ. Notably, high-attention genes were most enriched in regions of elevated immune activity.

## Results

### Overview of EcoNet

Immunotherapy response is governed by how the tumor microenvironment (TME) is organized, including the cell states present and the directional signaling among them, which isolated markers or signatures do not capture (Figure 1a). To capture this organization, we developed EcoNet, a computational framework that integrates TME profiling, signaling network reconstruction, and deep-learning-based prediction to model how TME organization shapes immunotherapy outcome. Through the integration of multi-omic data, EcoNet characterizes the transcriptional programs and intercellular signaling relationships that collectively regulate tumor-immune communication and drive response heterogeneity.

**Fig. 1.**
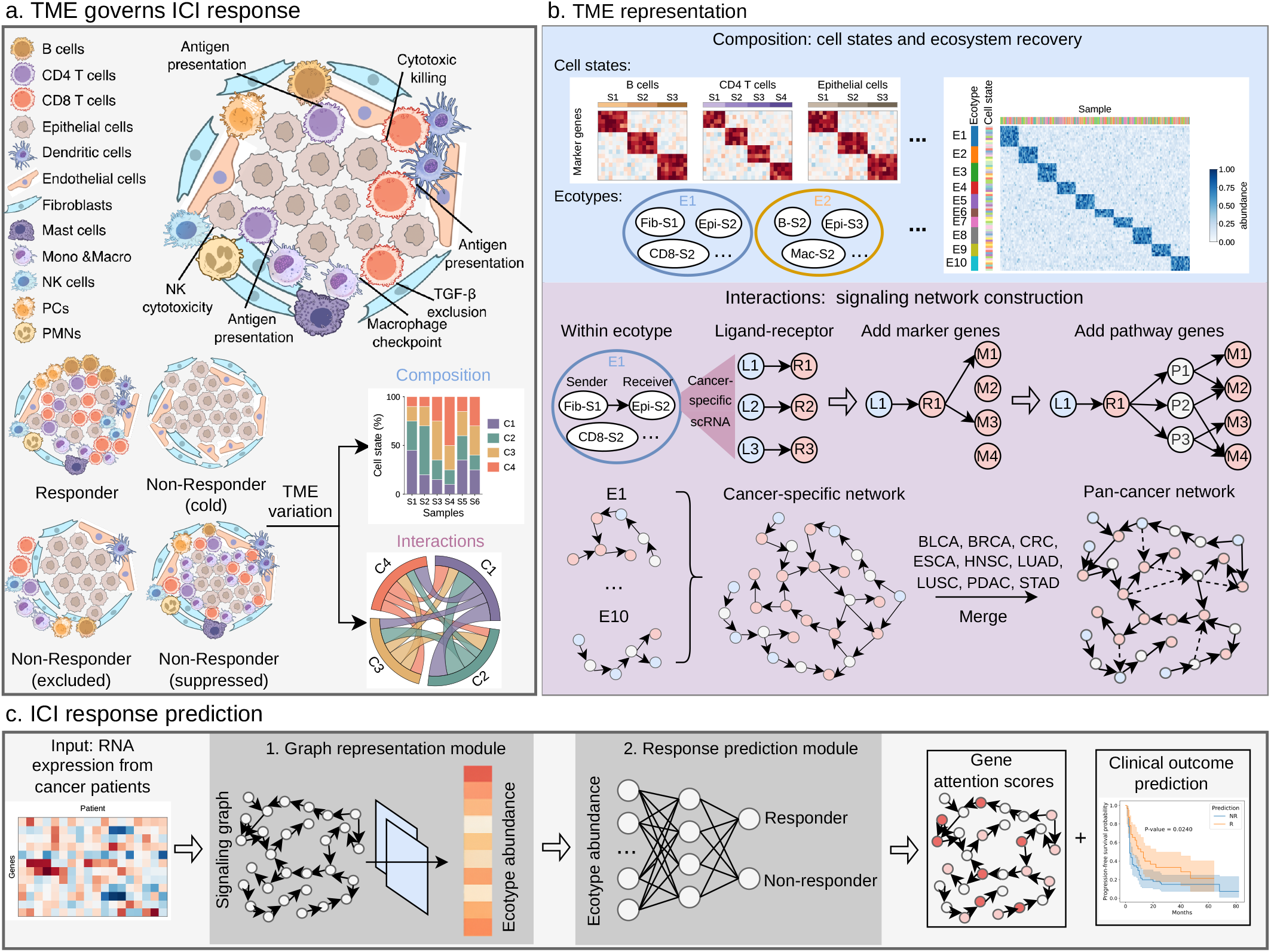
Schematic overview of EcoNet for predicting immunotherapy outcomes. **a**. The TME governs ICI response. The schematic shows the cell types and intercellular processes shaping tumor and immune cell interactions, contrasting a responder with three non-responders (cold, excluded, and suppressed). These responder and non-responder TMEs differ in the organization of cell states and the directional signaling among them. **b**. Representation of the TME through composition and interactions. For composition, we recovered cell states and ecotypes from the pan-cancer EcoTyper model. For interactions, using cancer-specific scRNA-seq data, for each ecotype we inferred regulatory relationships among the marker genes and ligand-receptor interactions based on gene expression and prior knowledge. We quantified ligand activities and retained the top five ligands to anchor the signaling networks, then incorporated pathway genes to increase connectivity and integrated networks across all ecotypes into a cancer-specific network for each of the nine cancer types. Together, these cancer-specific networks formed a pan-cancer signaling network. **c**. Immunotherapy outcome prediction. The signaling graphs, with nodes annotated by bulk RNA-seq expression, were used as input to EcoNet. EcoNet consists of a graph representation module that infers ecotype abundances and gene attention scores, and a response prediction module that predicts immunotherapy response. We trained the first module on TCGA samples and the second on pan-cancer immunotherapy cohorts.

EcoNet first represents the TME through its composition and interactions (Figure 1b). For composition, we applied EcoTyper to identify cell states within each cell type, defined by expression patterns that represent context-specific functional programs, and clustered co-occurring states across samples into higher-order ecosystems, termed *ecotypes*. Each ecotype defines a distinct TME configuration composed of interdependent immune, stromal, and tumor-intrinsic programs. For interactions, using scRNA-seq data, we inferred ligand-receptor interactions between cell states, identified ligands activating downstream marker genes, and connected them through shared pathway genes into a regulatory network for each ecotype. Integrating these ecotype-specific networks, we built a comprehensive signaling network that characterizes the gene-regulatory and signaling relationships within each cancer’s TME. We generated these networks for nine carcinomas and merged them into a pan-cancer signaling network, keeping interactions shared across cancers.

Building on the signaling network, EcoNet generates a graph for each sample, whose nodes are annotated by RNA-seq expression, and predicts immunotherapy outcome (Figure 1c). The graph representation module applies a graph attention network to generate ecotype-informed embeddings and compute gene attention scores; the response prediction module then uses a feed-forward network to derive response scores from the embeddings. We pretrained the model on pan-cancer immunotherapy cohorts to learn shared response patterns, and further fine-tuned it when adapting to a specific disease.

### Signaling networks reveal cancer-specific and shared regulatory programs across cancers

We built a signaling network for each of nine carcinomas (BLCA, BRCA, CRC, ESCA, HNSC, LUAD, LUSC, PDAC, and STAD), selected as carcinomas represented in the pan-cancer EcoTyper model (9) for which immunotherapy cohorts were available, and merged them into a pan-cancer signaling network. To build each network, we applied the published pan-cancer EcoTyper model (9) to identify cell states and carcinoma ecotypes (CE1-CE10) in each cancer’s scRNA-seq data and inferred intercellular and intracellular signaling from these data. The cancer-specific networks contained similar numbers of nodes (3,020-3,189) but more variable edge counts (12,827-21,570), reflecting cancer-specific differences in signaling interactions (Figure 2a). The merged pan-cancer network comprised 1,956 nodes and 12,219 edges, with a subset of high-degree hub genes.

**Fig. 2.**
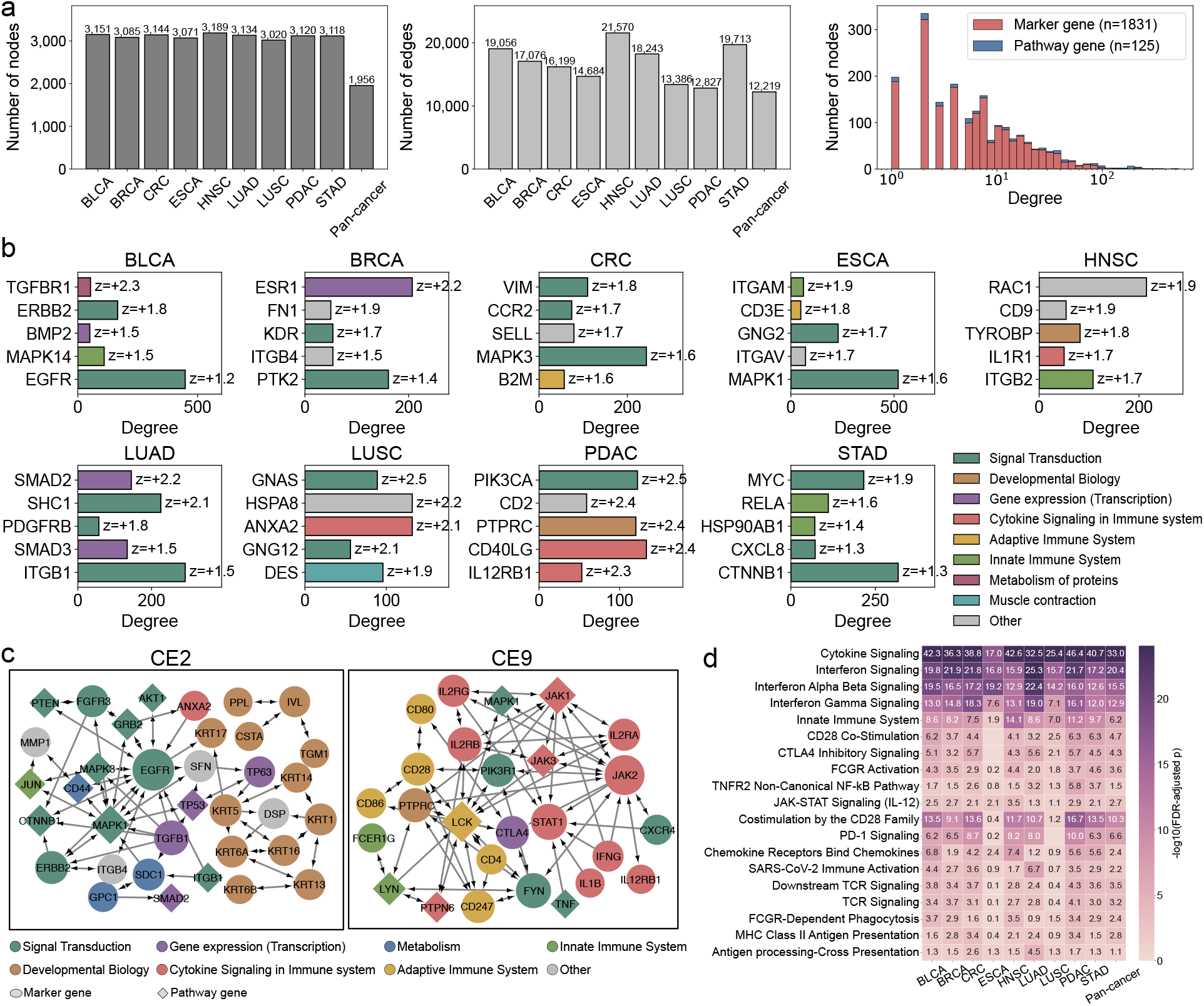
Characterization of the cancer-specific and pan-cancer signaling networks. **a**. Size and degree distribution of the networks. The left and middle barplots show the number of nodes and edges for each of the nine cancer-specific networks and the pan-cancer network. The right panel shows the degree distribution of the pan-cancer network, with nodes grouped into cell-state marker genes and intermediate pathway genes. **b**. Cancer-specific hub genes. For each cancer, the five most specific hub genes are shown, with degree on the x-axis and each bar annotated by its density-normalized degree z-score across the nine cancers. Higher z-scores indicate greater cancer-specific connectivity. Bar color indicates the Reactome functional class. **c**. Signaling subnetworks of ecotypes CE2 and CE9. Each network shows the top 40 genes by degree within the ecotype in the pan-cancer network. Node size scales with degree, node color indicates the Reactome functional class, and node shape distinguishes marker genes (circles) from pathway genes (diamonds). Directed edges indicate signaling direction. **d**. Immune-pathway enrichment of the CE9 network across cancers. The heatmap shows Reactome immune pathway enrichment of the CE9 network gene set in each cancer and in the pan-cancer network, with color indicating the enrichment significance.

The cancer-specific networks captured regulatory programs of each tumor type. To identify the hub genes most specific to each cancer, we adjusted each gene’s degree for the density of its network and z-scored these adjusted degrees across the nine cancers. The top five hub genes are shown for each tumor type (Figure 2b). These top hubs aligned with known cancer biology, such as *ESR1*, the key driver of luminal breast cancer (11); *EGFR*, a receptor tyrosine kinase amplified in bladder cancer (12); and *CTNNB1*, a common Wnt-pathway alteration in gastric cancer (13). These hubs spanned diverse functional classes, including signal transduction, cytokine signaling, and adaptive and innate immune programs.

In the pan-cancer network, we identified regulatory programs shared across tumor types. Within this network, each ecotype retained a coherent program of genes present across the nine cancers (Figure 2c). For example, the CE2 module centered on epithelial and developmental genes, while the CE9 module was composed largely of immune and cytokine-signaling genes, including *JAK1, JAK2, STAT1, LCK*, and *IFNG*. Each of the ten ecotypes carried a distinct, functionally coherent program, spanning immune, stromal, developmental, and contractile processes (Supplementary Figure S1). These programs were consistent across cancers, indicating that our pan-cancer network recovers regulatory programs conserved across tumor types.

Given that CE9 comprises immune genes and has been reported as an immunotherapy-favorable ecotype across cancers (9), we characterized its immune programs. Specifically, we performed pathway enrichment analysis on the CE9 genes with all network genes as background. CE9 was enriched for immune pathways, including interferon, cytokine and chemokine signaling, and antigen presentation, across all nine carcinomas and in the pan-cancer network (Figure 2d). These pathways are central to anti-tumor immunity and immunotherapy response, showing that the reconstructed CE9 network captures the immune program consistent with its association with favorable immunotherapy outcomes.

### EcoNet predicts pan-cancer immunotherapy response and identifies functionally important genes

EcoNet links immunotherapy outcome to TME patterns through a deep-learning model (Figure 3a). The model encodes bulk expression on the ecotype-informed signaling network, derives a latent ecosystem embedding via a graph attention network, and outputs the likelihood of response from the embedding. The attention network also assigns each gene an attention score, reflecting each gene’s contribution to characterizing the TME. It was pretrained on pan-cancer immunotherapy cohorts to learn shared response patterns (see Methods). The design incorporates biological priors, including cell states, cell-cell communications, and transcriptional programs, to guide learning.

**Fig. 3.**
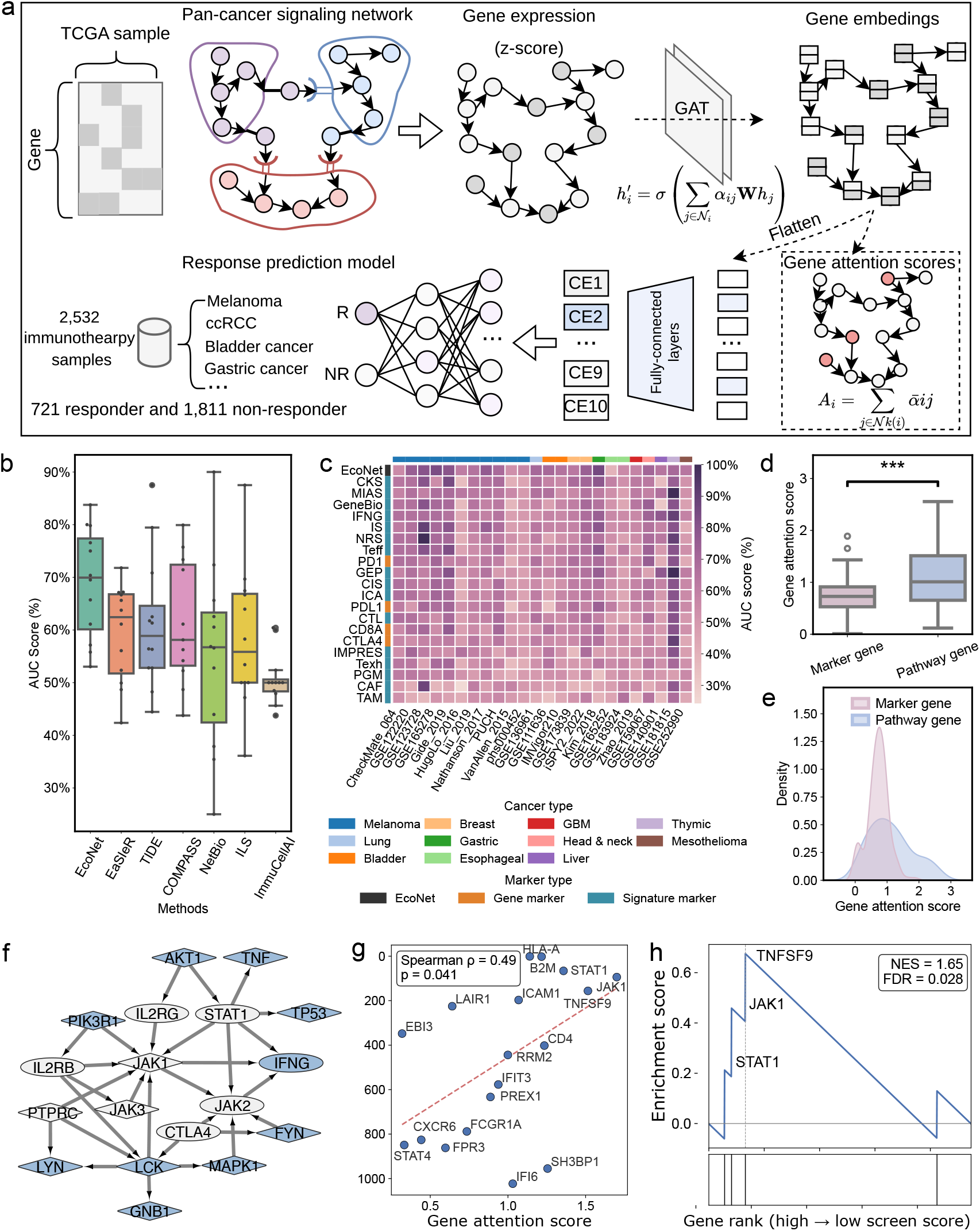
Performance of EcoNet in predicting pan-cancer immunotherapy outcomes. **a**. The architecture and training process of EcoNet. The input is the constructed gene signaling graph, where each node is assigned features derived from bulk RNA-seq expression profiles. EcoNet consists of two modules, a graph representation module that predicts the abundance of ten ecotypes while assigning attention scores to genes, and a response prediction module that predicts immunotherapy response from the inferred ecotype abundances. The first module is trained on TCGA pan-cancer samples, and the second is trained on pan-cancer immunotherapy cohorts. **b**. Comparison of EcoNet with existing immunotherapy response prediction tools, ordered by median AUC score. Boxplots summarize the distribution of AUC scores across the test datasets applicable to all tools. **c**. Comparison of EcoNet with individual gene and signature markers of immunotherapy response, ordered by median AUC score. The heatmap shows AUC scores across the test datasets, with cancer type and marker type annotated. **d**. Distribution of gene attention scores for marker genes and pathway genes within the CE9 network. **e**. Comparison of gene attention scores between marker genes and pathway genes within the CE9 network. **f**. Signaling paths among the genes with the highest EcoNet attention scores in the CE9 network. High-attention genes are shown in blue and the connecting genes in grey, circles and diamonds denote marker genes and pathway genes, respectively, and directed edges represent regulatory relationships. **g**. Correlation between EcoNet attention scores and gene ranks in an independent CRISPR immune-killing screen. Each point is a CE9 network gene present in the screen, with EcoNet attention score on the x-axis and its rank by requirement for T-cell-mediated tumor killing on the y-axis; the dashed line shows the linear fit, and the Spearman correlation is annotated. **h**. Gene set enrichment analysis of the top 30 CE9 attention genes using the published 947-gene CRISPR screen ranking. Four of these genes were present and included in the analysis. The running enrichment score is shown above and the positions of these four genes below, with the normalized enrichment score and FDR annotated.

Using a leave-one-cohort-out strategy across independent immunotherapy cohorts (Supplementary Table S1), we benchmarked EcoNet on immunotherapy response prediction against existing prediction methods and marker-based baselines (see Methods). Since the response prediction component of TIDE was established specifically on melanoma and non-small cell lung cancer (NSCLC) cohorts (5), we compared the prediction methods on cohorts of these two cancer types. EcoNet achieved the highest median AUC score of 69.9%, outperforming the other methods by 7.5% *−*19.9% (Figure 3b). Among the marker-based baselines, evaluated across 11 cancer types, EcoNet again performed best, exceeding all markers by 4.4%*−*21.2% (Figure 3c). Signature markers generally outperformed single-gene markers, and EcoNet outperformed both, consistent with response depending on the whole TME rather than on isolated features. An ablation analysis showed that each component contributes to this performance. Removing the GAT, ecotype supervision, graph edges, or the learned prediction head reduced the median AUC from 69.9% to 66.8%, 64.0%, 62.7%, and 56.4%, respectively (Supplementary Section S2.1 and Supplementary Figure S2).

We then examined the attention scores EcoNet assigns to genes, which quantify each gene’s contribution to the learned representation. In the immune-active ecotype CE9, pathway genes received higher attention than marker genes (*P* = 8.6 *×* 10^*−*5^; Figure 3d,e; Supplementary Figure S3 for other ecotypes), indicating that the model weights more the intermediate signaling genes connecting ecotype programs. Tracing signaling paths among the ten highest-attention genes in the CE9 network, we found they linked inflammatory signaling, chemokine production, antigen presentation, cytotoxic activation, and checkpoint regulation (Figure 3f).

To evaluate whether these high-attention genes are functionally important, we compared them against an independent genome-scale CRISPR screen for T-cell-mediated tumor killing (14). In this screen, genes are ranked by the extent to which their knockout reduces killing, with top-ranked genes (smaller rank numbers) most needed for tumors to be killed. We compared EcoNet attention scores with these ranks for the CE9 network genes present in the screen. Attention score correlated with screen rank, with higher-attention genes more strongly required for killing (Spearman *ρ* = 0.49, *P* = 0.041; Figure 3g). Of the 30 highest-attention CE9 genes, four were present in the published 947-gene ranking from Patel et al. Exploratory GSEA of these four genes showed enrichment toward the top of the list (NES= 1.65, FDR= 0.028; Figure 3h), with *STAT1, JAK1*, and *TNFSF9* ranking highest among them. Together, these results show that within CE9, EcoNet assigns its highest attention scores to genes required for immune killing, linking them to the mechanism underlying immunotherapy response. We further showed that attention scores were robust to training-set composition (Supplementary Figure S4).

### ccRCC ecotype analysis identifies RE2 as an immunotherapy-favorable ecosystem

We next turned to ccRCC as a disease-specific application. Since ccRCC was excluded from the pan-cancer EcoTyper discovery cohort based on its distinct housekeeping gene expression profile (9), we built a ccRCC-specific model to capture its TME organization. Applying EcoTyper to the 614 TCGA-KIRC bulk RNA-seq samples resolved eleven distinct ecotypes within the ccRCC TME, denoted RE1-RE11, distinct from the pan-cancer carcinoma ecotypes CE1-CE10. (Figure 4a). Among these ecotypes, RE8 was strongly enriched in adjacent normal tissues and represented a nonmalignant context. A UMAP plot of samples based on ecotype marker genes showed clear separation among ecotypes, and RE8 formed a distinct cluster apart from the tumor-dominated ecotypes (Supplementary Figure S5). We evaluated cell-state recovery by mapping single cells in independent scRNA-seq cohorts to the inferred cell states. Our ccRCC model recovered a higher proportion of cells than either the pan-cancer model or a ccRCC model built using the original EcoTyper workflow (Supplementary Result S2.2 and Supplementary Table S2 and S3).

**Fig. 4.**
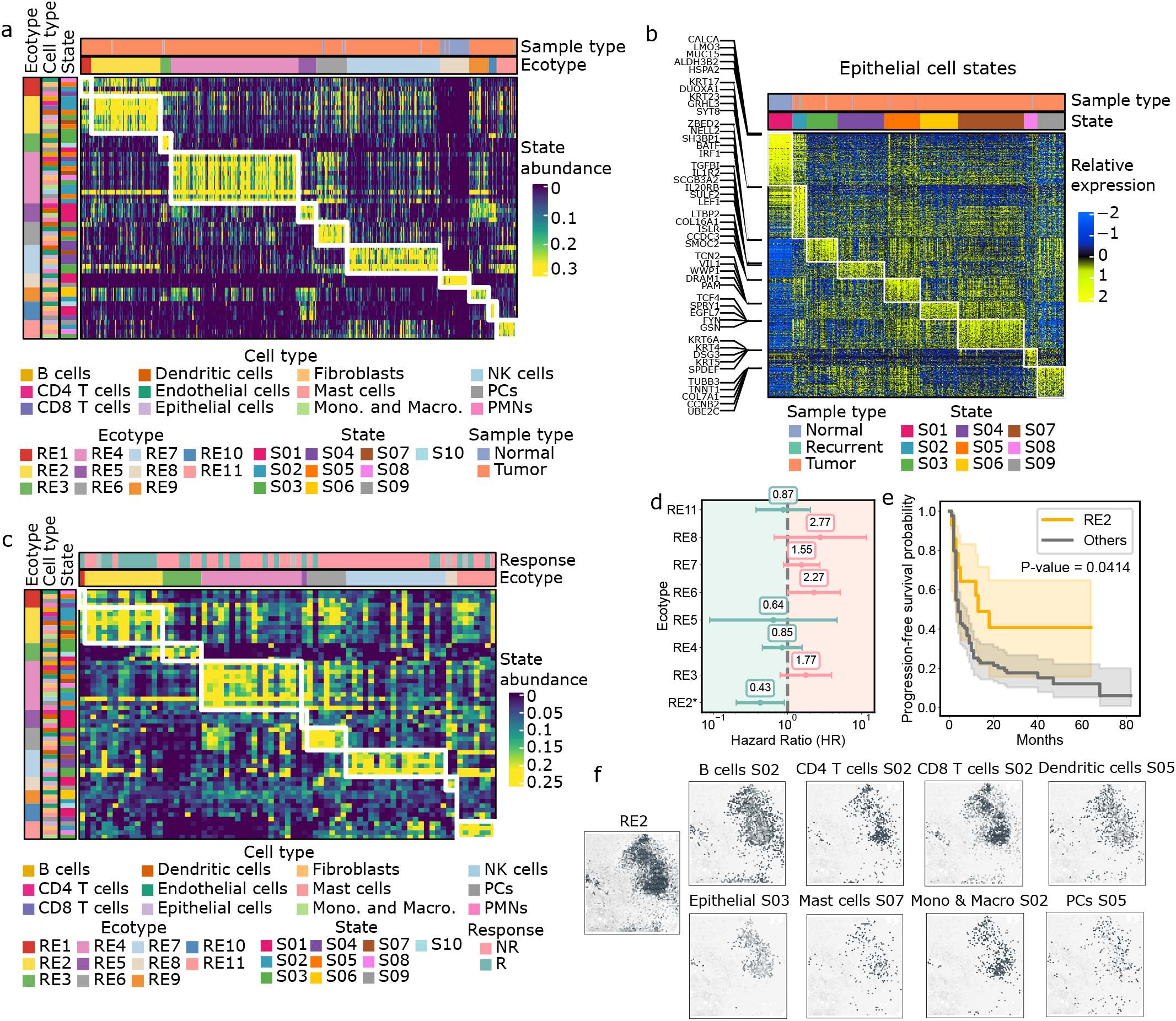
Discovery of ccRCC ecotypes and association of RE2 with immunotherapy response. **a**. Heatmap of cell state abundance across TCGA-KIRC samples, organized into eleven ecotypes. Each row represents a cell state and each column a sample, and color indicates state abundance. The three vertical bars to the left annotate the ecotype assignment, the corresponding cell type, and the specific state identity of each row. The horizontal bars above indicate the sample type and the assigned ecotype. **b**. Heatmap of epithelial cell states across TCGA-KIRC samples, organized into nine distinct states. Each row represents a marker gene and each column a sample, with color intensity reflecting relative expression. The top bar indicates epithelial state assignments, and five representative marker genes for each state are highlighted on the left. **c**. Recovery of ecotypes in the ccRCC Leuven cohort. The heatmap shows the cell state abundance profile across Leuven samples, with rows corresponding to cell states and columns to samples. Color indicates state abundance. The three vertical bars to the left annotate the ecotype assignment, the corresponding cell type, and the state identity of each row. The horizontal bars above indicate the clinical response annotation and the assigned ecotype. **d**. Cox proportional hazards regression for ecotypes in the Leuven cohort. The dots represent the estimated HR for progression, and horizontal lines indicate 95% confidence intervals. The vertical dashed line marks HR = 1, bars to the left of 1 indicate protective associations, whereas bars to the right indicate increased risk. **e**. Progression-free survival analysis comparing patients assigned to RE2 versus all other ecotypes, with P-value annotated. **f**. Recovery of RE2 and its cell states in spatial transcriptomics data (GSM5924036). Dot intensity reflects the abundance of RE2 and corresponding cell states, overlaid on tissue sections with epithelial regions marked in grey.

We identified 70 cell states across twelve cell types. For example, the epithelial population comprises nine states (Figure 4b). Epithelial state S01 was enriched in adjacent normal samples and marked by *MUC15, ALDH3B2*, and *HSPA2*, consistent with a differentiated epithelial program (15, 16). Tumor-associated epithelial states were heterogeneous with discrete transcriptional programs. Among them, S03 shows an IFN-responsive and immune-interacting epithelial state marked by *ZBED2, NELL2, BATF, IRF1*, and *SH3BP1* (17, 18). Marker genes for all cell types are shown in Supplementary Figure S6. The epithelial state markers were enriched for cell migration and adhesion programs, and the normal-enriched state additionally showed renal absorption terms (Supplementary Result S2.3, Supplementary Figure S7). These markers were expressed most highly in KIRC across TCGA cancer types (Supplementary Result S2.4, Supplementary Figure S8). To assess the robustness of these cell states, we applied the ccRCC EcoTyper model to independent scRNA-seq cohorts, and the model significantly recovered 57 of 70 of these cell states (81.4%; Supplementary Result S2.5, Supplementary Table S4, Supplementary Figure S9).

We applied the ccRCC EcoTyper model to an independent dataset of 98 samples for which immunotherapy response data were also available (the Leuven ccRCC immunotherapy cohort) (2), to recover cell states and ecotypes. We observed enrichment of RE2 among responders, suggesting that it defines an ICI-responsive ecotype. Of the RE2-assigned samples, nine were responders, four were non-responders, and one lacked response information (Figure 4c). To further evaluate this association, we performed Cox proportional hazards modeling for progression-free survival (PFS) and found RE2 was significantly associated with clinical benefit (Figure 4d). In addition, grouping samples by RE2 versus others showed improved PFS for RE2 (*P*-value = 0.0414; Figure 4e), whereas the immunotherapy-favorable ecotype CE9 from the pan-cancer EcoTyper model (9) achieved a *P*-value of 0.0741 (Supplementary Figure S10). We also showed PFS stratified by each ecotype in Supplementary Figure S11. After fine-tuning on ccRCC cohorts, EcoNet was the only method to significantly stratify the Leuven cohort by both PFS and OS and achieved the highest response-prediction AUC among all compared methods (Supplementary Figure S12, S13, and S14). Consistent with this association, RE2 showed stronger epithelial-immune communication than other ecotypes in single-cell data (Supplementary Result S2.6, Supplementary Figure S15-S17).

To examine the spatial correlation of the cell states in RE2, we recovered ecotypes and constituent cell states in a ccRCC 10x Visium dataset (19). We then computed per-spot ecotype abundance and observed variable RE2 abundance across samples, with mean per-spot fractions ranging from 0.076 to 0.240 (Supplementary Figure S18), and subsequently mapped the spatial localization of ecotypes and cell states. In one representative sample (GSM5924036), RE2 states co-localized within shared neighborhoods (Figure 4f, Supplementary Figure S19), suggesting spatial proximity compatible with interactions. Additional RE2-abundant samples (mean abundance *>* 0.15) also exhibited similar patterns (Supplementary Figure S20-S26).

### EcoNet predicts ccRCC immunotherapy outcome and identifies RE2 regulatory drivers

Based on the ecotype abundance derived for the scRNA-seq samples (Supplementary Figure S15), we selected the sample with the highest abundance for each ecotype to build an ecotype-specific intercellular and intracellular signaling network, and subsequently combined these ecotype-specific networks into a global ccRCC TME network (see Methods).

To illustrate the RE2 program, we highlighted the subnetwork between epithelial S03 and CD8 T cells S02 (Figure 5a), whose markers included major targets of ICI therapies. Epithelial cells presented ligands such as CD274 (PD-L1), CD86, and CCL3 that engage T-cell receptor and co-stimulatory modules, with downstream targets including LCK, ZAP70, and CD247; in turn CD8 T cells were predicted to signal back with IFNG and TNF, activating epithelial JAK-STAT and interferon response genes such as *STAT1, JAK1*, and *OAS1*. Together these circuits indicate coordinated checkpoint and cytokine communication that structures epithelial-immune crosstalk in RE2.

**Fig. 5.**
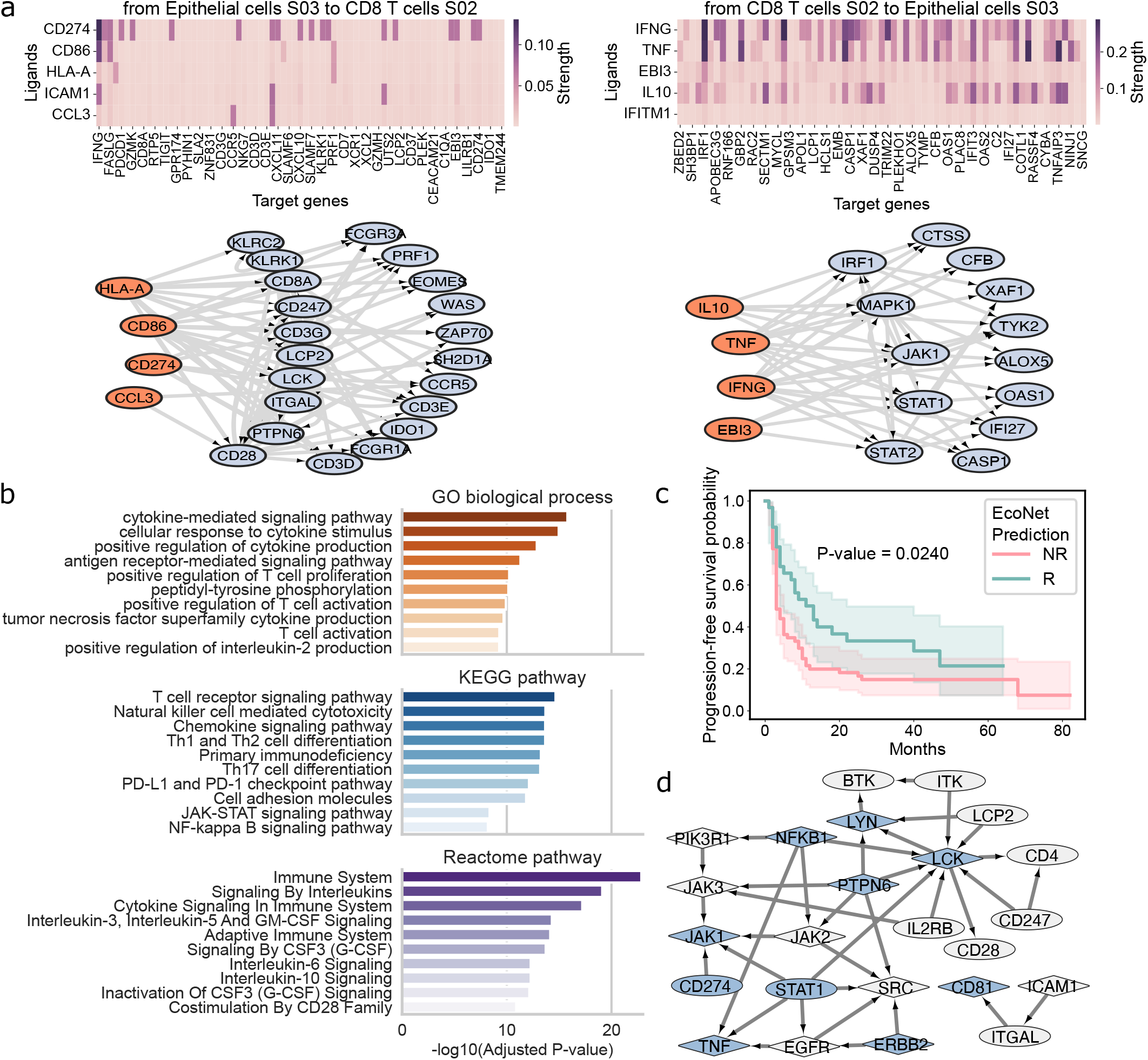
EcoNet predicts ccRCC immunotherapy outcomes and identifies key regulatory genes. **a**. Signaling paths between epithelial cells S03 and CD8 T cells S02 from the ICI-responsive ecotype RE2. The upper heatmaps show the predicted activity of the top five ligands in the sender populations on marker genes of the receiver populations. The lower networks display the signaling paths, with ligands shown as orange nodes and regulated target genes as blue nodes. Arrows indicate the regulatory relationships between the genes. **b**. Functional enrichment of RE2-specific hub genes. Barplots show the top ten enriched terms across GO biological process, KEGG pathway, and Reactome pathway categories. **c**. Progression-free survival analysis of the Leuven cohort stratified by EcoNet-predicted responder and non-responder groups, with P-value annotated. **d**. Signaling paths among the genes with the highest EcoNet attention scores in the RE2 network. High-attention genes are shown in blue and the connecting genes in grey, circles and diamonds denote marker genes and pathway genes, respectively, and directed edges represent regulatory relationships.

To characterize RE2-specific network features, we compared node centrality between the global and RE2 networks (Supplementary Figure S27). In the global network, central nodes were generic signaling hubs such as *UBC* and *MAPK1*, consistent with broad receptor-tyrosine-kinase and cytoskeletal signaling across the TME. In contrast, the RE2 network shifted toward immune-activation hubs, with *LCK* and *CD28* among the most connected nodes and *JAK2* indicating cytokine-responsive signaling. To further identify pathways emphasized in RE2, we performed enrichment analyses using GSEApy on the top 50 high-degree genes from the RE2 network that were not top-ranked globally (Figure 5b). The results highlighted cytokine-mediated signaling, antigen receptor and T-cell receptor signaling, chemokine programs, NK-cell cytotoxicity, and PD-1/PD-L1 checkpoint. Together, these analyses define an RE2-specific regulatory niche dominated by interferon-driven activation and checkpoint-modulated epithelial-immune communication.

We pretrained EcoNet on pan-cancer immunotherapy cohorts to learn shared response patterns, and fine-tuned it on ccRCC cohorts to align with the disease-specific context (see Methods). We then applied the fine-tuned model to the Leuven cohort, which was kept separate from training. Predicted responder and non-responder groups showed a progression-free survival difference (*P*-value = 0.0240; Figure 5c), demonstrating effective clinical stratification.

As in the pan-cancer model, pathway genes received higher attention than marker genes across the ccRCC ecotypes (Supplementary Figure S29). To study the mechanism of the high-attention genes in shaping the immune-responsive TME, we traced signaling paths among the highest-attention genes in the RE2 network (Figure 5d). The network connected T-cell receptor and co-stimulatory signaling with JAK-STAT cytokine signaling, adhesion, and checkpoint regulation, collectively shaping the RE2 ecosystem.

### Spatial transcriptomic profiling validates the RE2 immune-enriched niche in ccRCC

To validate RE2-associated niches at single-cell spatial resolution, we prepared two tissue microarrays (TMAs) for 46 samples from ccRCC patients treated at Stanford Hospital from 2014 to 2022. From these TMAs, we used CosMx spatial transcriptomics to profile a 1,000-plex gene panel at single-cell resolution across ~450,000 cells (Figure 6a). Subsequently, we annotated eleven cell types and decontaminated the counts with DeSpotX (20) (Supplementary Figure S30, Supplementary Figure S31, and S32), then recovered EcoTyper cell states and ecotypes for the cells (Figure 6b). As a cross-platform validation, epithelial state S01, enriched in adjacent normal samples in TCGA-KIRC dataset (Figure 4b), was significantly more abundant in the eleven adjacent normal samples than in 35 tumor samples (*P*-value = 7.64 1*×*0^*−*7^; Figure 6c), supporting the applicability of the state definitions to the CosMx data.

**Fig. 6.**
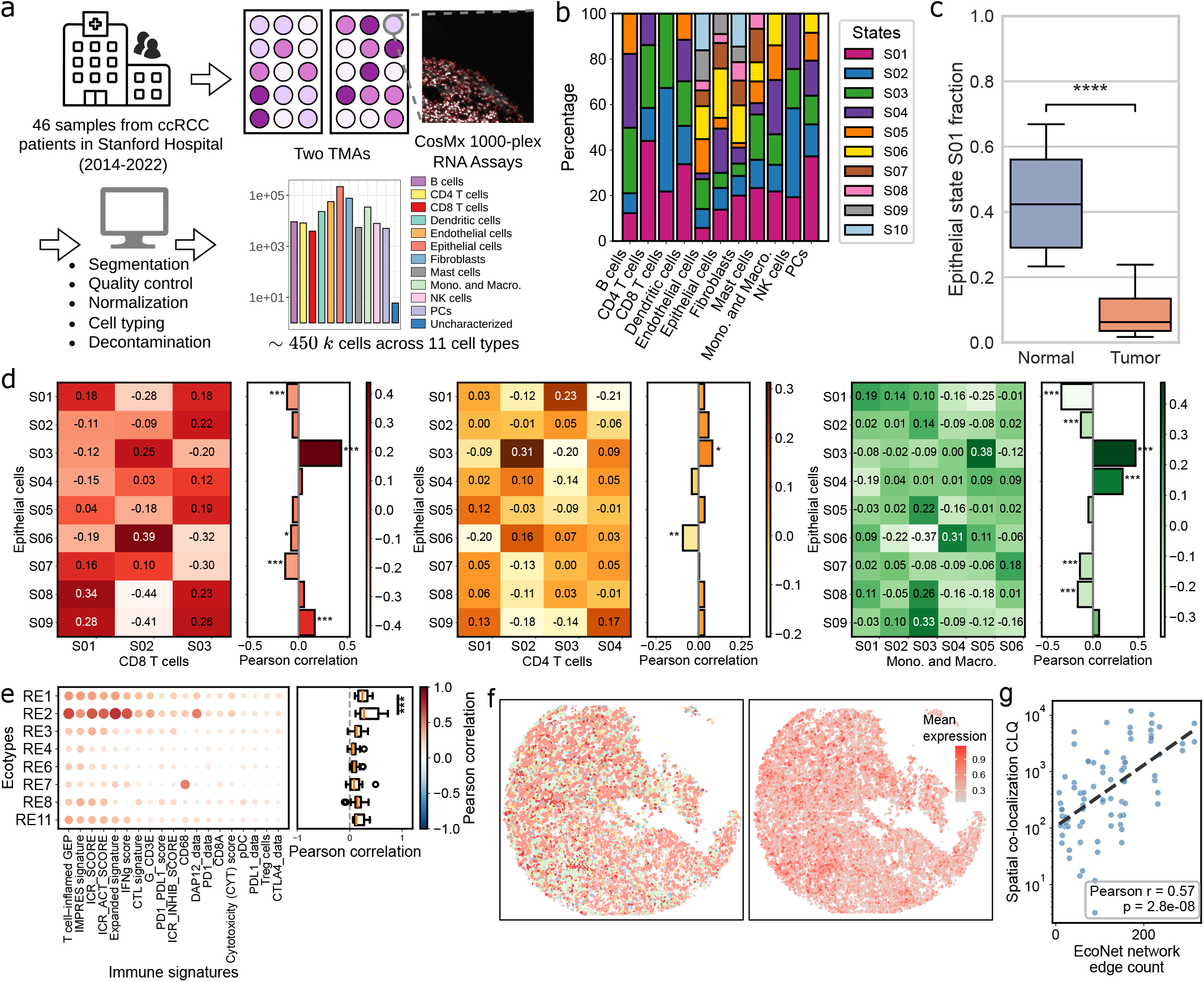
Evaluation of ecotypes and attention scores using ccRCC CosMx data. **a**. Study overview: From 46 ccRCC samples collected from Stanford Hospital, we created two TMAs and applied CosMx 1000-plex profiling. **b**. Cell state composition across the eleven cell types in the CosMx dataset, with stacked bars representing the fraction of the cell states within each cell type. **c**. Distribution of epithelial cell state S1 fractions across normal and tumor samples. **d**. Co-localization of epithelial states with CD8 T cells, CD4 T cells, as well as monocytes and macrophages. Heatmaps show correlations between epithelial and immune cell state fractions within local spatial bins, while barplots show Pearson correlations between epithelial-state fractions and the total fraction of the corresponding immune cell type. **e**. Correlation between ecotype-associated gene expression and established immune signatures. The left heatmap displays Pearson correlations computed on local neighborhoods, with point size indicating the magnitude of correlation and color indicating its direction. The right boxplot summarizes the overall distribution of correlation values for each ecotype across immune signatures. **f**. Spatial distributions of a representative sample, showing cell type localization (left, sharing legend with panel a) and mean expression of RE2 marker genes (right). **g**. Correlation between EcoNet network edge count and spatial co-localization quotient (CLQ). Each point represents a cell-state pair, with edge count defined as the number of ligand-receptor links in the RE2 network and CLQ computed from *k*-nearest-neighbor spatial graphs. Pearson correlation and *P*-value are annotated.

To characterize the spatial immune context of RE2 in tissue, we quantified co-localization between epithelial and immune states. We partitioned the slides into 0.5 mm*×*0.5 mm bins, and within each bin we derived the fraction of each state. We computed the Pearson correlation between per-bin state fractions. Notably, epithelial S03 showed the highest co-localization with CD8 T cells S02, CD4 T cells S02, and monocytes and macrophages S05, consistent with the RE2 composition (Figure 6d). In agreement with this, mean marker expression for epithelial S03 correlated strongly with other RE2 states (Supplementary Figure S33). In addition, we compared epithelial-state fractions to overall immune-cell abundance across bins. S03 exhibited the highest correlation, indicating immune-enriched neighborhoods. We further validated RE2 as an immune-active niche using established immune signatures (2). Specifically, we calculated ecotype expression per spot using the average marker-gene expression, and correlated these with immune signature expression computed as the average of constituent genes; RE2 exhibited the highest Pearson correlation scores across ecotypes (Figure 6e).

We examined a representative sample and mapped the mean expression of RE2 marker genes, which showed contiguous regions of elevated expression within immune-enriched areas (Figure 6f). To determine whether these patterns reflect coherent tissue domains, we applied BANKSY (21) with varying spatial weights. With *λ* = 0.5, recommended for single-cell resolution, we obtained six clusters, one of which showed significantly higher expression of RE2 marker genes (Supplementary Figure S34a). Increasing *λ* to 0.9, which is dominated by spatial proximity, again revealed a cluster with elevated expression (Supplementary Figure S34b). These results indicated that RE2 was enriched within organized spatial domains. Furthermore, the RE2-associated cluster at *λ* = 0.5 was enriched for immune cell types including B cells, CD4 T cells, CD8 T cells, dendritic cells, and NK cells (Supplementary Figure S35a); the enrichment persisted but was reduced at *λ* = 0.9 (Supplementary Figure S35b).

To validate the EcoNet signaling network in spatial data, we computed the co-localization quotient (CLQ) from *k*-nearest-neighbor spatial graphs (*k* = 50) for each cell-state pair and compared it with the number of ligand-receptor links in the RE2 network. Cell-state pairs connected by more edges were significantly more co-localized in tissue (Pearson *r* = 0.57, *P* = 2.8 *×* 10^*−*8^; Figure 6g), supporting that the inferred network structure reflects spatial organization in the TME.

## Discussion

Immunotherapy outcome is shaped by the multicellular organization within the TME, not just the presence of individual populations or interactions. Through ecotype analysis, we resolved this organization and highlighted immune-active communities in which antigen-presenting cells, cytotoxic T cells, and regulatory subsets work together. To further characterize how regulation mediates interplay among these populations, we constructed signaling networks that integrate intracellular regulation and intercellular ligand-receptor communications. Building on these networks, EcoNet encodes bulk expression over the signaling graph through a graph attention network, embedding each sample in an ecotype-informed latent space from which it predicts immunotherapy response. Because the graph encodes known regulatory structure, the attention scores weight genes and edges by their influence on the prediction, revealing the regulatory drivers within the network.

Applying this framework across cancers, we built signaling networks for nine carcinomas and merged them into a pan-cancer network. Each cancer-specific network recovered hub genes consistent with its known biology, while the merged network retained regulatory programs conserved across tumor types, including the immune-active CE9 program enriched for interferon, cytokine, and antigen-presentation pathways. This merged network captures the regulatory programs conserved across cancers, serving as the shared structure for pan-cancer prediction. Pretrained on pan-cancer immunotherapy cohorts, EcoNet outperformed both existing prediction methods and marker-based signatures, indicating that modeling the structured, directional organization of the TME improves response prediction.

ccRCC exhibits disease-specific heterogeneity and is excluded from the pan-cancer ecotype model (9). We therefore built a ccRCC-specific model via a modified EcoTyper pipeline that leverages the pan-cancer cell-type gene sets and infers ccRCC cell states from TCGA-KIRC expression profiles. This model captured ccRCC-specific organization that the pan-cancer model does not represent. ccRCC ecotypes shared marker genes with several pan-cancer ecotypes but did not map directly onto individual pan-cancer ecotypes (Supplementary Figure S36), and in single-cell cohorts the model recovered a higher fraction of predefined states than the pan-cancer model, supporting improved coverage of ccRCC cell-state diversity. Among the resolved ecotypes, we identified RE2 as an immunotherapy-favorable, immune-active community of antigen-presenting cells, cytotoxic T cells, and regulatory subsets. Fine-tuned on ccRCC cohorts, EcoNet transferred its pan-cancer training to this setting and stratified an independent cohort by progression-free survival, showing that the framework adapts to a single disease while retaining the response signal learned across cancers.

Attention scores on the signaling network quantify each gene’s contribution to propagating regulation along edges, indicating gene-level importance within the ecotype-informed signaling network. In both the CE9 and RE2 networks, attention concentrated on intermediate pathway genes rather than ecotype markers, indicating that the scores capture the signaling genes that connect state-defining programs. Since each gene’s score is computed from the same fixed number of edges, this difference is not attributable to pathway genes having higher degree. In the pan-cancer CE9 network, high-attention genes were enriched among those required for T-cell-mediated tumor killing in an independent CRISPR screen, linking them to the mechanism underlying immunotherapy response. In the ccRCC RE2 network, the highest-attention genes mapped to pathways connecting T-cell activation, immune infiltration, and epithelial responses, with checkpoint and cytokine-JAK-STAT pathways among the candidate programs that shape the RE2 ecosystem.

Beyond immunotherapy response prediction, the EcoNet framework is applicable to other downstream tasks. The graph attention layers of EcoNet generate embeddings that integrate gene expression, signaling pathways, and ecotype structure into unified representations. These embeddings summarize the higher-order organization of cellular programs and can serve as transferable features for downstream analyses. Serving as the inputs of supervised or unsupervised models, they can be applied to tasks such as prognosis modeling (22, 23), treatment stratification (24, 25), and subtype discovery (26).

## Methods

### Identification of cell states and ecotypes

For the pan-cancer analyses, we used the cell states and carcinoma ecotypes (CE1-CE10) from the published pan-cancer EcoTyper model (9). For ccRCC, which is excluded from the pan-cancer model, we derived a disease-specific set of cell states and ecotypes from TCGA-KIRC as follows.

We downloaded bulk RNA-seq data for 614 clear cell renal cell carcinoma (ccRCC) samples from the TCGA-KIRC cohort (accessed in July 2024). The expression profiles were provided in transcripts per million (TPM) and organized into a non-negative matrix *X* ∈ℝ^*G×N*^, where *G* denotes the number of genes and *N* the number of tumor samples.

We then applied EcoTyper on the TCGA-KIRC samples to identify cell states within each cell type and group co-occurring states across samples into *ecotypes* (9). In the original implementation, EcoTyper estimates cell-type-specific expression profiles via *in silico* deconvolution using CIBERSORTx. However, the limited number of TCGA-KIRC samples and the high transcriptomic complexity of the TME led to unreliable estimates. To ensure robust state inference, we modified the workflow to operate directly on cell-type-specific subsets of *X*. Specifically, for each of the 12 cell types, we defined a curated gene set *G*_*c*_ from the pan-cancer EcoTyper reference (9). The corresponding cell-type-specific expression submatrix was denoted as 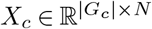, containing expression values of genes *g* ∈*G*_*c*_ across all samples.

For each *X*_*c*_, EcoTyper applied non-negative matrix factorization (NMF) to decompose the expression profiles into *k*_*c*_ latent transcriptional states as

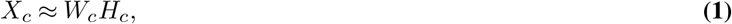

where 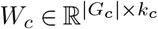 encodes state-specific gene signatures, and 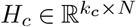 represents the abundance of each state across samples. Low-quality states were filtered based on marker-gene enrichment, following the standard EcoTyper criteria. Across all cell types, pairwise co-abundance of state activities was quantified to form a cross-state similarity matrix *C ∈* ℝ^*S×S*^, where 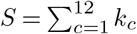 represents the total number of identified cell states. Hierarchical clustering of *C* grouped co-occurring states into eleven ecotypes, each representing a distinct cell-state composition characteristic of the ccRCC TME.

For each sample *i* and ecotype *e*, the ecotype abundance *A*_*e,i*_ was computed as the mean abundance of its constituent cell states 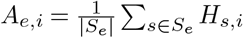, where *S*_*e*_ denotes the set of cell states assigned to ecotype *e*. Consequently, we obtained an ecotype-by-sample abundance matrix *A* ∈ℝ^*E×N*^, where *E* = 11, providing a quantitative representation of the TME composition across ccRCC samples. In addition, ecotype assignment was defined as *e*_*i*_ = argmax_*e*_ *A*_*e,i*_, summarizing the dominant TME configuration for each sample.

### Recovery of cell states and ecotypes across multi-omics datasets

We projected cell states and ecotypes onto independent bulk, single-cell, and spatial transcriptomic datasets using the EcoTyper framework. For the pan-cancer analysis, we recovered the published pan-cancer states and ecotypes in each of the nine carcinomas (BLCA, BRCA, CRC, ESCA, HNSC, LUAD, LUSC, PDAC, and STAD) from its TCGA bulk RNA-seq and scRNA-seq data (Supplementary Table S5). For ccRCC, we recovered the ccRCC-specific states and ecotypes on the Leuven bulk cohort and on independent single-cell and spatial datasets.

For a given bulk RNA-seq cohort, represented by 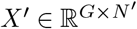, we fixed the state-specific gene signature matrices *W*_*c*_ and optimized the state abundance matrix with

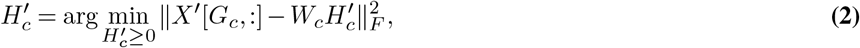

for each cell type *c*. The resulting 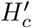 matrices were then combined across cell types to calculate ecotype abundances as described above.

For scRNA-seq datasets, given a cell type *c*, the expression vector of each cell 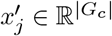 was projected onto the reference weight matrix *W*_*c*_ by solving

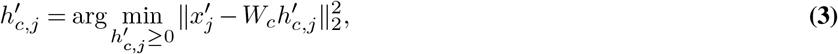

where 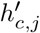 denotes the inferred state abundance profile for cell *j*. Each cell was then assigned to the state with maximal abundance, and cell-level state abundances were subsequently aggregated within each sample to infer sample-level ecotype composition.

For 10x Visium spatial transcriptomic data, the spot-level expression matrix was decomposed against the reference state signature matrices following the bulk-recovery procedure to estimate state and ecotype abundances per spot. CosMx spatial transcriptomic data were analyzed analogously to the single-cell datasets, where each cell was projected onto the reference state signatures corresponding to its annotated cell type and subsequently mapped to its spatial coordinates.

### Construction of intercellular and intracellular signaling networks

#### Identification of ligand-receptor interactions between cell states

We recovered cell states and ecotypes from each cancer’s scRNA-seq data (Supplementary Table S5), which served as the reference to infer ligand-receptor (LR) interactions between cell states and characterize intercellular communication.

For each scRNA-seq sample, we inferred interactions between cell states based on probabilistic modeling of ligand-receptor expression, as described in CellChat (27). Specifically, we first applied Wilcoxon rank-sum tests to identify genes overexpressed within each state, defined by adjusted *p <* 0.05 and positive *log*_2_ fold-change. We then mapped the overexpressed genes to the curated NicheNet LR database (28) and retained LR pairs with at least one overexpressed partner as candidate interactions. For each candidate LR pair (*l, r*), We estimated the communication probability between sender state *i* and receiver state *j* using a Hill-type function

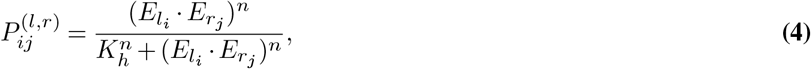

where 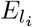 and 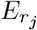 denote the mean expression of the ligand *l* and receptor *r* in states *i* and *j*, respectively. The Hill constant *K*_*h*_ was set to 0.5, and the Hill coefficient *n* to 1. Using 100 bootstrap permutations of cell-state labels, we identified significant ligand-receptor interactions with *p <* 0.05.

#### Selection of ligands activating downstream marker genes

Although ligand-receptor interactions define potential communication routes, they do not reveal which ligands activate transcriptional programs that define receiver cell states. We therefore assessed ligand activity following the NicheNet framework (28), evaluating each ligand’s ability to predict the expression of receiver-state marker genes. For each sender-receiver pair, we selected candidate ligands from the significant LR interactions and defined receiver-state marker genes as downstream targets. Ligand-target prior probabilities were retrieved from NicheNet, which links ligands to putative gene targets. We quantified ligand activity by comparing predicted target profiles with the observed marker set using the area under the precision-recall curve (AUPR). The top five ligands with the highest AUPR scores per sender-receiver pair were incorporated into subsequent network construction.

Building on the identified ligand-target relationships, we constructed ecotype-specific signaling networks by linking active ligands to downstream marker genes through curated ligand-receptor and gene-regulatory interactions from NicheNet. The resulting directed, weighted network represents signaling and transcriptional pathways within each ecotype.

#### Integration of pathway genes into signaling networks

To enable reconstruction of continuous signaling routes from ligands to transcription factors, we integrated intermediate pathway genes into the signaling networks.

For each ligand-target pair, we enumerated signaling paths of up to two regulatory links and calculated a cumulative probability for each path

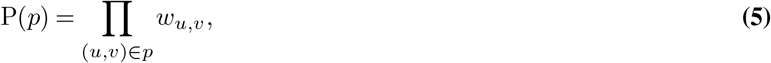

where *w*_*u,v*_ denotes the interaction weight between genes *u* and *v*. We then scored the intermediate genes by the summed probability of all traversing paths,

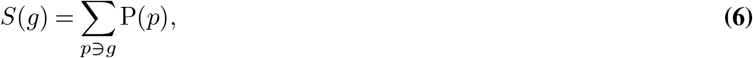

reflecting their contributions to signal transmission within the ligand-target cascade. For each ligand, we retained the top two pathway genes with the highest scores, and the resulting subgraphs containing ligands, pathway genes, and downstream marker genes were merged to form ecotype-specific signaling networks.

#### Construction of cancer-specific and pan-cancer networks

We merged the ecotype-specific networks of each of the nine carcinomas into a cancer-specific network representing regulation within that cancer’s TME, then combined the nine into a pancancer network. To capture regulation shared across tumor types, we retained edges present in at least five of the nine cancers, along with the genes they connect. We set each edge weight to its mean per-cancer weight, scaled by its cross-cancer frequency. This yielded a pan-cancer network of 1,956 nodes and 12,219 edges.

For ccRCC, we integrated the networks from the eleven ecotypes into a global signaling network, characterizing the intercellular and intracellular regulatory relationships across the ccRCC TME.

### Graph-based modeling of TME for immunotherapy response prediction

#### Representation learning from signaling graphs with graph attention network

We represented the TME as a directed graph *G* = (*V, ℰ*) derived from the integrated intercellular and intracellular signaling network described above, using the pan-cancer network for the pan-cancer model and the ccRCC network for the ccRCC model. Each node *v*_*i*_ *∈ V* corresponds to a gene, and each edge (*v*_*i*_, *v*_*j*_) *∈ ℰ* encodes a regulatory interaction with weight *w*_*ij*_. For each sample, we used the *z*-score-normalized bulk RNA-seq expression profile **x** ∈ℝ^|*V*|^ as node features (Supplementary Method S1.1 and Supplementary Figure S37), so that the samples share the same graph topology but differ in their expression attributes.

To capture structured gene-gene dependencies that define ecotype composition, we implemented an ecotype-guided representation learning framework based on a graph attention network. The model comprises two attention layers that iteratively update node embeddings by weighting neighbor contributions according to learned gene-gene relevance. For each layer *l*, the model computes the attention weight 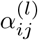 between connected nodes *i* and *j* as

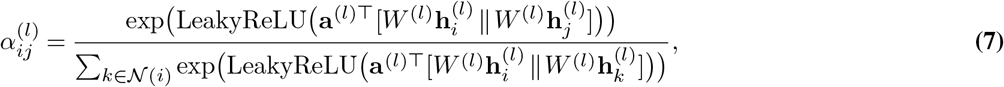

where *N*(*i*) denotes the neighbors of node *i, W* ^(*l*)^ is the learnable linear transformation, and **a**^(*l*)^ is the learnable attention kernel. The initial node representation is given by 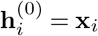, and node embeddings are then updated as

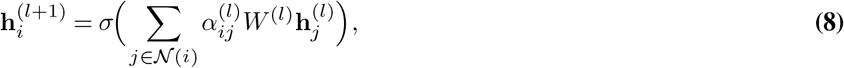

where *σ* denotes the exponential linear unit (ELU) activation function. The first layer uses multi-head attention with four independent heads concatenated to enrich representation diversity, while the second layer aggregates one head into a unified latent space.

We concatenated all node embeddings into a single graph-level representation and passed it through two fully connected layers to obtain an ecotype-aligned latent vector **z** *∈*ℝ^*E*^, where *E* is the number of ecotypes (*E* = 10 for the pan-cancer model and *E* = 11 for ccRCC). The first fully connected layer transforms the flattened embedding of |*V*| nodes into a 128-dimensional latent space with rectified linear unit (ReLU) activation, and the second maps this representation to the ecotype space with softmax normalization. The model was trained to minimize the Kullback-Leibler divergence between the predicted representation and the EcoTyper distribution **a**.

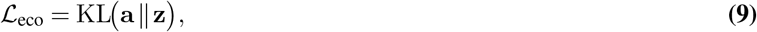

where **a**∈ℝ^*E*^ denotes the ecotype distribution from the EcoTyper-derived abundance matrix *A*. This objective function guides the model to capture transcriptomic variation consistent with EcoTyper-defined TME organization, whereas the learned attention coefficients highlight gene-gene interactions that contribute most to ecotype differentiation.

We trained the graph representation module on 6,498 TCGA samples from the nine carcinomas for the pan-cancer model, and on the 614 TCGA-KIRC samples for the ccRCC model, using Adam optimization, with a learning rate of 5 *×*10^*−*4^ and weight decay of 2*×*10^*−*4^. We applied dropout with a rate of 0.2 to both attention and fully-connected layers to mitigate overfitting, and adopted an early stopping strategy to determine convergence within 1,000 epochs. A detailed summary of model hyperparameters is provided in Supplementary Table S6. To confirm that the module learns stable graph embeddings, we performed five-fold cross-validation within the training set (Supplementary Figure S38).

#### Pretraining on pan-cancer immunotherapy datasets

To provide sufficient training data and enable the response prediction module to capture generalizable transcriptomic patterns associated with immunotherapy outcomes, we pretrained both the pancancer and ccRCC models on our recently curated pan-cancer immunotherapy dataset (29), each on its own signaling graph.

From the resource, we selected the treatment-naive primary samples with matched bulk expression profiles and clinical response annotations (*n* = 2, 532) for model pretraining. We computed gene-wise *z*-scores within each dataset separately, ensuring that normalization did not use information from other cohorts (Supplementary Method S1.1 and Supplementary Figure S37) and imputed the missing values using a *k*-nearest neighbor approach (Supplementary Method S1.2). For each sample, the normalized expression vector was projected onto the corresponding signaling graph and embedded through the trained graph representation module, yielding an ecotype representation **z** ∈ℝ^*E*^ (*E* = 10 for the pan-cancer model and *E* = 11 for the ccRCC model) that captures the transcriptomic state in the learned ecotype space.

We implemented the response prediction module as a feed-forward network *f*_*θ*_ : ℝ^*E*^ *→*ℝ^2^ that maps the ecotype representation **z** to a probability vector **ŷ** = *f*_*θ*_(**z**) for responder and non-responder outcomes. The classifier comprised two fully connected layers with 32 and 16 hidden units, each followed by a rectified linear activation. Batch normalization was applied after the first layer to stabilize optimization. The final softmax layer generated a responder and non-responder probability vector, denoted as **ŷ**.

We optimized the model parameters by minimizing a weighted cross-entropy loss,

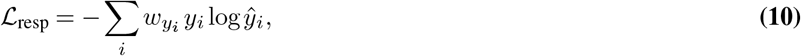

where *y*_*i*_ denotes the one-hot-encoded true label for sample *i*, and *ŷi* represents the predicted probability for each response class. The class weights *wy*_*i*_ were inversely proportional to class frequencies across cohorts to correct for the imbalance between responder and non-responder samples. We performed the optimization using the Adam optimizer with a learning rate of 10^*−*3^, a batch size of 16, and a dropout rate of 0.1. Model performance was monitored by early stopping based on area under the receiver operating characteristic curve (AUC) scores to prevent overfitting. We summarized the model hyperparameters in Supplementary Table S7. The pretraining step enabled the network to capture transcriptomic features associated with immunotherapy response. The pretrained pan-cancer model was used directly for prediction, whereas the pretrained ccRCC model was further fine-tuned on ccRCC cohorts as described below.

#### Fine-tuning on cancer-specific immunotherapy cohorts

We fine-tuned the pretrained ccRCC EcoNet model on ccRCC immunotherapy cohorts (*n* = 365) to adapt it to ccRCC-specific molecular and immune features. The fine-tuning process jointly optimized the graph representation and response prediction modules over *T* = 20 epochs by iteratively minimizing the two objective functions (Algorithm 1). Hyperparameter details are provided in Supplementary Table S8.

##### Algorithm 1

Iterative fine-tuning of EcoNet on ccRCC immunotherapy cohorts

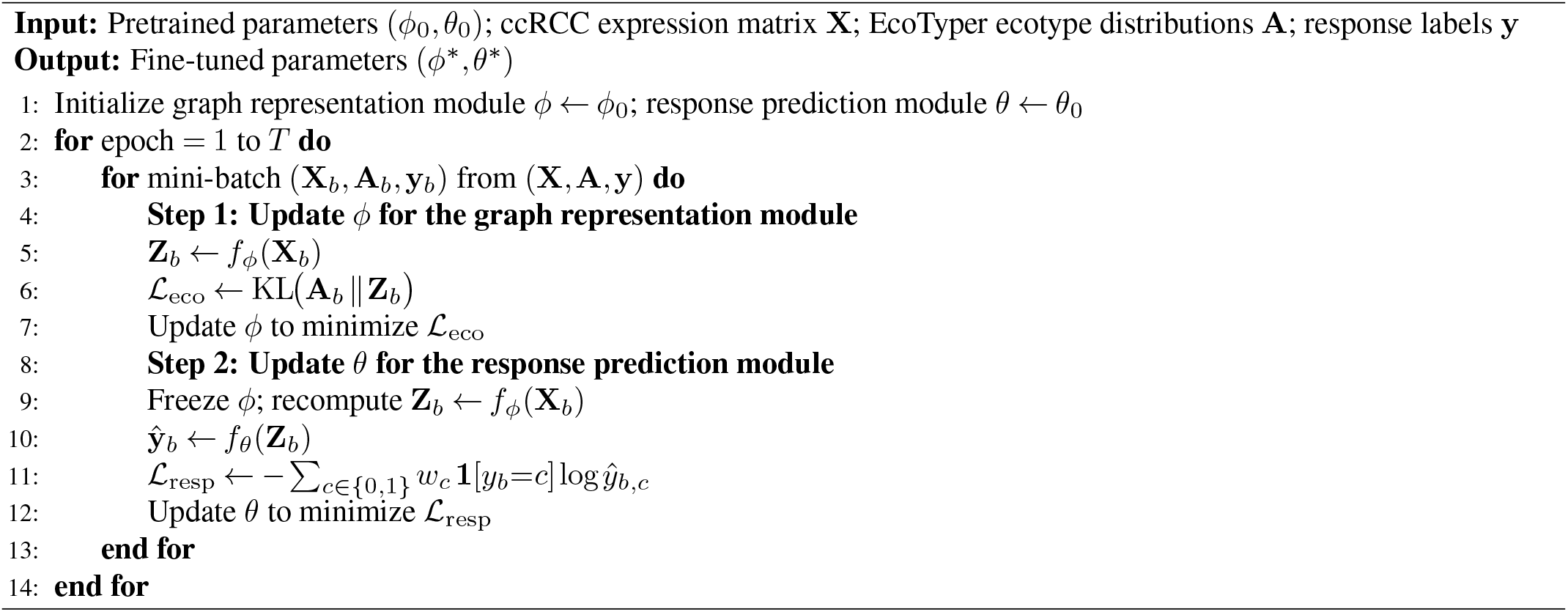

After fine-tuning, we extracted edge-level attention coefficients *α*_*ij*_ from the graph representation module and averaged them across all patients to derive global attention weights for gene-gene interactions

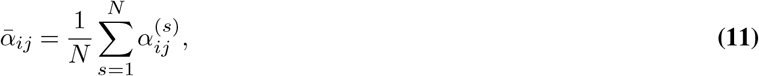

where *N* denotes the number of ccRCC samples. For each gene *i*, we computed an aggregate attention score as the sum of its top *k* (*k* = 5) highest-attention edges,

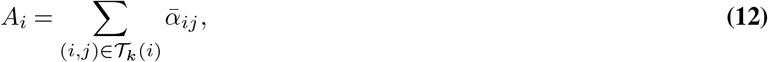

quantifying its overall regulatory influence within the ccRCC signaling network. By restricting the sum to a fixed number of edges (*k* = 5), the attention score reflects the strength of a gene’s strongest connections rather than the degree, thereby avoiding a bias toward highly connected genes. Genes with high attention scores were identified as key regulators shaping the transcriptional and immune organization of the ccRCC TME.

#### Benchmark methods for immunotherapy response prediction

We compared EcoNet with existing prediction methods and with marker-based baselines under a leave-one-cohort-out strategy over the 43 immunotherapy cohorts, holding out each cohort in turn. Evaluation was restricted to the 24 cohorts containing both responders and non-responders. ccRCC cohorts were excluded from the pan-cancer benchmark and analyzed with the ccRCC-specific EcoNet model.

The prediction methods included TIDE (5), ImmuCellAI (30), EaSIeR (31), NetBio (32), COMPASS (33) and the immune landscape signature (ILS) (2). For these methods, we followed the provided tutorials and applied the default parameter settings. TIDE, ImmuCellAI, EaSIeR, NetBio, and COMPASS predict response from bulk expression profiles, for which we used TPM data as their inputs. For TIDE, we additionally log-transformed the expression and centered each gene to its cohort mean. For COMPASS, we used its provided leave-one-cohort-out models for cohorts within its training set. For ILS, which uses precomputed immune-signature scores, we grouped samples into ILS-high versus ILS-low by Ward-linkage hierarchical clustering and designated ILS-high as responders (Supplementary Method S1.3). We restricted this comparison to the 12 melanoma and NSCLC cohorts where all methods were applicable, since the response prediction component of TIDE was established on these cancers (5).

The marker-based baselines included single-gene markers (*PDCD1, CD274, CD8A* and *CTLA4*), the combined checkpoint marker GeneBio, and immune signature scores, adopted from COMPASS (33) (Supplementary Table S9). For each marker, we z-scored the values across samples and grouped the samples by Ward-linkage hierarchical clustering into two groups, then designated the higher-scoring group as responders (Supplementary Method S1.4). These baselines were compared across all 24 cohorts.

#### TMA preparation and CosMx spatial transcriptomic profiling

We constructed two ccRCC tissue microarrays (TMAs) from formalin-fixed paraffin-embedded (FFPE) tissue blocks of samples collected at Stanford Hospital between 2014 and 2022. The TMAs were assembled using a manual tissue arrayer with 1 mm cores and 0.5 mm spacing; each sample was represented in triplicate, and a placenta core was included at the end of the grid for orientation. In total, 35 tumor and 11 adjacent normal samples were represented across two TMAs.

The two TMA blocks were sectioned at 5 *μ*m thickness and mounted on LEICA Bond Plus slides. Tissue preparation followed the CosMx SMI Manual Slide Preparation for RNA Assays (MAN-10184-04), using default conditions for target retrieval time, digestion buffer concentration, digestion time, and fiducial concentration. Sections were hybridized overnight with the CosMx Universal Cell Characterization 1,000-plex RNA probe panel and stained the following day with CosMx DAPI nuclear stain, CosMx Hs CD298/B2M, and CosMx Hs PanCK/CD45 morphology markers.

#### CosMx data processing and analysis

We processed CosMx TMA data following the NanoString-recommended pipeline via a Seurat workflow (34). We excluded cells with total RNA counts *<* 20 and fields of view with *<* 100 cells. Then we normalized gene counts using SCTransform, which modeled counts by regularized negative binomial regression and applied residual clipping to mitigate depth effects and regularize gene-wise mean-variance properties.

For cell typing, we first constructed a ccRCC-specific signature matrix from a scRNA-seq dataset (35) using CIBERSORTx (36) with default parameters, restricting genes to those on the CosMx panel (Supplementary Figure S39). We then applied Insitutype supervised clustering (37) with the reference. We incorporated negative-control probe intensities to model background noise and used 10 random starts with a maximum of 30 iterations to stabilize convergence. Insitutype returned cell types and posterior probabilities for each cell, and we retained cells with the posterior assignment probability *>* 0.8. Starting from this annotation, we decontaminated the CosMx counts with DeSpotX (20), iterating decontamination and re-annotation three times to remove ambient contamination.

## Data availability

This study was approved by the Stanford University Institutional Review Board (Protocol #68299). Test data for EcoNet have been deposited in Zenodo at https://doi.org/10.5281/zenodo.22986445. The CosMx data generated in this study have been deposited in the Gene Expression Omnibus (GEO) and will be publicly available upon publication.

## Code availability

The source code for EcoNet was deposited to GitHub https://github.com/Gentles-lab/EcoNet.

## Acknowledgements

This work was supported by the National Institutes of Health NCI Cancer Systems Biology Consortium under award number U01CA264611.

## Author contributions

R.H.W. and A.J.G conceived the idea. A.J.G and B.H supervised the project. R.H.W. developed the methods and performed the experiments. B.A.B collected pan-cancer immunotherapy samples. R.A.H, S.N, and B.H. generated the CosMx data. R.H.W. and I.I. processed the CosMx data. R.H.W. wrote the manuscript. B.A.B., I.I., S.Z., R.A.H, S.N., B.H., and A.J.G revised the manuscript.

