## Supplemental Materials for "Deep-learning-driven tumor microenvironment profiling improves immunotherapy response prediction"

October 6, 2026

**S3 Supplementary Tables** **6**

**S4 Supplementary Figures** **14**

### S1 Supplementary Methods

#### S1.1 Cohort-wise $z$-score normalization of bulk RNA-seq data

To harmonize gene expression profiles across multiple cohorts, we normalized bulk RNA-seq data to gene-wise $z$-scores within each dataset. This step ensured that expression values were placed on a comparable scale across studies while preserving the relative expression structure within each cohort. We applied the same normalization approach to both training and evaluation datasets to maintain consistent feature scaling.

For each cohort $c$ and gene $g$, we computed the mean and standard deviation of expression across all samples belonging to that dataset. The normalized expression for sample $i$ in cohort $c$ was defined as

$$z_{g,i} = \frac{x_{g,i} - \mu_{g,c}}{\sigma_{g,c}},$$

where $x_{g,i}$ denotes the raw expression of gene $g$ in sample $i$, and $\mu_{g,c}$ and $\sigma_{g,c}$ represent the mean and standard deviation of that gene within dataset $c$. The resulting standardized expression values were used as node feature input for EcoNet graph-based modeling. We present the UMAP plots of pan-cancer immunotherapy cohorts before and after $z$-score normalization; the results indicate enhanced cross-cohort comparability following normalization (Supplementary Figure S37).

#### S1.2 Imputation of missing expression values

To ensure feature consistency across datasets and enable cross-cohort comparability, we used the TCGA-KIRC cohort as the reference expression space and imputed missing expression values after gene-wise $z$-score normalization. Missing values were estimated using a $k$-nearest neighbor (KNN) approach implemented in the `scikit-learn` package (v1.5), which infers unobserved values from the most similar samples in the high-dimensional expression space. This approach preserves the intrinsic gene co-expression structure while enabling consistent representation across cohorts.

For a gene $g$ and a sample $i$ with missing expression $x_{g,i}$, the imputed value $\hat{x}_{g,i}$ was computed as the average expression of $g$ among its $k$ nearest neighbors with observed values:

$$\hat{x}_{g,i} = \frac{1}{k} \sum_{j \in \mathcal{N}_k(i)} x_{g,j},$$

where $\mathcal{N}_k(i)$ denotes the index set of the $k$ nearest samples ($k = 16$) of $i$ in the gene expression space. We then used the imputed expressions as model inputs for subsequent EcoNet training and evaluation.

#### S1.3 Immune Landscape Signature benchmark

Following [1], we began with 115 immune landscape gene signatures and selected candidate benefit-associated signatures based on pan-cancer immunotherapy analyses. For each cohort, we first retained signatures with $\geq 90\%$ of the gene set covered by the cohort's RNA-seq profiling and computed signature scores for each sample as the mean expression of genes. To select benefit-associated signatures, we applied two filters: (i) PFS association via univariable Cox models, retaining signatures with HR $< 0.9$ and p $< 0.25$; and (ii) ORR separation using two-sample tests (equal-variance t-test, Welch's t-test, or Mann-Whitney, chosen based on normality and variance), retaining signatures with p $< 0.25$ and higher mean in responders than non-responders. The union of PFS- and ORR-selected sets defined a benefit-associated ILS panel. For the signatures in the panel, we $z$-standardized the scores across samples, then clustered samples on these $z$-scored features using Ward's hierarchical clustering ($k = 2$), designating the higher-scoring cluster as ILS-high and the other as ILS-low.

#### S1.4 Gene signature benchmark

For the PD1, PDL1, and CD8A signatures, we $z$-standardized gene expression values across samples and clustered samples using Ward's hierarchical clustering ($k = 2$). The cluster with the higher mean expression was labeled as the responder group, and the other the non-responder.

### S2 Supplementary Results

#### S2.1 Ablation analysis of EcoNet components

To quantify the contribution of each component of EcoNet, we compared the full model against five ablation variants, evaluated under the leave-one-cohort-out strategy. (1) *Without GAT* replaces GAT-derived ecotype abundances with raw EcoTyper abundances, keeping the response predictor unchanged. (2) *Without ecotype* replaces ecotype-supervised pre-training with end-to-end training on response labels. (3) *Without edges* removes all graph edges and uses a feed-forward network to map gene expression to ecotype abundances. (4) *Without learned head* removes the trained response predictor and instead classifies a sample as a responder when its most abundant predicted ecotype is CE9 ($\text{argmax}_e\, z_e = \text{CE9}$). (5) *Without z-normalization* trains the graph model on raw TPM values rather than per-cohort gene-wise $z$-scored expression, testing whether input normalization is necessary for cross-cohort generalization.

Removing any component reduced performance (Supplementary Figure S2). The full EcoNet model achieved a median AUC of 69.9% across the 12 cohorts, compared with 66.8% without the GAT, 64.0% without ecotype supervision, 62.7% without edges, 56.4% without the learned head, and 50.0% without $z$-normalization. Without $z$-normalization, the model trained on raw TPM collapsed to a constant output (AUC = 50.0% in every cohort), reflecting the cross-cohort batch effects shown in Supplementary Figure S37.

#### S2.2 Benchmarking three EcoTyper models in ccRCC

We compared the cell-level recovery rate in ccRCC single-cell data across three models, the ccRCC EcoTyper model used in this study, the pan-cancer EcoTyper model [2], and a ccRCC model built with a conventional workflow. The conventional model used CIBERSORTx [3] to deconvolute bulk expression into cell-type-specific expression profiles and then applied non-negative matrix factorization within each cell type to define states.

We collected ccRCC single-cell datasets from diverse cohorts to test recovery robustness (Supplementary Table S2). From the merged pool, we randomly sampled 100,000 cells to limit computational cost. Of these, 90,855 cells were annotated to eleven of the twelve cell types in the EcoTyper models. On this set, the ccRCC EcoTyper model recovered cell states for 54.2% of the cells; the ccRCC model built with the conventional workflow recovered 48.6%; the pan-cancer model recovered 46.2% (Supplementary Table S3). In summary, the ccRCC EcoTyper model achieves the highest recovery rate. The pan-cancer model was broadly applicable across cancer types but showed lower recovery on ccRCC. This aligns with prior findings that ccRCC is not well profiled by pan-cancer frameworks and requires a tumor-specific model [2].

#### S2.3 Functional enrichment of epithelial state markers

We performed Gene Ontology (GO) enrichment analysis on epithelial state markers using GSEApy [4], and the results highlighted programs of cell migration and adhesion (Supplementary Figure S7a). The analysis also recovered renal absorption terms from the normal-enriched state S01, indicating that the model captures kidney-specific biology. Excluding S01 left migration, adhesion, and immune-response programs dominant (Supplementary Figure S7b).

#### S2.4 Enrichment of ccRCC epithelial markers in KIRC

We quantified epithelial-state marker expression across TCGA cancer types by computing the standardized mean for each cancer type. As expected, KIRC showed the highest expression of ccRCC model epithelial state markers, followed by KIRP (papillary renal cell carcinoma); while KICH (chromophobe renal cell carcinoma) expressed few of them (Supplementary Figure S8). This observation indicates that across the three kidney cancer cohorts, KIRC and KIRP share similar expression profiles, whereas KICH is distinct, consistent with previous reports [5].

#### S2.5 Significant recovery rate of ccRCC EcoTyper cell states in single-cell data

We next evaluated the recovery rate at the state level in the collected ccRCC single-cell data. For each cell type, EcoTyper computes a recovery $Z$-score for each predefined cell state; states with $Z > 1.96$ (one-sided $p < 0.05$) were considered significantly recovered. We summarize performance as the state-level significant recovery rate, defined as the number of significantly recovered states divided by the total number of states per lineage and overall. Across cell types, the ccRCC EcoTyper model significantly recovered 57 of 70 states (81.4%) (Supplementary Table S4 and Supplementary Figure S9). The 13 unrecovered states were distributed across seven cell types.

#### S2.6 Enriched epithelial-immune communication in RE2

To examine the roles of cell-state interactions in this association, we leveraged a ccRCC scRNA-seq dataset by mapping ecotype cell states to it and then inferring inter-cellular crosstalk [6]. Among eight samples, one mapped to RE2, two to RE4, one to RE8, three to RE10, and one was unassigned (Supplementary Figure S15). For each sample we inferred state-to-state intercellular communications with CellChat [7]. As representative cases, sample P915 (assigned to RE2) displayed prominent epithelial-immune edges with higher weights, whereas sample P913 (assigned to RE4) showed little epithelial-immune interactions (Supplementary Figure S16a; additional samples in Supplementary Figure S17). We summarized pathway-level communication probabilities for P915, with strong signals in interferon and chemokine signaling, complement and antigen presentation, and extracellular matrix and adhesion pathways concentrated on epithelial-immune interactions (Supplementary Figure S16b).

### S3 Supplementary Tables

Supplementary Table S1: Evaluation cohorts used for benchmarking.

Independent immunotherapy cohorts used to benchmark EcoNet under a leave-one-cohort-out strategy. Marker scores were evaluated across all cohorts, whereas prediction tools were evaluated on the melanoma and NSCLC
cohorts to maintain consistency with TIDE, excluding cohorts on which ImmuCellAI did not run successfully.

| **Cohort** | **Samples (n)** | **Cancer type** | **Immunotherapy** |
|---|---|---|---|
| GSE123728 [8] | 13 | Melanoma | anti-PD1 |
| Gide_Cell_2019 [9] | 47 | Melanoma | anti-PD1 & anti-CTLA4 |
| GSE165278 [10] | 22 | Melanoma | anti-CTLA4 |
| GSE122220 | 10 | Melanoma | NA |
| CheckMate_064 [11] | 42 | Melanoma | anti-PD1 & anti-CTLA4 |
| Liu_NatMed_2019 [12] | 83 | Melanoma | anti-PD1 |
| phs000452 [13] | 153 | Melanoma | anti-CTLA4 |
| PUCH [14] | 55 | Melanoma | anti-PD1 |
| Nathanson_2017 [15] | 64 | Melanoma | anti-CTLA4 |
| GSE136961 [16] | 21 | Non-small cell lung cancer | anti-PD1 |
| VanAllen_antiCTLA4_2015 [17] | 41 | Melanoma | anti-CTLA4 |
| HugoLo_IPRES_2016 [18] | 17 | Melanoma | anti-PD1 |
| IMVigor210 [19] | 182 | Urothelial (bladder) | anti-PD-L1 (atezolizumab) |
| GSE111636 | 11 | Urothelial (bladder) | anti-PD-1/PD-L1 |
| Kim_NatMed_2018 [20] | 45 | Gastric | anti-PD1 (pembrolizumab) |
| Zhao_NatMed_2019 [21] | 25 | Glioblastoma | anti-PD1 (pembrolizumab) |
| iSPY2_2022 [22] | 69 | Breast | neoadjuvant anti-PD1 + chemo |
| GSE173839 [23] | 105 | Breast | durvalumab + olaparib (I-SPY2) |
| GSE159067 [24] | 102 | Head & neck | anti-PD1 |
| GSE140901 [25] | 24 | Hepatocellular (liver) | anti-PD1 |
| GSE181815 [26] | 10 | Thymic epithelial | anti-PD1 (pembrolizumab) |
| GSE165252 [27] | 71 | Esophageal | anti-PD-L1 (atezolizumab) |
| GSE183924 [28] | 37 | Esophagogastric | anti-PD1 |
| GSE252990 [29] | 40 | Mesothelioma | anti-PD1 ± anti-CTLA4 |

Supplementary Table S2: Single-cell ccRCC cohorts used in benchmarking.

| **BioProject** | **Series** | **Accession Number** | **SampleName** |
|---|---|---|---|
| PRJNA923244 | GSE222703 | SRS16435936 | GSM6929206 |
| PRJNA931987 | GSE224630 | NA | GSM7028034 |
| PRJNA931987 | GSE224630 | NA | GSM7028035 |
| PRJNA931987 | GSE224630 | NA | GSM7028036 |
| PRJNA931987 | GSE224630 | NA | GSM7028037 |
| PRJNA931987 | GSE224630 | NA | GSM7028039 |
| PRJNA855923 | GSE207493 | SRS13709498 | GSM6290520 |
| PRJNA855923 | GSE207493 | SRS13709500 | GSM6290522 |
| PRJNA855923 | GSE207493 | SRS13709502 | GSM6290524 |
| PRJNA855923 | GSE207493 | SRS13709504 | GSM6290544 |
| PRJNA855923 | GSE207493 | SRS13709506 | GSM6290546 |
| PRJNA855923 | GSE207493 | SRS13709508 | GSM6290548 |
| PRJNA855923 | GSE207493 | SRS13709510 | GSM6290550 |
| PRJNA855923 | GSE207493 | SRS13709512 | GSM6290526 |
| PRJNA855923 | GSE207493 | SRS13709514 | GSM6290518 |
| PRJNA855923 | GSE207493 | SRS13709515 | GSM6290528 |
| PRJNA855923 | GSE207493 | SRS13709518 | GSM6290552 |
| PRJNA855923 | GSE207493 | SRS13709520 | GSM6290554 |
| PRJNA855923 | GSE207493 | SRS13709522 | GSM6290530 |
| PRJNA855923 | GSE207493 | SRS13709524 | GSM6290532 |
| PRJNA855923 | GSE207493 | SRS13709526 | GSM6290534 |
| PRJNA855923 | GSE207493 | SRS13709528 | GSM6290536 |
| PRJNA855923 | GSE207493 | SRS13709530 | GSM6290538 |
| PRJNA855923 | GSE207493 | SRS13709532 | GSM6290542 |
| PRJNA855923 | GSE207493 | SRS13709533 | GSM6290540 |
| PRJNA436710 | GSE111360 | NA | KIRC_GSE111360 |
| PRJNA497905 | GSE121636 | NA | KIRC_GSE121636 |
| PRJNA667716 | GSE159115 | NA | KIRC_GSE159115 |
| PRJNA641020 | GSE152938 | SRS6885097 | GSM4630028 |
| PRJNA641020 | GSE152938 | SRS6885098 | GSM4630029 |

Supplementary Table S3: Comparison of cell state recovery across three EcoTyper models on ccRCC single-cell data.

| **Cell type** | **Total number (n)** | **Recovered by ccRCC EcoTyper** | **Recovered by pan-cancer EcoTyper** | **Recovered by ccRCC conventional** |
|---|---|---|---|---|
| B cells | 943 | 450 | 460 | 548 |
| CD4 T cells | 2,909 | 1,314 | 1,345 | 1,509 |
| CD8 T cells | 10,489 | 5,533 | 1,828 | 5,408 |
| Dendritic cells | 384 | 217 | 117 | 168 |
| Endothelial cells | 8,396 | 4,789 | 3,905 | 3,570 |
| Epithelial cells | 30,017 | 15,415 | 15,203 | 14,618 |
| Fibroblasts | 6,285 | 3,421 | 3,291 | 2,861 |
| Mast cells | 637 | 282 | 345 | 330 |
| Monocytes and Macrophages | 18,527 | 9,432 | 10,155 | 7,645 |
| NK cells | 11,545 | 8,178 | 5,079 | 7,117 |
| Plasma cells | 723 | 246 | 247 | 417 |
| **All cells** | **90,855** | **49,277** | **41,975** | **44,191** |

Supplementary Table S4: State-level significant recovery rate in single-cell data using the ccRCC EcoTyper model. States with $Z > 1.96$ (one-sided $p < 0.05$) were considered significantly recovered.

| **Cell type** | **States (n)** | **Significantly recovered (n)** |
|---|---:|---:|
| B cells | 5 | 4 |
| CD4 T cells | 4 | 3 |
| CD8 T cells | 3 | 3 |
| Dendritic cells | 5 | 3 |
| Endothelial cells | 10 | 9 |
| Epithelial cells | 9 | 9 |
| Fibroblasts | 10 | 7 |
| Mast cells | 8 | 4 |
| Monocytes and Macrophages | 6 | 6 |
| NK cells | 4 | 4 |
| Plasma cells | 6 | 5 |
| **Total** | **70** | **57** |

Supplementary Table S5: Single-cell RNA-seq datasets used to construct the per-cancer regulatory networks.

| **Cancer** | **Accession** | **Samples** | **Cells** |
|---|---|---|---|
| BLCA | SRR12603780-790 | 11 | 28,358 |
| BRCA | GSE176078 | 26 | 43,402 |
| CRC | GSE132465 | 23 | 22,404 |
| HNSC | GSE164690 | 18 | 29,002 |
| LUAD | GSE131907 | 15 | 22,712 |
| LUSC | GSE148071 | 16 | 49,905 |
| PDAC | GSE155698 | 15 | 18,211 |
| ESCA | GSE160269 | 93 | 208,658 |
| STAD | GSE167297 | 5 | 22,464 |
| ccRCC | [6] | 8 | 14,733 |

Supplementary Table S6: Hyperparameters of the EcoNet graph representation module.

The hyperparameters were selected via grid search using five-fold cross-validation.

| Component | Hyperparameter | Symbol / Setting | Description |
| --- | --- | --- | --- |
| **Input graph** | Topology | $\mathcal{G} = (\mathcal{V}, \mathcal{E})$ | Integrated intercellular and intracellular signaling network |
| | Node features | $\mathbf{x}_i \in \mathbb{R}^{\|\mathcal{V}\|}$ | $z$-score-normalized bulk RNA-seq expression values per gene |
| | Edge weights | $w_{ij}$ | Encoded regulatory or signaling strengths from prior network integration |
| **Graph attention layer 1** | Input dimension | 1 | Normalized expression |
| | Output dimension | 8 × 4 heads | Multi-head attention with concatenated outputs |
| | Activation function | ELU | Applied to attention-aggregated features |
| | Dropout rate | 0.2 | Applied to attention coefficients and node embeddings |
| **Graph attention layer 2** | Input dimension | 32 | Aggregated multi-head outputs from the previous layer |
| | Output dimension | 8 | Single-head aggregation producing latent node representations |
| | Activation function | ELU | Nonlinear activation following aggregation |
| | Dropout rate | 0.2 | Applied to node embeddings |
| **Fully connected layer 1** | Input dimension | $\|\mathcal{V}\| \times 8$ | Flattened node embeddings |
| | Output dimension | 128 units | Projects graph representation to latent feature space |
| | Activation function | ReLU | Nonlinear transformation of latent representation |
| | Dropout rate | 0.2 | Applied before the next layer |
| **Fully connected layer 2** | Input dimension | 128 | Latent representation from the previous layer |
| | Output dimension | 11 | Ecotype representation dimension |
| | Activation function | Softmax | Normalizes output to probability space |
| **Loss and optimization** | Objective function | $\mathcal{L}_{\text{eco}} = \text{KL}(\mathbf{a} \,\\|\, \mathbf{z})$ | KL divergence between predicted and EcoTyper-derived ecotype distributions |
| | Optimizer | Adam | Adaptive gradient optimization |
| | Learning rate | $5 \times 10^{-4}$ | Optimization step size |
| | Weight decay | $2 \times 10^{-4}$ | L2 regularization strength |
| | Batch size | 8 | Number of samples per iteration |
| | Training epochs | $\leq 1000$ | Monitored using early stopping |

Supplementary Table S7: Hyperparameters of the EcoNet response prediction module pretrained on pan-cancer immunotherapy datasets.

| Component | Hyperparameter | Symbol / Setting | Description |
|---|---|---|---|
| **Input features** | Input dimension | $E = 11$ | Ecotype representation vector derived from the graph representation module |
| **Fully-connected layer 1** | Output dimension | 32 | First transformation of input |
| | Activation function | ReLU | Nonlinear transformation of hidden representation |
| | Batch normalization | Applied | Stabilizes feature distributions |
| **Fully-connected layer 2** | Output dimension | 16 | Second transformation of latent features |
| | Activation function | ReLU | Nonlinear activation of transformed space |
| | Dropout rate | 0.1 | Regularization to reduce overfitting |
| **Output layer** | Output dimension | 2 | Binary classification: responder vs. non-responder |
| | Activation function | Softmax | Converts logits to normalized class probabilities |
| **Loss and optimization** | Objective function | $\mathcal{L}_{\text{resp}} = -\sum_i w_{y_i} y_i \log \hat{y}_i$ | Weighted cross-entropy correcting for class imbalance |
| | Class weighting | $w_{y_i} \propto 1/n_{y_i}$ | Inverse-frequency weighting by class size |
| | Optimizer | Adam | Adaptive gradient optimization |
| | Learning rate | $10^{-3}$ | Step size for parameter updates |
| | Weight decay | $1 \times 10^{-4}$ | L2 regularization strength |
| | Batch size | 16 | Number of samples per iteration |
| | Training epochs | $\leq 50$ | Monitored using early stopping |

Supplementary Table S8: Hyperparameters of EcoNet fine-tuning on ccRCC immunotherapy cohorts.

| Component | Hyperparameter | Symbol / Setting | Description |
|---|---|---|---|
| **Graph representation module ($\phi$)** | Objective function | $\mathcal{L}_{\mathrm{eco}}$ | Kullback-Leibler divergence between predicted and reference ecotype distributions |
| | Optimizer | Adam | Adaptive gradient optimization |
| | Learning rate | $1 \times 10^{-4}$ | Step size for parameter updates |
| | Weight decay | $1 \times 10^{-8}$ | L2 regularization term |
| | Batch size | 8 | Samples per training iteration |
| **Response prediction module ($\theta$)** | Objective function | $\mathcal{L}_{\mathrm{resp}}$ | Weighted cross-entropy loss |
| | Class weighting | $w_c \propto 1/n_c$ | Inverse-frequency weighting by response category |
| | Optimizer | Adam | Adaptive gradient optimization |
| | Learning rate | $1 \times 10^{-8}$ | Fine-tuning step size |
| | Batch size | 8 | Synchronized with graph module updates |
| **Training procedure** | Epochs | 20 | Maximum fine-tuning iterations with early stopping |
| | Regularization | Dropout = 0.2 | Applied to both attention and dense layers |

Supplementary Table S9: Marker-based baselines used for benchmarking.

Details of the single-gene markers and immune signatures used for benchmarking; the immune signatures are adopted from COMPASS [30].

| **Marker** | **Full name** | **Type** | **Genes (n)** |
|---|---|---|---|
| PD1 | Programmed cell death protein 1 (PDCD1) | Gene marker | 1 |
| PDL1 | Programmed death-ligand 1 (CD274) | Gene marker | 1 |
| CD8A | CD8a molecule | Gene marker | 1 |
| CTLA4 | Cytotoxic T-lymphocyte-associated protein 4 | Gene marker | 1 |
| GeneBio | Combination of PD-1, PD-L1 and CTLA-4 | Signature | 3 |
| CKS | Chemokine signature score | Signature | 12 |
| GEP | T cell-inflamed gene expression profile | Signature | 18 |
| IMPRES | Immuno-predictive score | Signature | 15 |
| IFNG | Interferon-$\gamma$ signature score | Signature | 18 |
| Teff | T-effector interferon-$\gamma$ signature | Signature | 8 |
| NRS | Neoadjuvant response signature | Signature | 69 |
| CIS | Cytotoxic immune signature | Signature | 7 |
| CTL | Cytotoxic T lymphocyte markers | Signature | 5 |
| TAM | Tumor-associated macrophages | Signature | 50 |
| Texh | T cell exhaustion | Signature | 50 |
| IS | Roh immune score | Signature | 41 |
| ICA | Immune cytolytic activity score | Signature | 2 |
| MIAS | MHC I association immune score | Signature | 100 |
| PGM | Paired gene markers | Signature | 2 |
| CAF | Cancer-associated fibroblast signature score | Signature | 5 |

### S4 Supplementary Figures

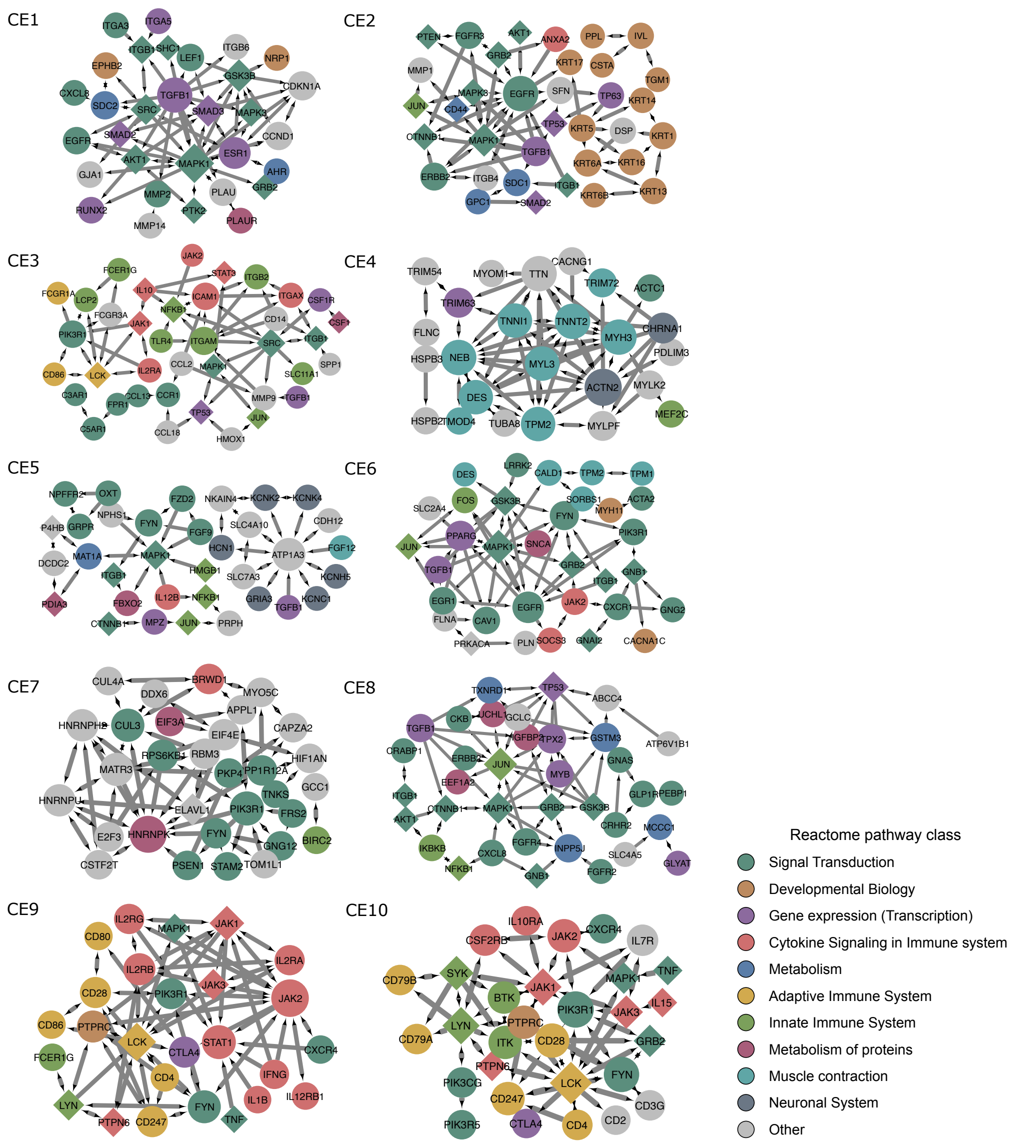

Supplementary Figure S1: **Signaling subnetworks of the ten carcinoma ecotypes (CE1-CE10).** Each network shows the top 40 genes by degree within the ecotype in the pan-cancer network. Node size scales with degree, node color indicates the Reactome functional class, and node shape distinguishes marker genes (circles) from pathway genes (diamonds). Directed edges indicate signaling direction.

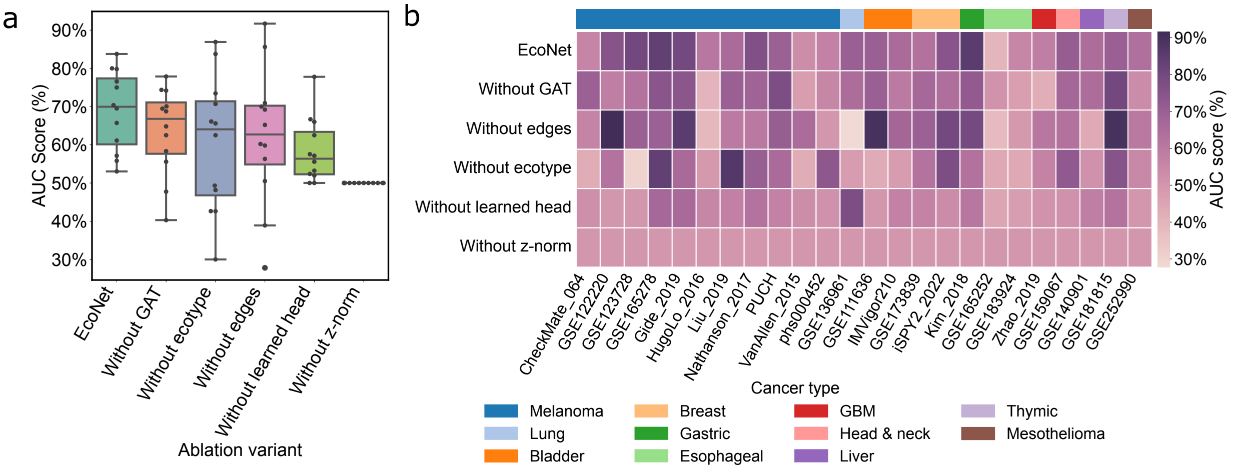

Supplementary Figure S2: **Ablation analysis of EcoNet components.** **a**. Comparison of EcoNet with its four ablated variants, ordered by median AUC score. Boxplots summarize the distribution of AUC scores across the test datasets applicable to all variants. **b**. Comparison of EcoNet with its four ablated variants, ordered by median AUC score. The heatmap shows AUC scores across the test datasets, with cancer type annotated.

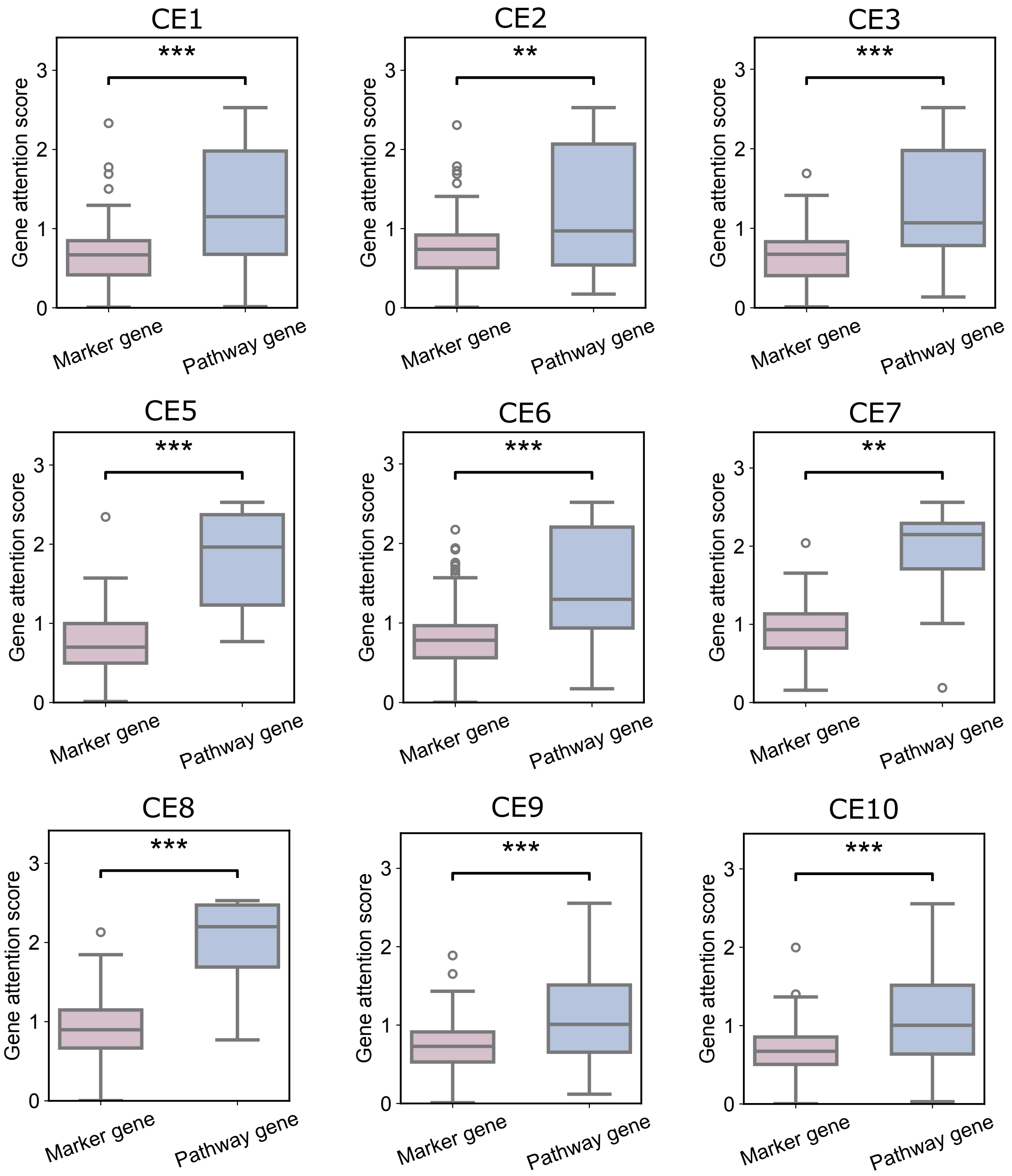

Supplementary Figure S3: **Comparison of attention scores between ecotype marker genes and pathway genes for each pan-cancer ecotype network.** CE4 network was excluded, which lacked pathway genes.

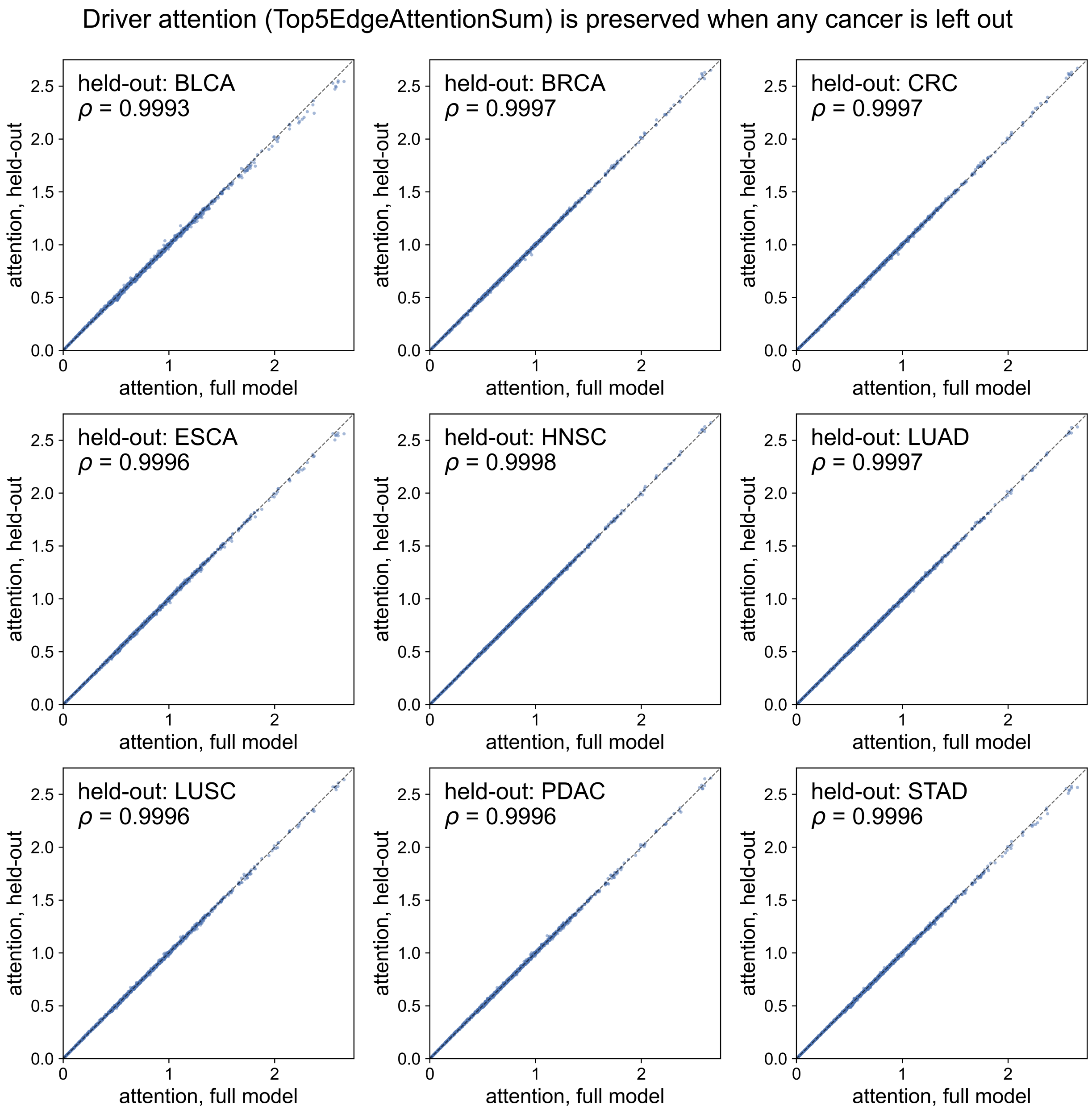

Supplementary Figure S4: **Robustness of gene attention scores to training-set composition.** Each panel compares per-gene attention scores from the full model with those from a model trained with one cancer type held out. Spearman correlations ($\rho$) are annotated in each panel. The dashed line indicates the identity line.

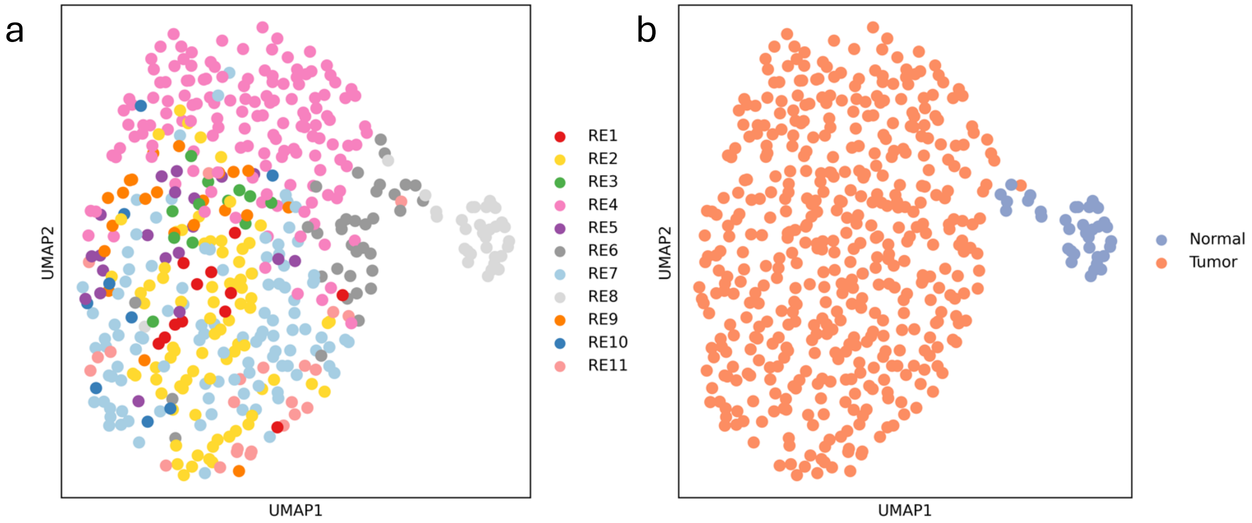

Supplementary Figure S5: **UMAP projections of TCGA-KIRC samples. a.** UMAP of TCGA-KIRC expression profiles, with samples colored by assigned ecotype. **b.** UMAP of the same projection, with samples colored by clinical sample type.

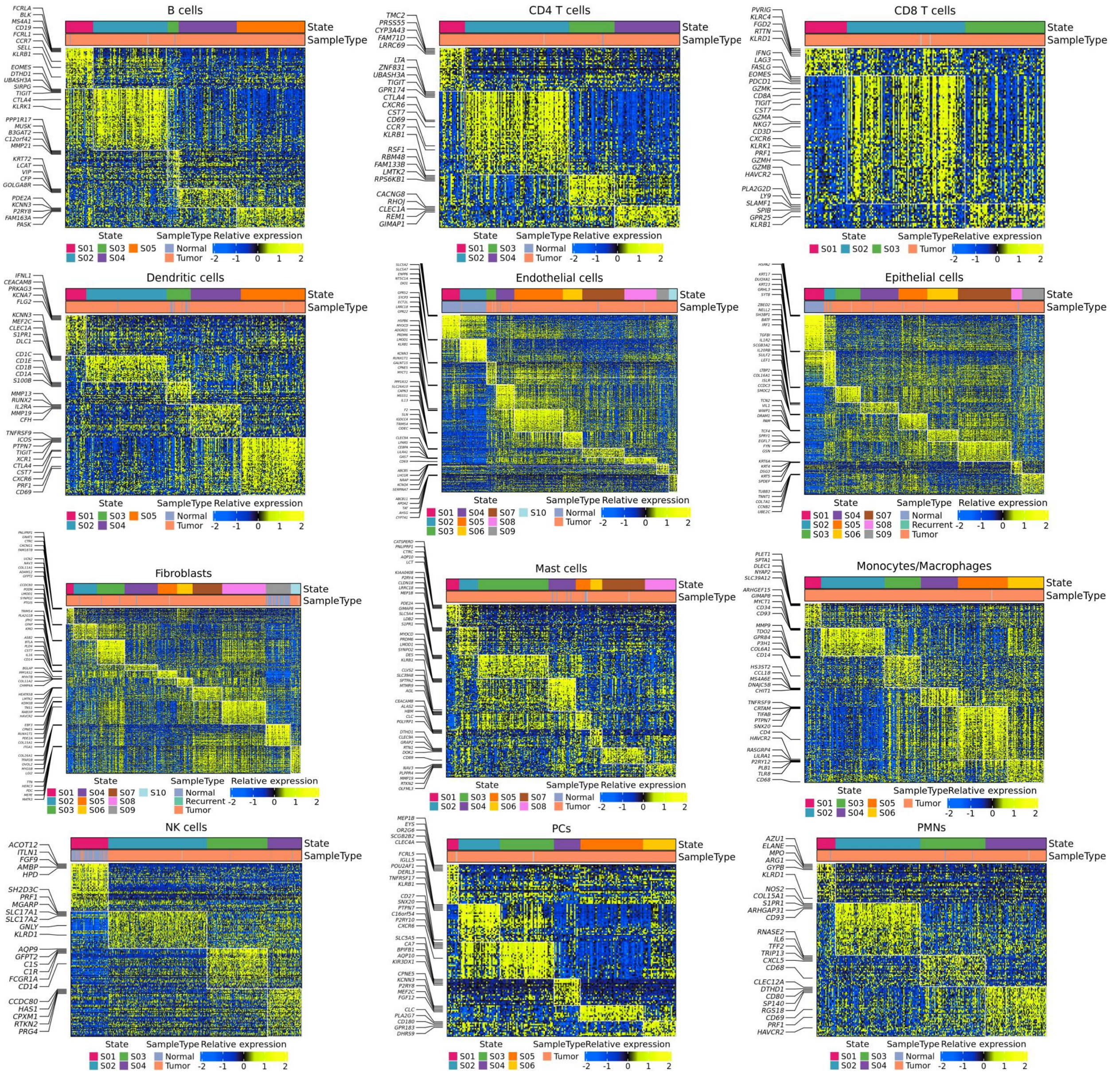

Supplementary Figure S6: **Heatmaps of cell states across TCGA-KIRC samples by cell type.** Each row represents a marker gene and each column a sample, with color intensity reflecting relative expression. The top bar indicates epithelial state assignments, and five representative marker genes for each state are highlighted on the left.

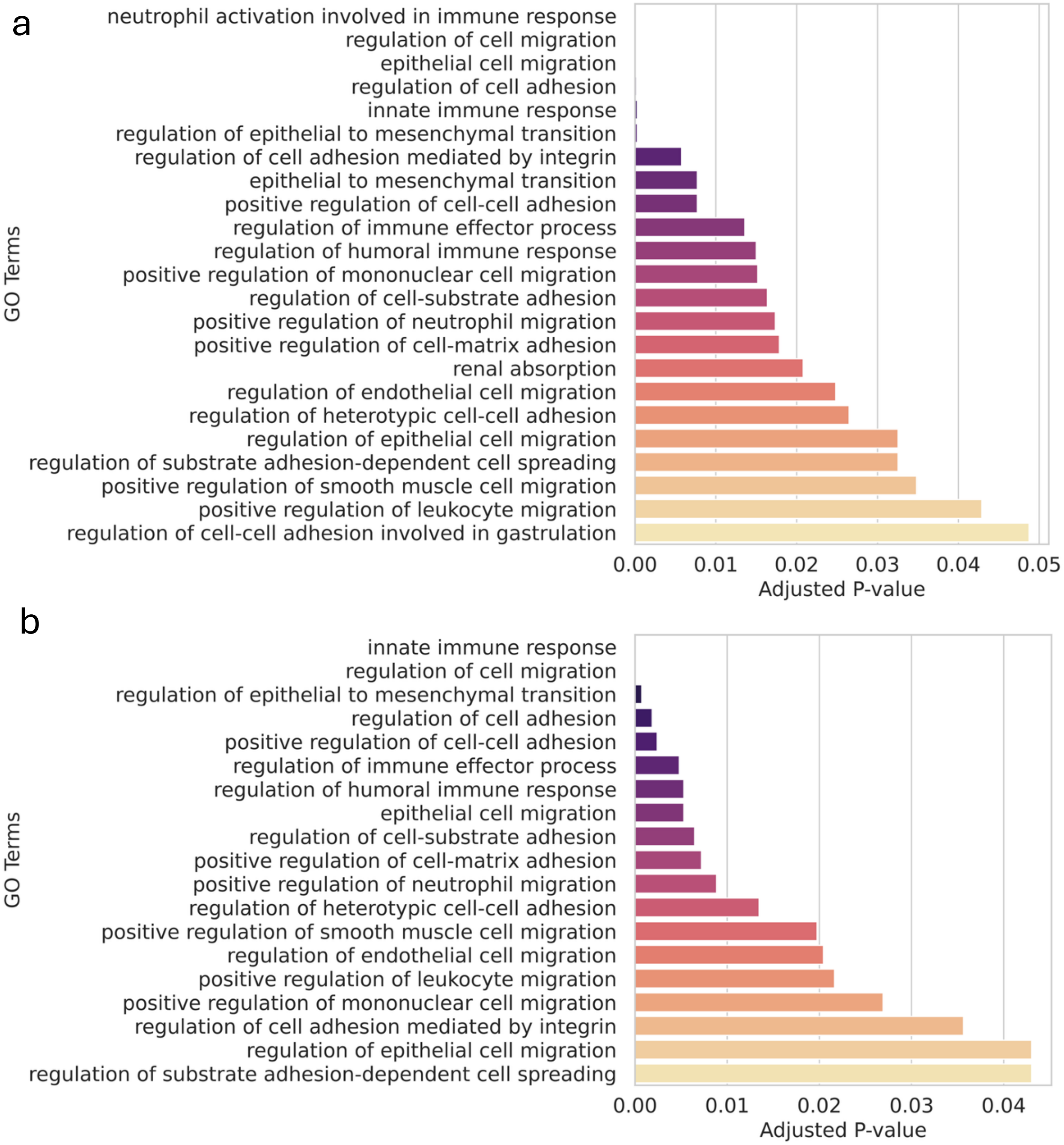

Supplementary Figure S7: **Functional enrichment of epithelial cell state marker genes. a.** The barplot shows selected GO terms enriched among marker genes with adjusted $P$-values$< 0.05$. **b.** The same analysis after excluding state S01.

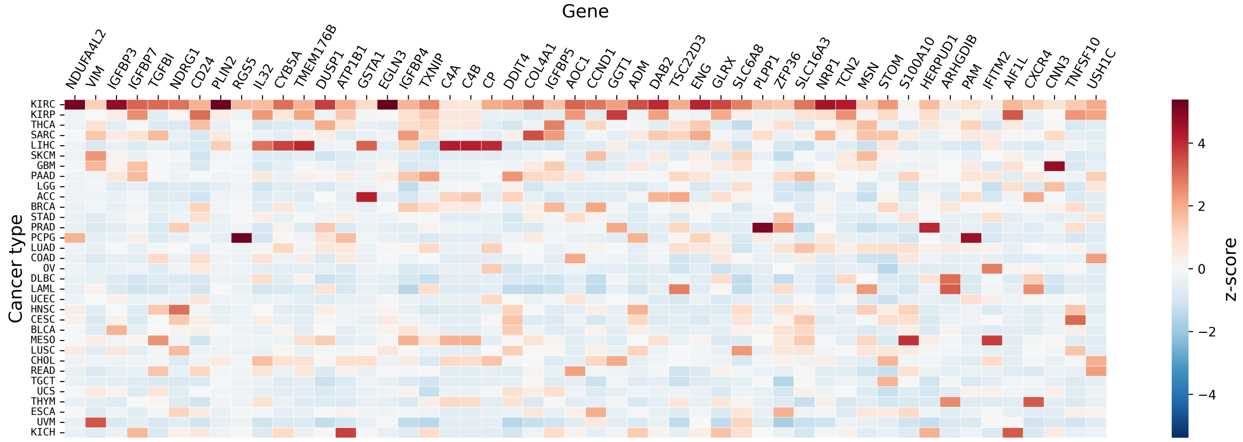

Supplementary Figure S8: **Heatmap showing the standardized mean expression of epithelial state marker genes across multiple cancer types.** Cancer types are ordered by median standardized expression, and genes are ordered by differential expression between KIRC and other cancer types, with the top 50 retained.

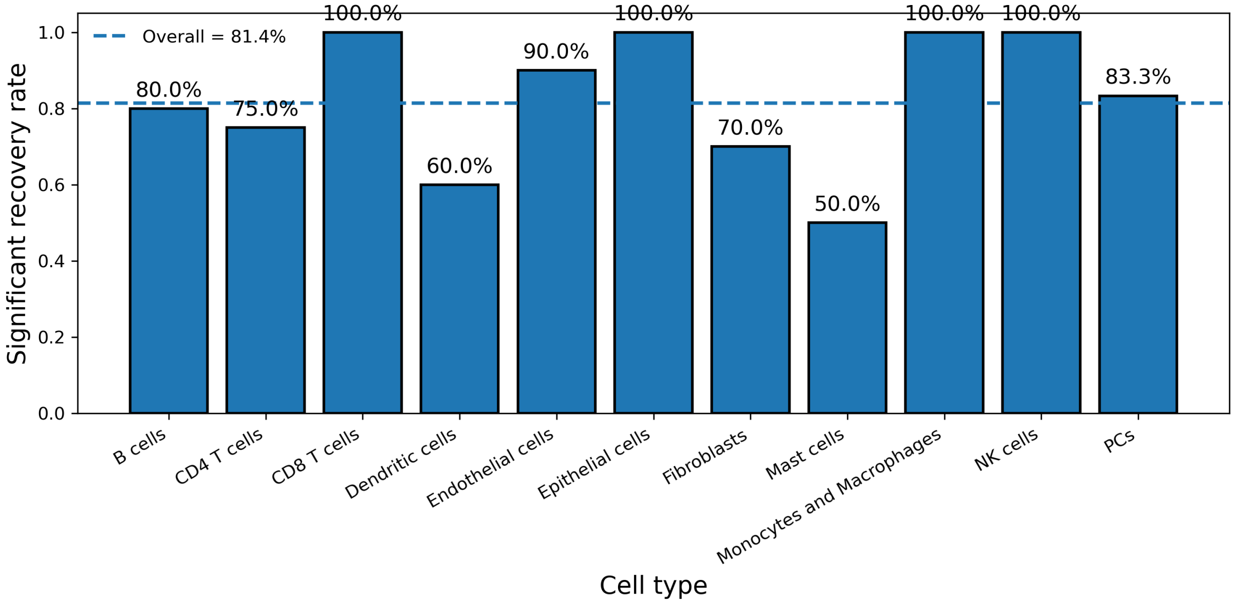

Supplementary Figure S9: **State-level significant recovery rate in single-cell data using the ccRCC EcoTyper model.** Bars show the fraction of predefined states significantly recovered per cell type. The dashed line indicates the overall recovery rate.

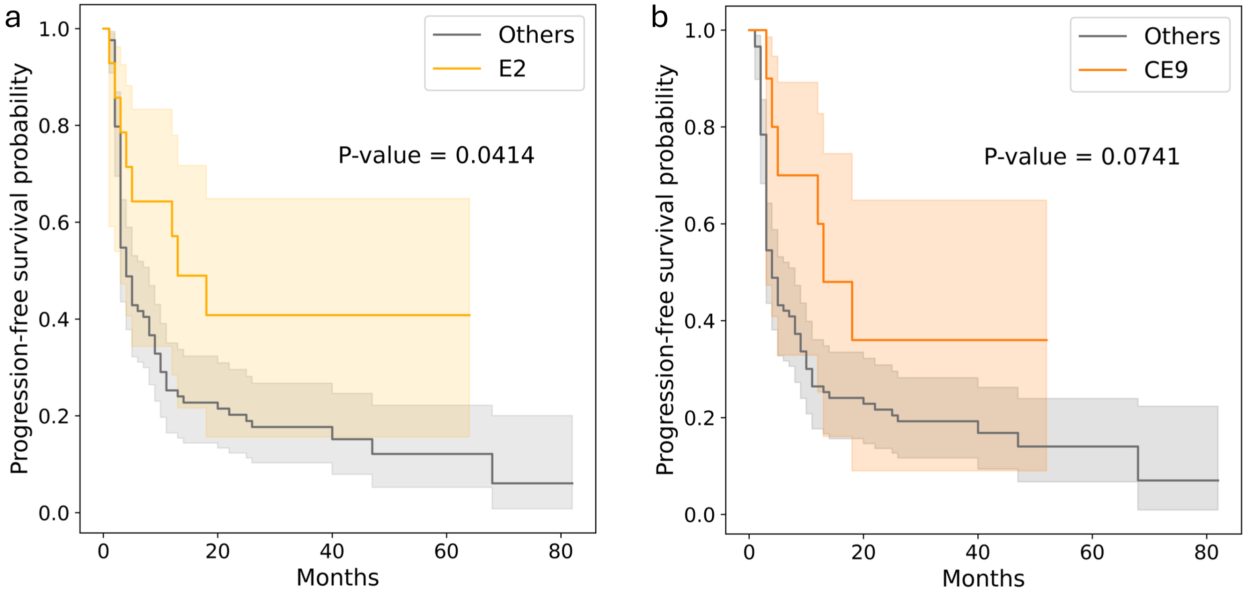

Supplementary Figure S10: **Comparison between RE2 and CE9 stratifications for progression-free survival. a.** Progression-free survival analysis comparing patients assigned to RE2 versus all other ecotypes in our ccRCC EcoTyper model, with $P$-value annotated. **b.** Progression-free survival analysis comparing patients assigned to CE9 versus all other ecotypes in the pan-cancer EcoTyper model, with $P$-value annotated.

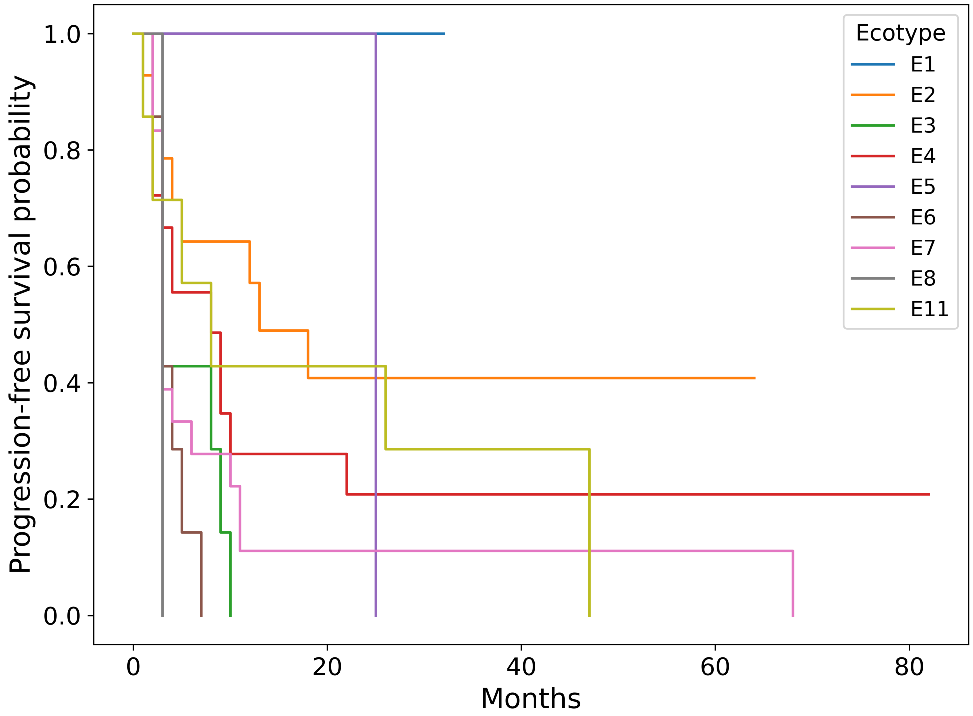

Supplementary Figure S11: **Progression-free survival stratified by ecotype.** Each curve denotes the Kaplan-Meier estimate of remaining progression-free over follow-up.

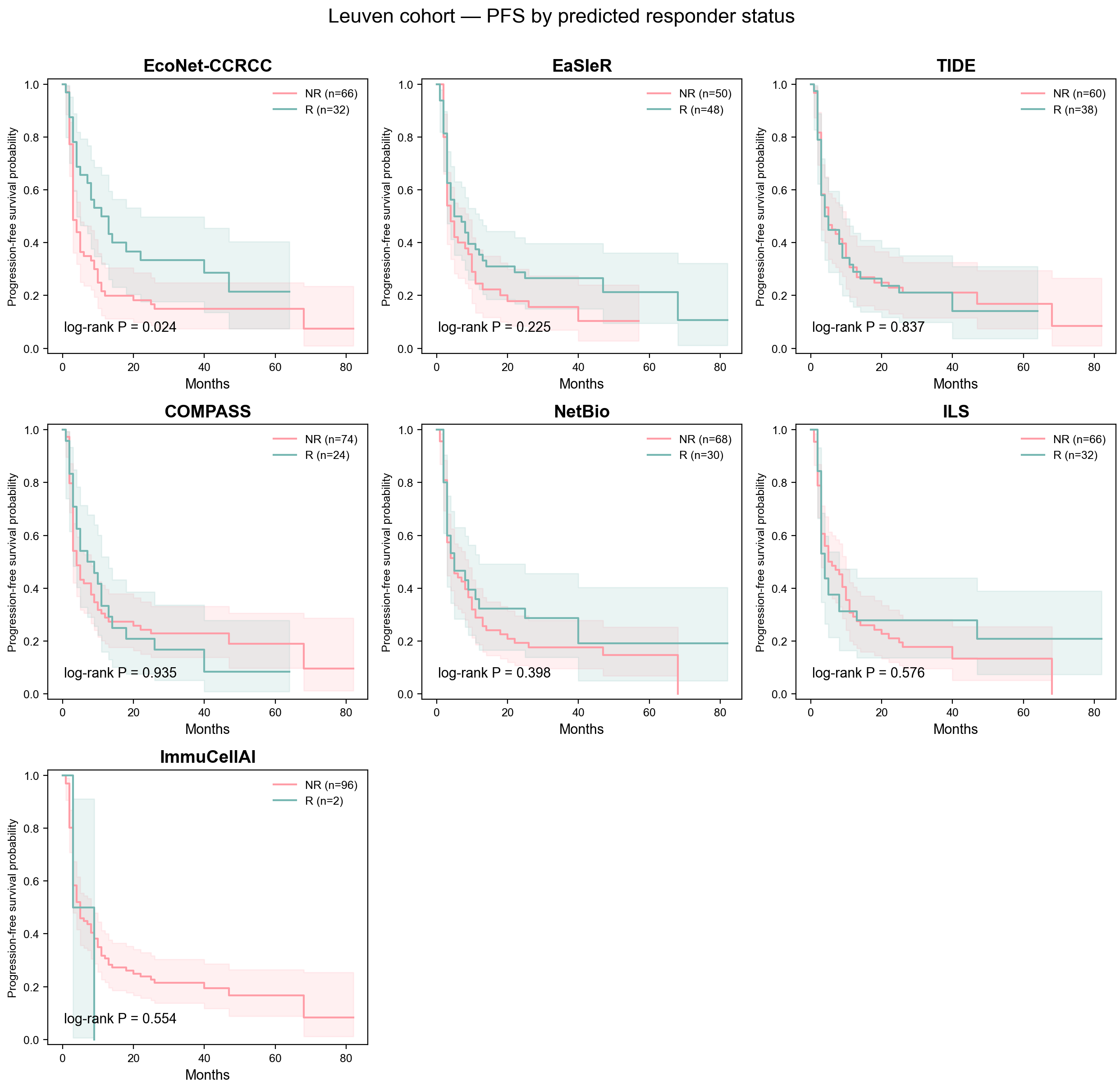

Supplementary Figure S12: **Progression-free survival analysis of the Leuven cohort stratified by predicted response across methods.** Each panel shows the Kaplan–Meier estimate comparing predicted responders and non-responders from one prediction method, with log-rank $P$-value annotated.

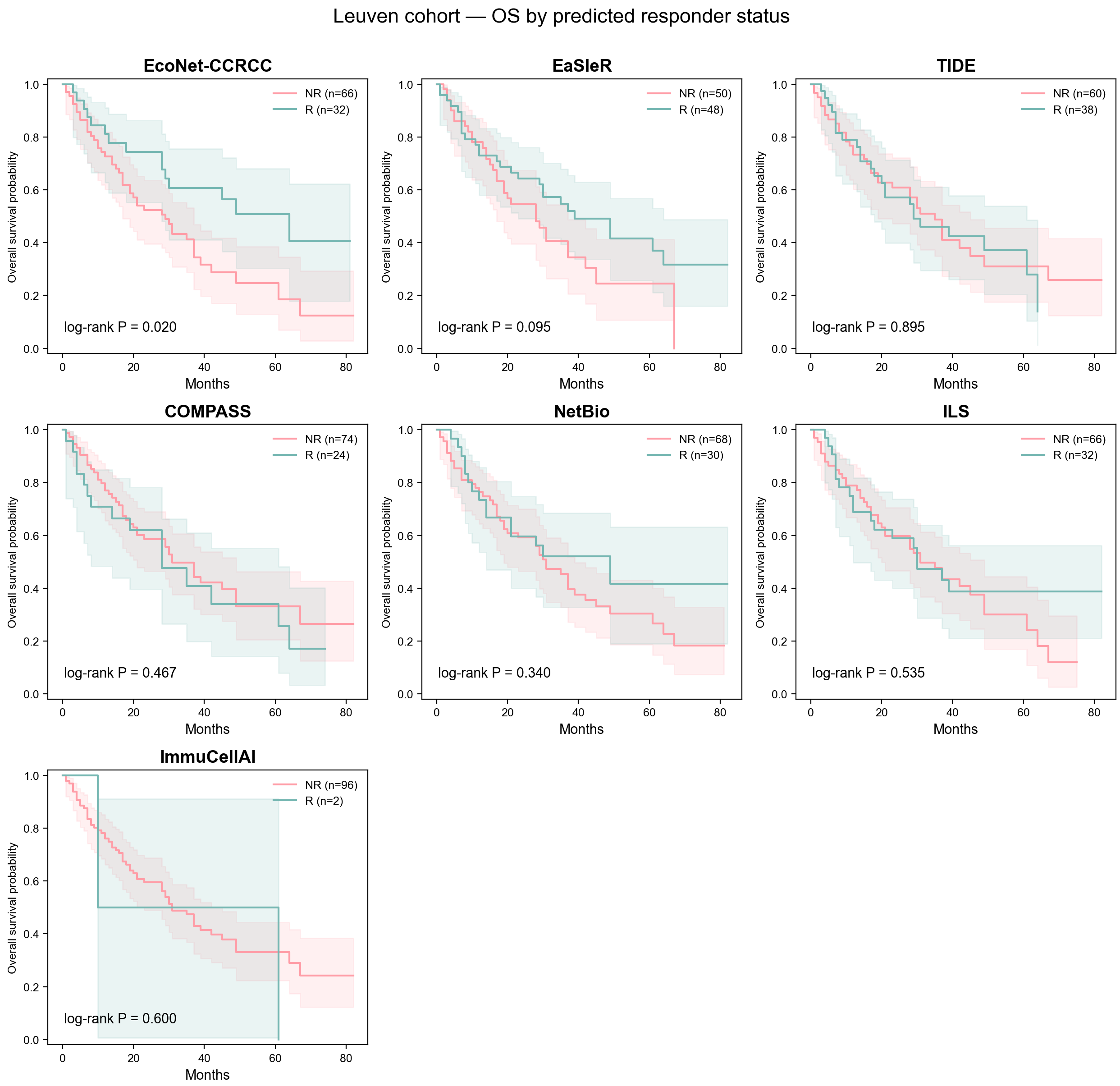

Supplementary Figure S13: **Overall survival analysis of the Leuven cohort stratified by predicted response across methods.** Each panel shows the Kaplan–Meier estimate comparing predicted responders and non-responders from one prediction method, with log-rank $P$-value annotated.

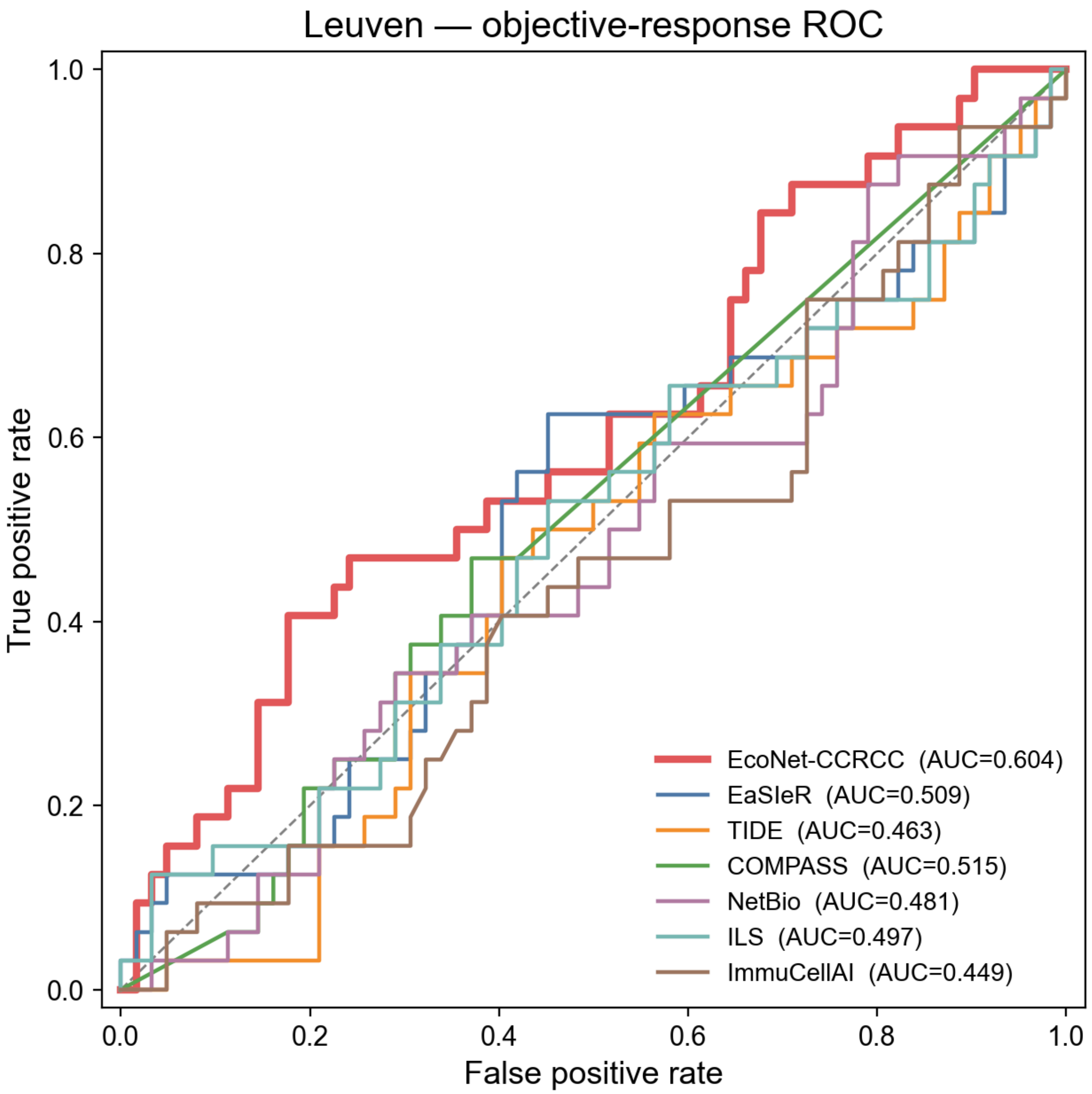

Supplementary Figure S14: **ROC curves for response prediction across methods on the Leuven cohort.** Each curve represents one prediction method, with AUC values annotated in the legend.

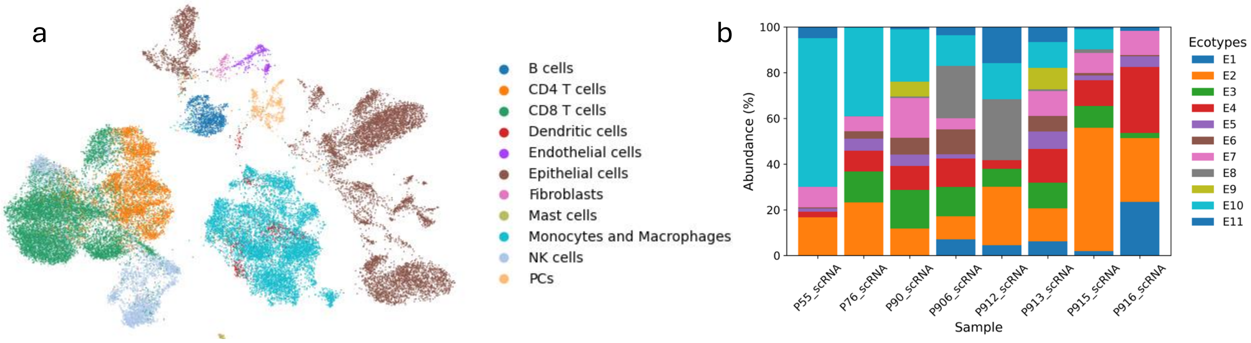

Supplementary Figure S15: **Ecotype abundance across ccRCC scRNA-seq samples in the SCP1288 dataset. a.** The UMAP plot shows the cells in the SCP1288 dataset colored by cell type. **b.** The cumulative bar plot shows the ecotype abundance for each sample.

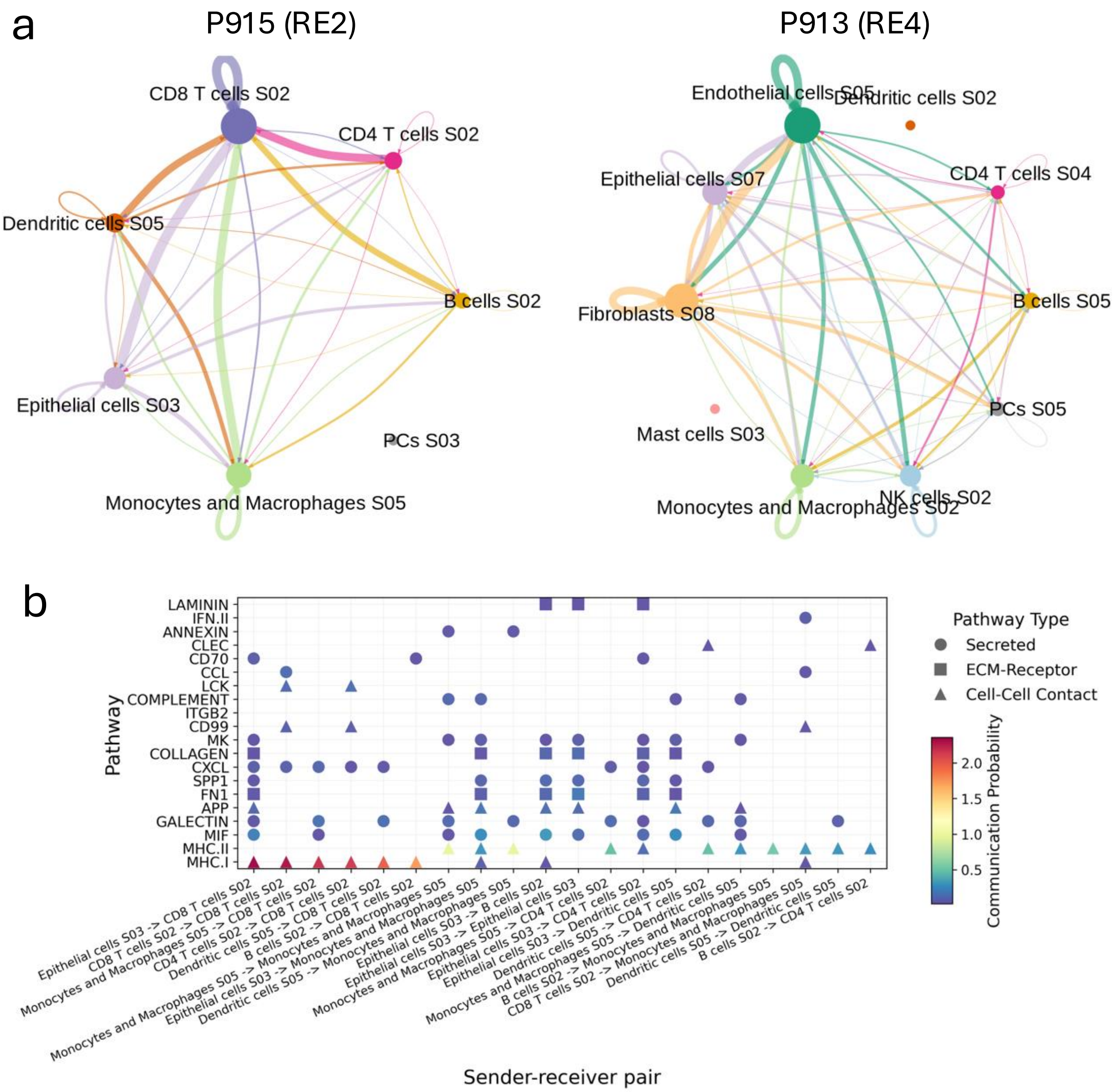

Supplementary Figure S16: **Cell-cell communication among cell states in RE2 and RE4.** **a**. Cell-cell communication networks reconstructed for representative samples P915 (assigned to RE2) and P913 (assigned to RE4). Nodes represent the corresponding cell states, and edge thickness reflects the inferred communication weight between sender and receiver populations. **b**. Intercellular communication among cell states in RE2. The dot plot summarizes pathway-level communication probabilities between cell states in sample P915. The dot size indicates communication probability, and the shape indicates pathway type.

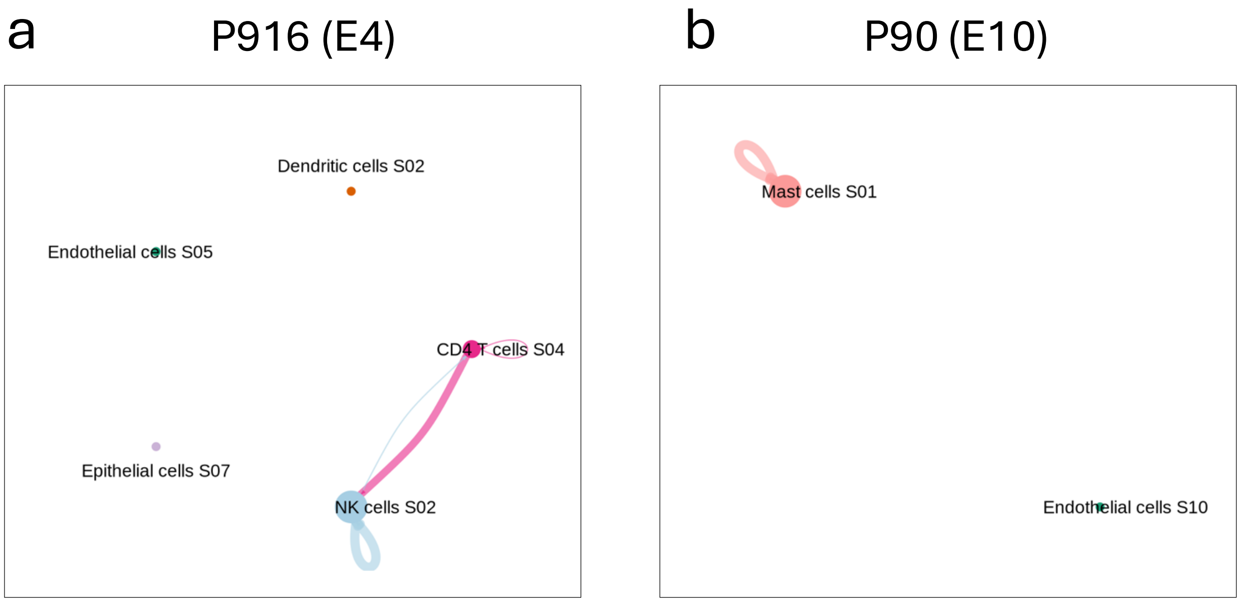

Supplementary Figure S17: **Cell-cell communication networks reconstructed for representative samples P916 (assigned to E4) and P90 (assigned to E10).** Nodes represent the corresponding cell states, and edge thickness reflects the inferred communication weight between sender and receiver populations. For P55 (assigned to E10), P76 (assigned to E10), and P912 (assigned to E8), no significant interactions were detected among the selected states.

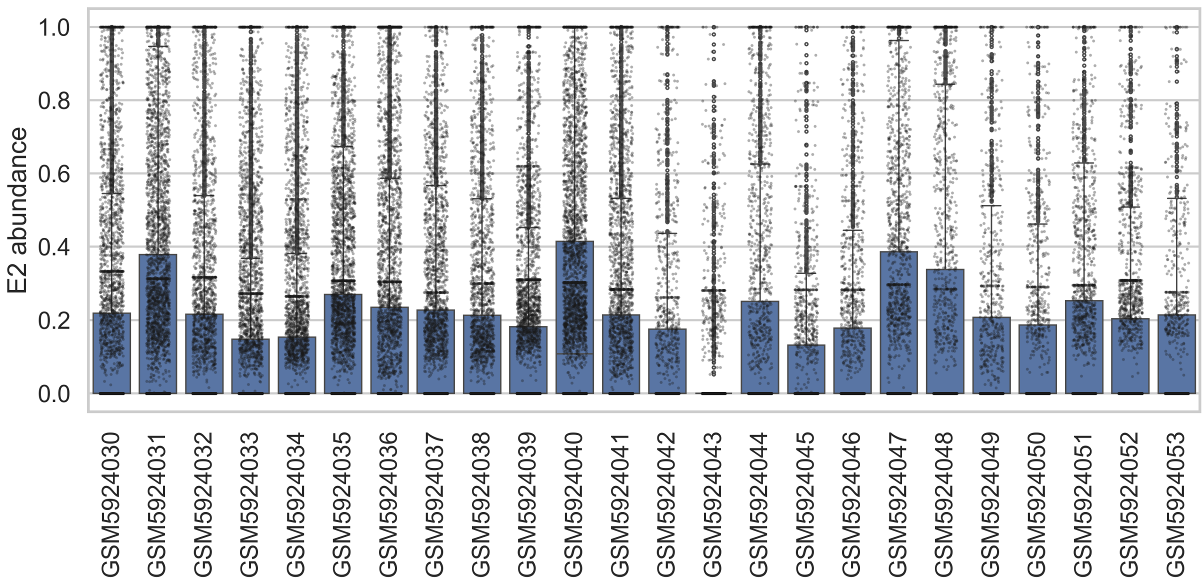

Supplementary Figure S18: **Distributions of RE2 abundance in ccRCC 10x Visium samples.** The boxplot shows the RE2 abundance across the spots for each of the 24 samples [31]; lines indicate medians, boxes indicate IQRs, and whiskers indicate 1.5×IQR.

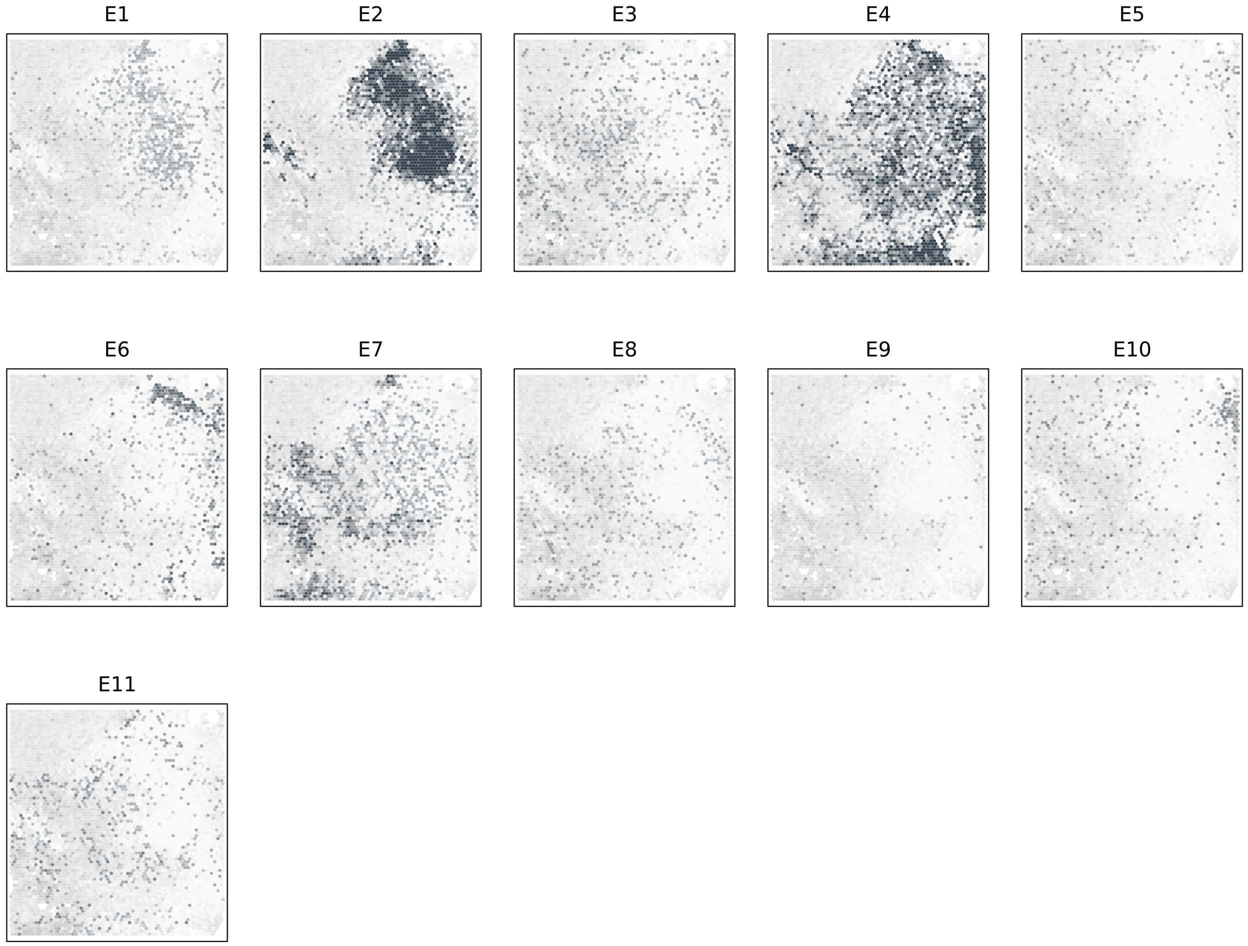

Supplementary Figure S19: **Recovery of ecotypes in spatial transcriptomics data (GSM5924036).** Dot intensity reflects the abundance of ecotypes, overlaid on tissue sections with epithelial regions marked in grey.

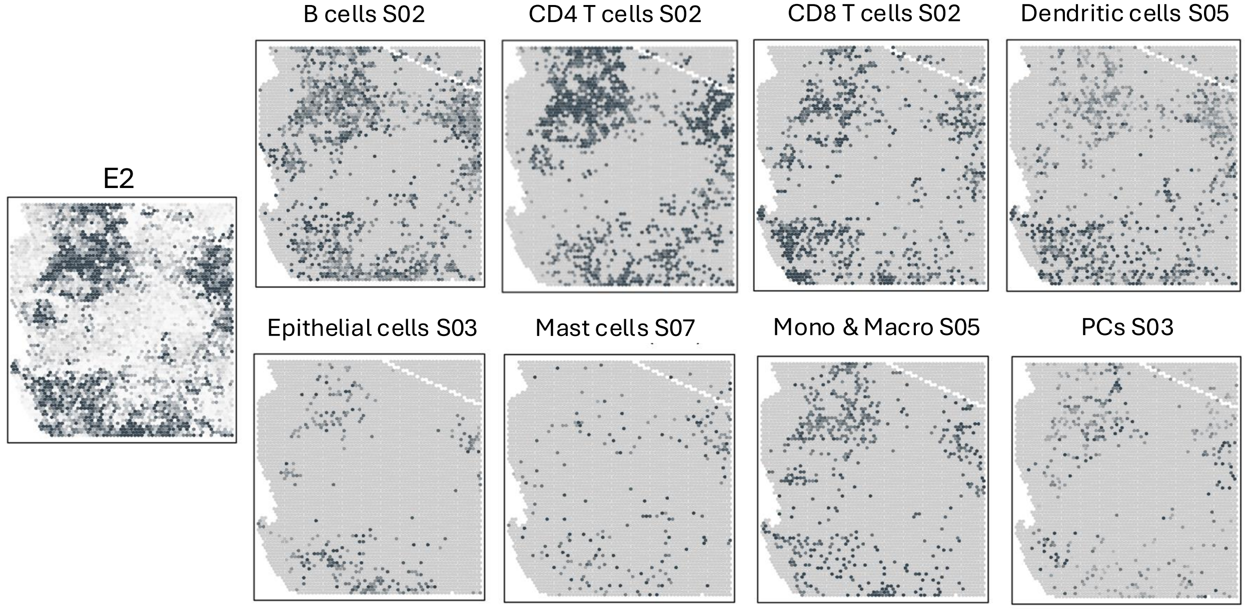

Supplementary Figure S20: **Recovery of RE2 and its cell states in spatial transcriptomics data (GSM5924031).** Dot intensity reflects the abundance of RE2 and corresponding cell states, overlaid on tissue sections with epithelial regions marked in grey.

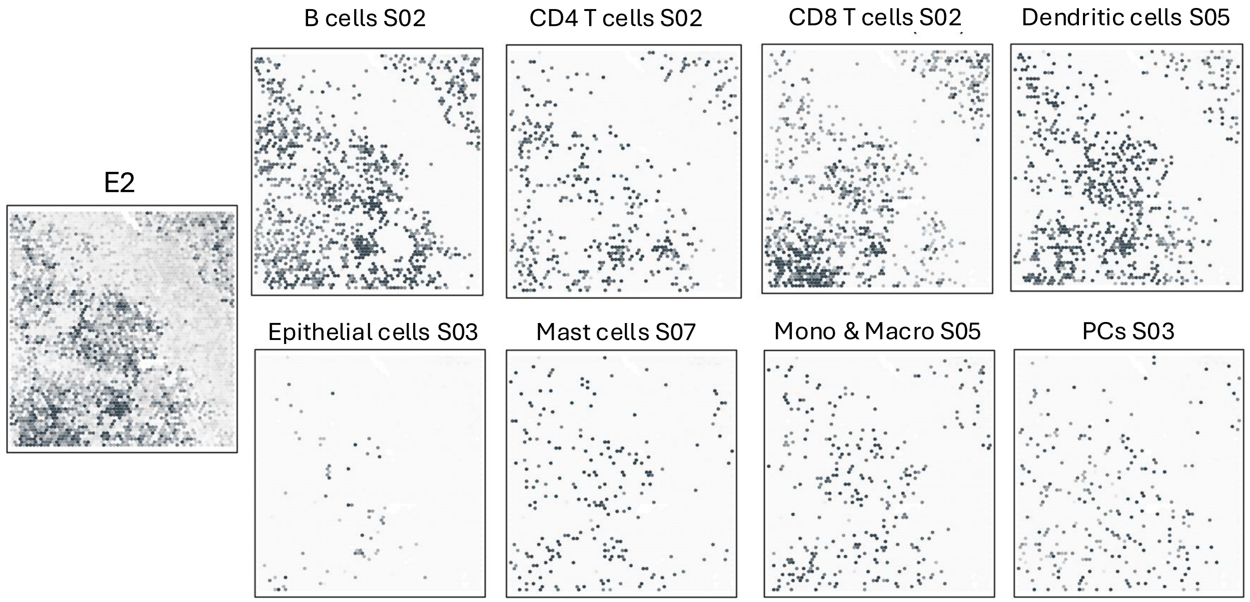

Supplementary Figure S21: **Recovery of RE2 and its cell states in spatial transcriptomics data (GSM5924035).** Dot intensity reflects the abundance of RE2 and corresponding cell states, overlaid on tissue sections with epithelial regions marked in grey.

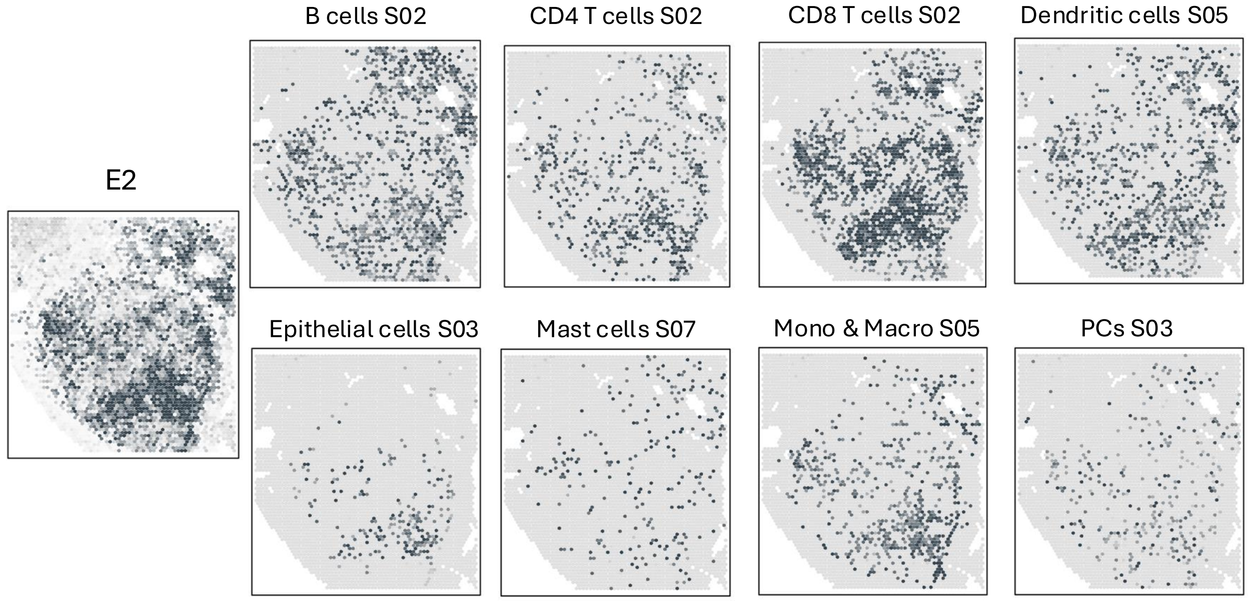

Supplementary Figure S22: **Recovery of RE2 and its cell states in spatial transcriptomics data (GSM5924040).** Dot intensity reflects the abundance of RE2 and corresponding cell states, overlaid on tissue sections with epithelial regions marked in grey.

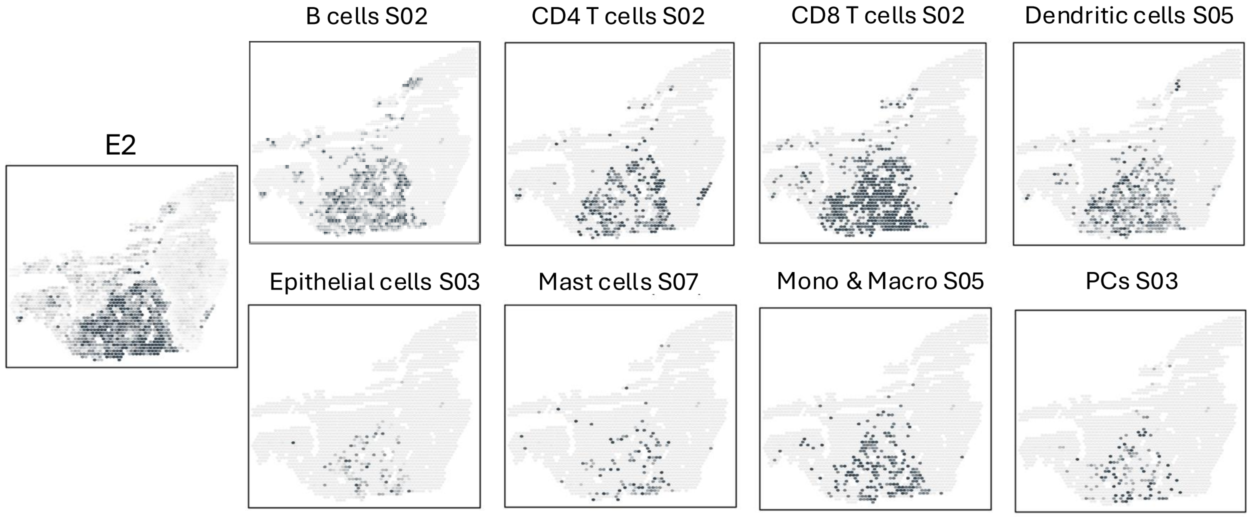

Supplementary Figure S23: **Recovery of RE2 and its cell states in spatial transcriptomics data (GSM5924044).** Dot intensity reflects the abundance of RE2 and corresponding cell states, overlaid on tissue sections with epithelial regions marked in grey.

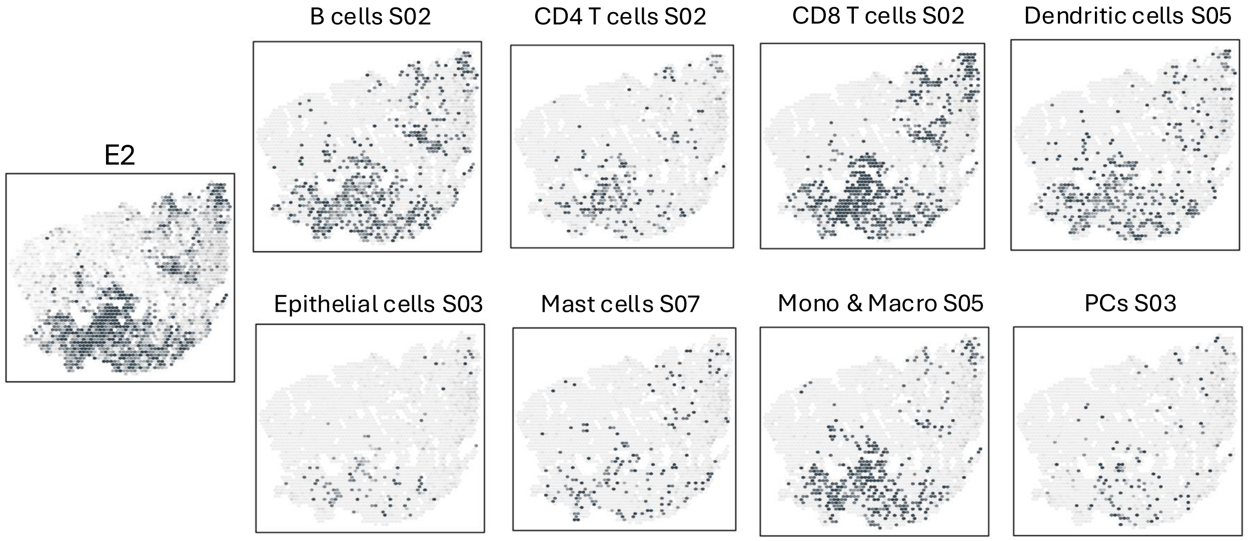

Supplementary Figure S24: **Recovery of RE2 and its cell states in spatial transcriptomics data (GSM5924047).** Dot intensity reflects the abundance of RE2 and corresponding cell states, overlaid on tissue sections with epithelial regions marked in grey.

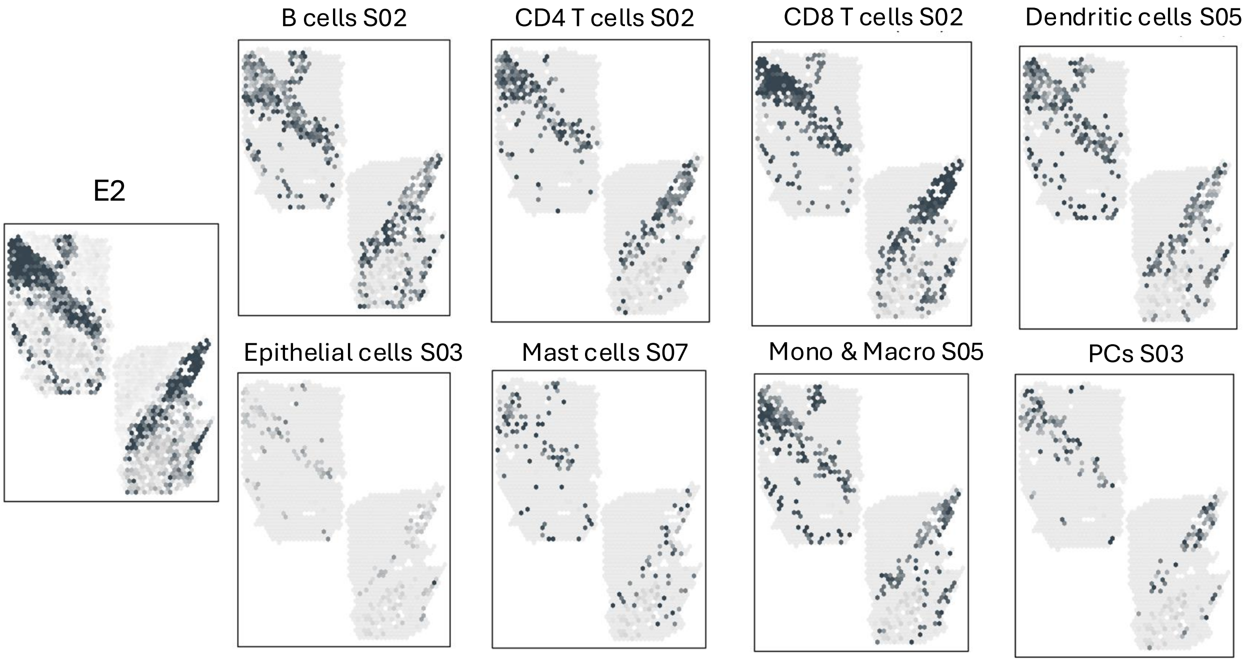

Supplementary Figure S25: **Recovery of RE2 and its cell states in spatial transcriptomics data (GSM5924048).** Dot intensity reflects the abundance of RE2 and corresponding cell states, overlaid on tissue sections with epithelial regions marked in grey.

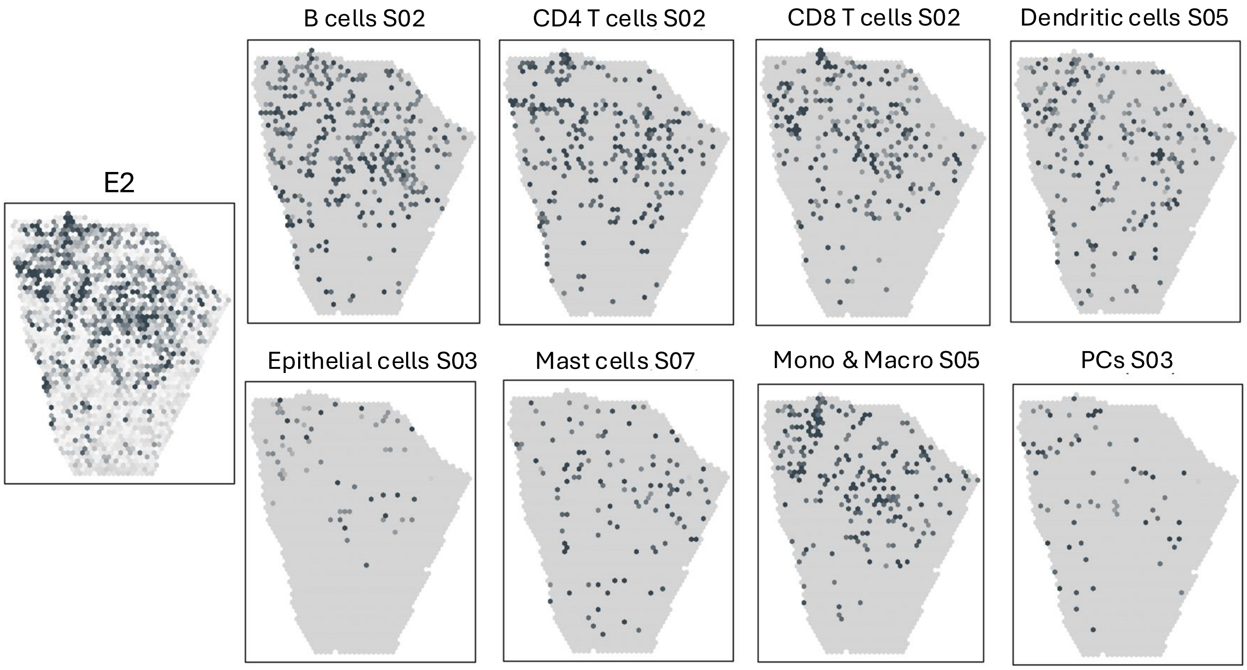

Supplementary Figure S26: **Recovery of RE2 and its cell states in spatial transcriptomics data (GSM5924051).** Dot intensity reflects the abundance of RE2 and corresponding cell states, overlaid on tissue sections with epithelial regions marked in grey.

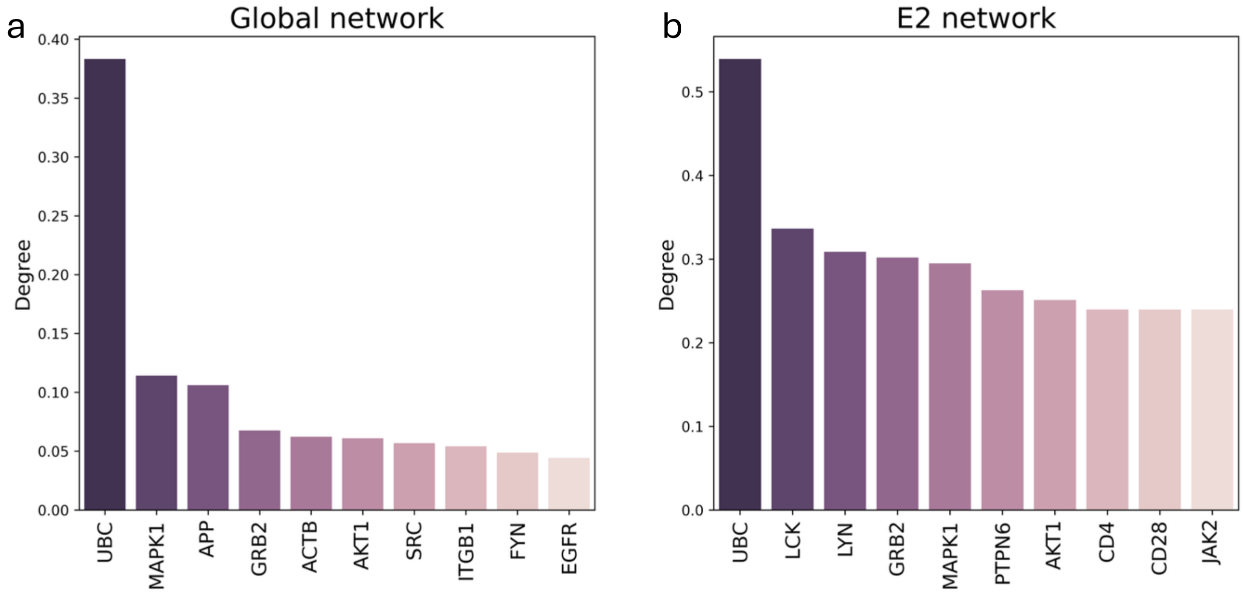

Supplementary Figure S27: **Hub genes in the global and RE2 ccRCC signaling networks. a.** Top ten genes ranked by degree centrality in the global ccRCC TME network. **b.** Top ten genes ranked by degree centrality in the RE2-specific network.

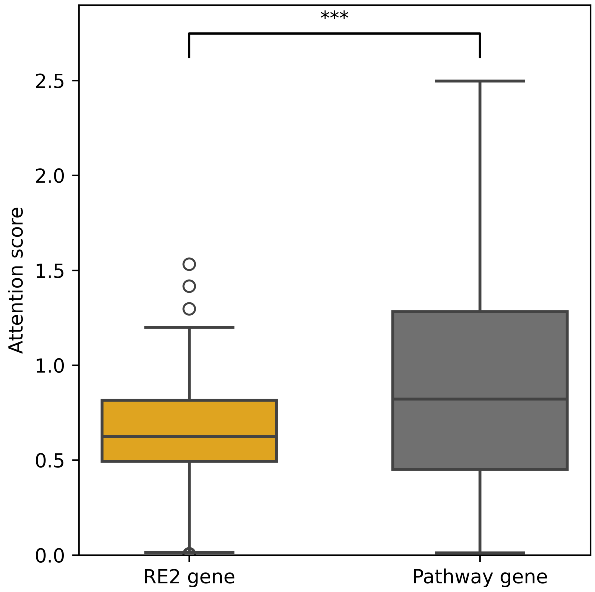

Supplementary Figure S28: **Gene attention scores for RE2 marker and pathway genes.** Comparison of EcoNet attention scores between RE2 marker genes and pathway genes within the RE2 network ($P = 3.9 \times 10^{-4}$).

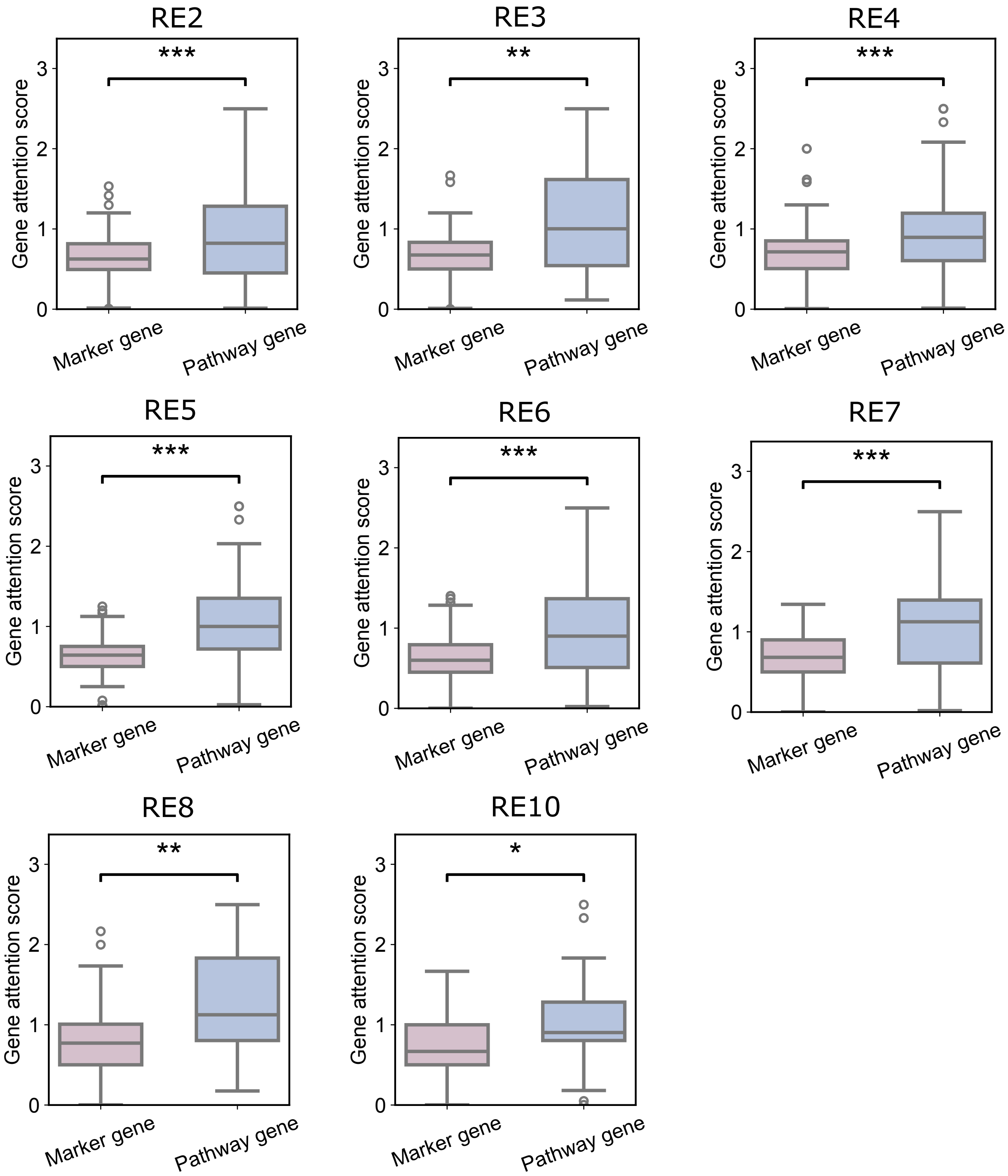

Supplementary Figure S29: **Comparison of attention scores between ecotype marker genes and pathway genes for each CCRCC ecotype network.** RE1, RE9, and RE11 networks were excluded, which lacked pathway genes.

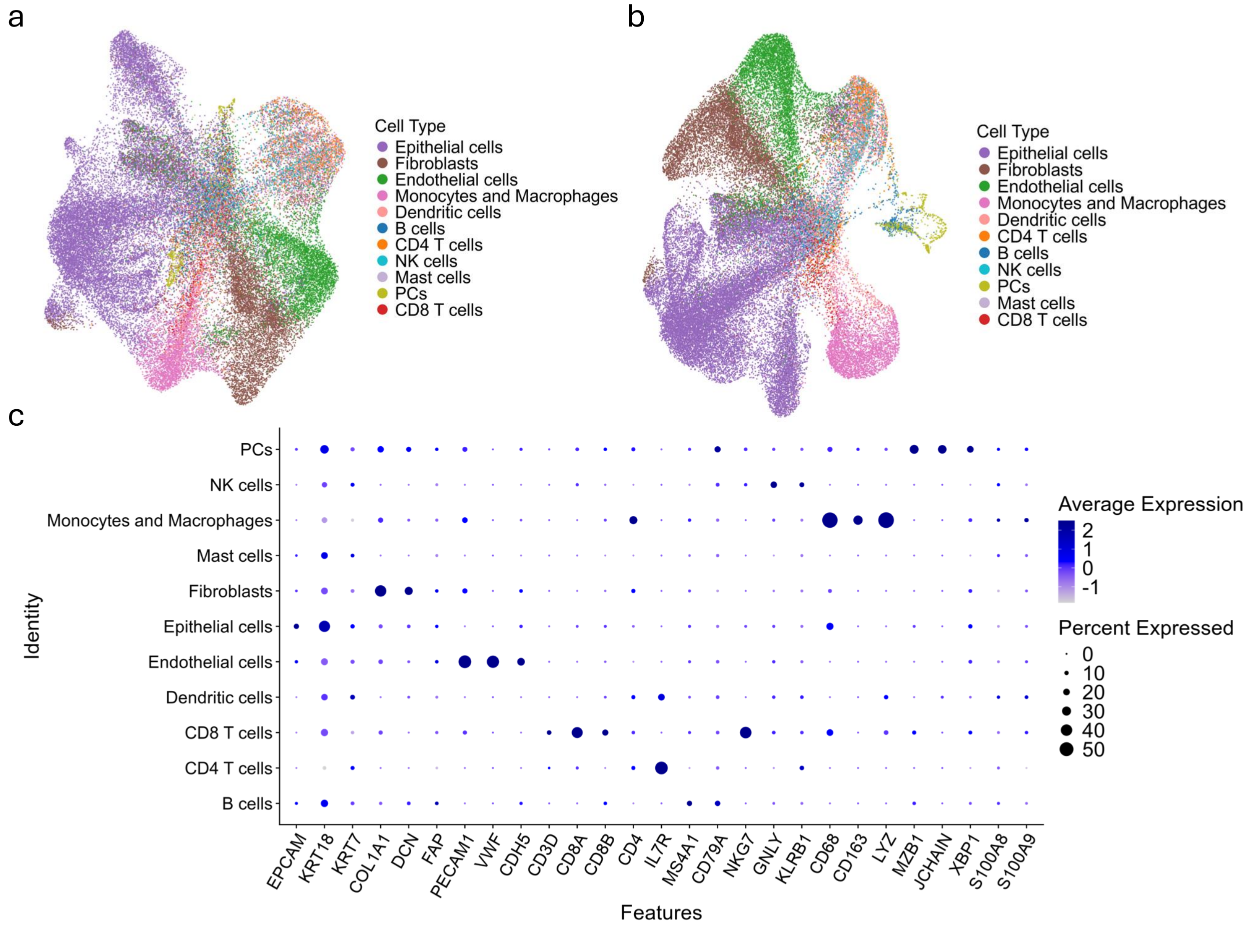

Supplementary Figure S30: **DeSpotX decontamination performance on ccRCC CosMx data.** **a**. UMAP of CosMx cells before decontamination, colored by cell type. **b**. UMAP of CosMx cells after three iterations of DeSpotX decontamination and re-annotation, showing improved separation of cell types. **c**. Dot plot of canonical marker gene expression across cell types after decontamination. Dot size indicates the percentage of cells expressing each gene, and color indicates the average expression level.

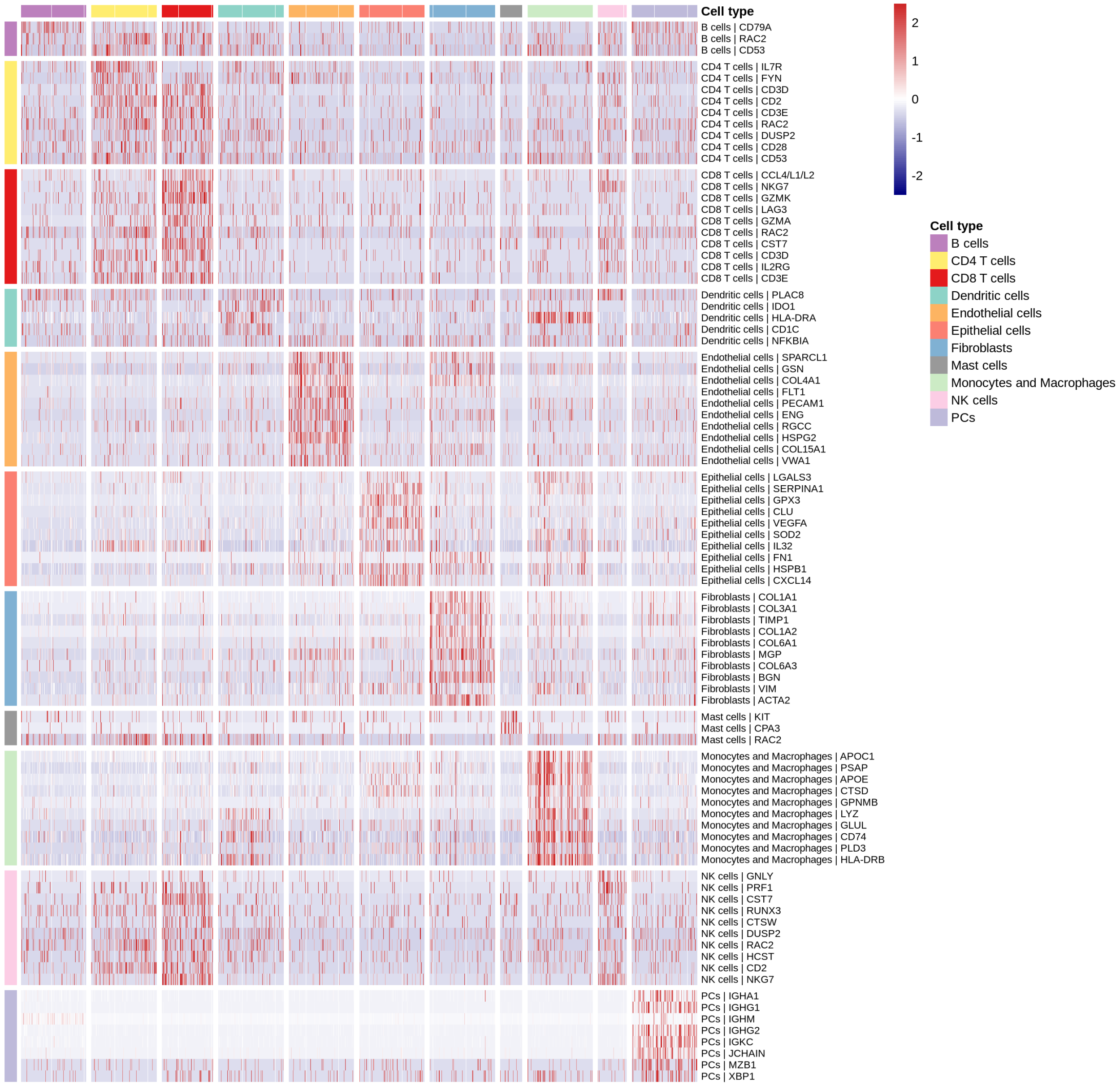

Supplementary Figure S31: **Heatmap of $z$-scored expression for top 10 positive marker genes per cell type in CCRCC CosMx data.** To evaluate the annotation results, we identified the marker genes for each cell type with Seurat (`FindAllMarkers`; $\log_2$ FC $> 0.25$, `min.pct` $> 0.25$). Columns are randomly sampled cells ($\leq$ 1000 per type), grouped by cell type; rows are marker genes labeled "Cell type | Gene".

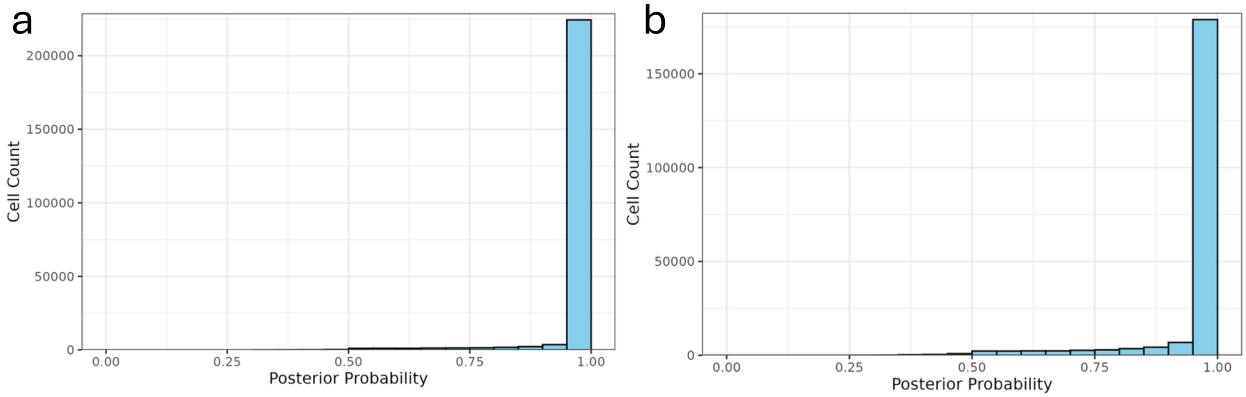

Supplementary Figure S32: **Distributions of posterior assignment probabilities from Insitutype.** Bar plots show the distributions of posterior assignment probability for the cells on the two slides (**a** and **b**). The cells with the posterior assignment probability $> 0.8$ were retained for subsequent analyses.

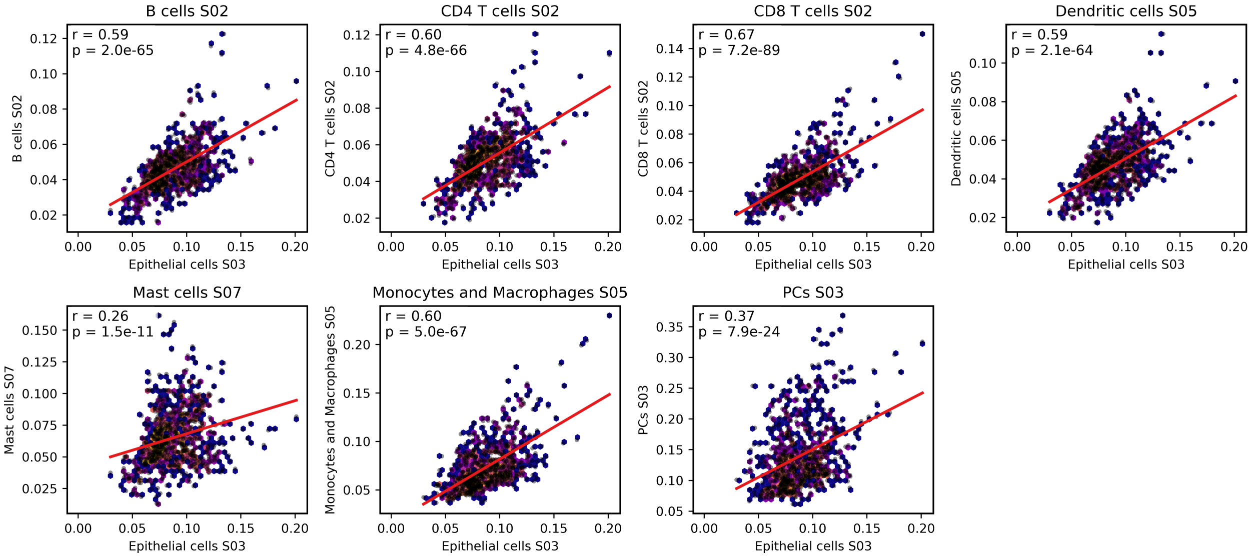

Supplementary Figure S33: **Correlation of marker gene expressions between epithelial cell state and other cell states in RE2 across FOVs.** Each panel shows the pairwise relationship between the mean marker gene expression of epithelial cells S03 and that of another RE2 cell state across fields of view (FOVs). The red line shows the linear regression fit; inset values denote the Pearson correlation coefficient ($r$) and $P$-value ($p$).

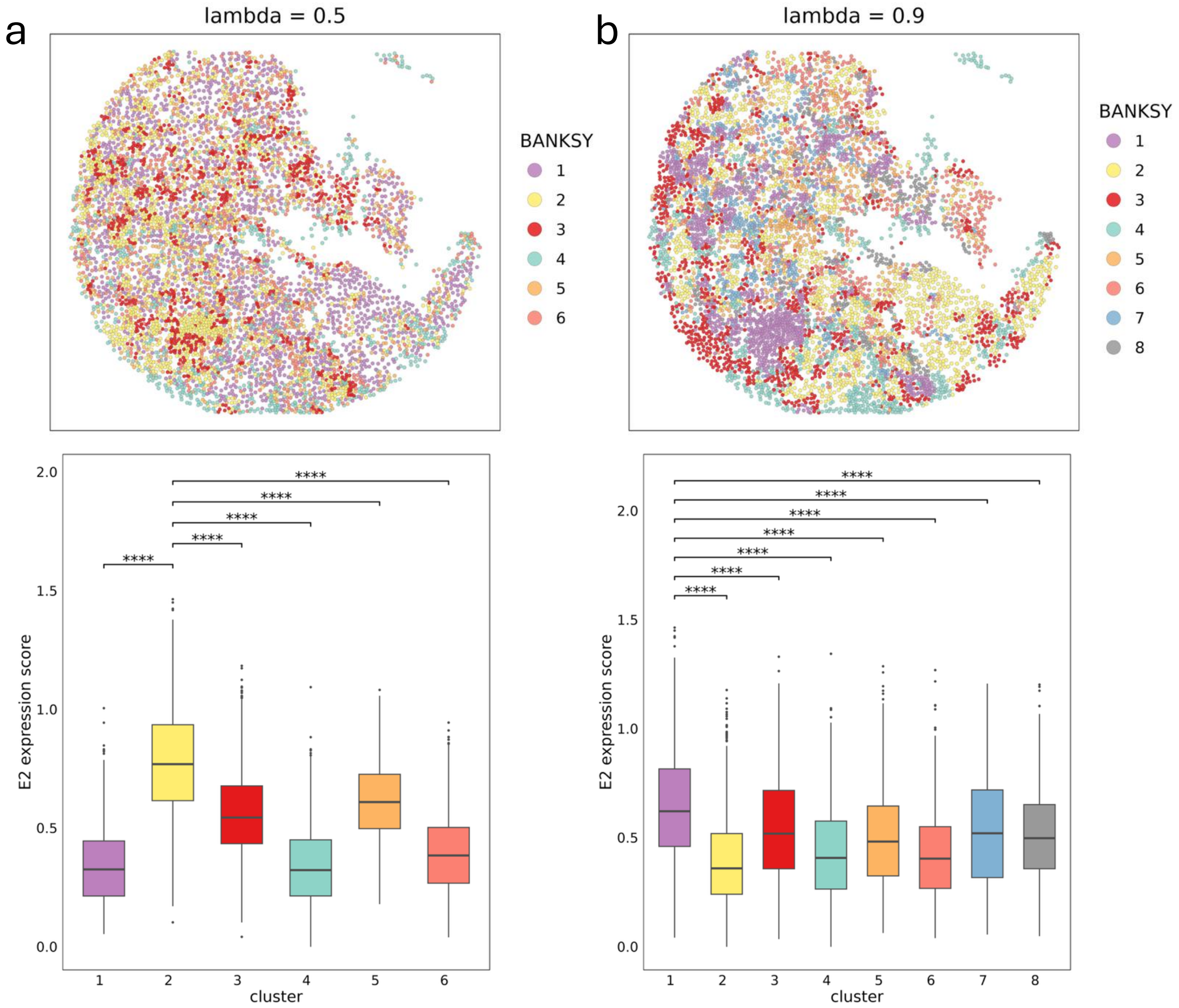

Supplementary Figure S34: **Distribution of RE2 expression scores across BANKSY-defined spatial clusters.** For BANKSY-defined clusters with **(a)** $\lambda = 0.5$, and **(b)** $\lambda = 0.9$, we show the spatial maps for the representative sample (top) and RE2 marker expression distributions across the spatial clusters (bottom) ($^{****}P < 0.0001$).

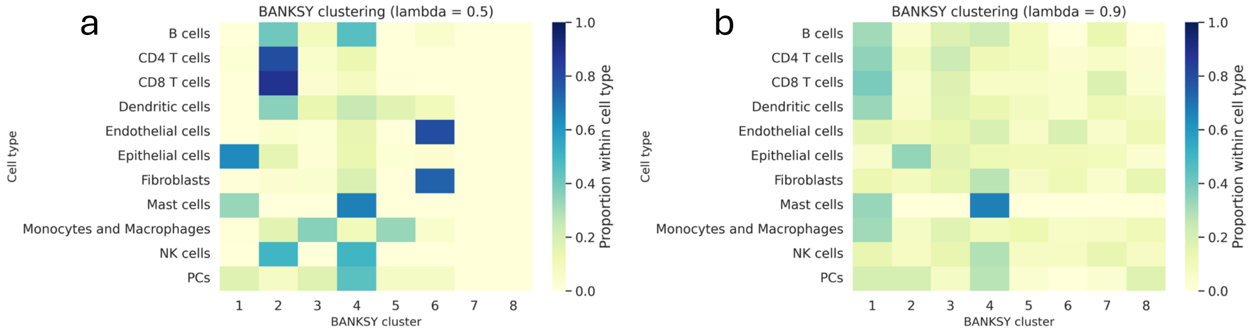

Supplementary Figure S35: **Distribution of cell types across BANKSY-defined spatial clusters.** The heatmap shows the proportional distribution of cell types across BANKSY-defined spatial clusters with **(a)** $\lambda = 0.5$, and **(b)** $\lambda = 0.9$ in the representative sample. Each row represents a cell type, and each column corresponds to a spatial cluster. Color intensity indicates the proportion of each cell type within the cluster.

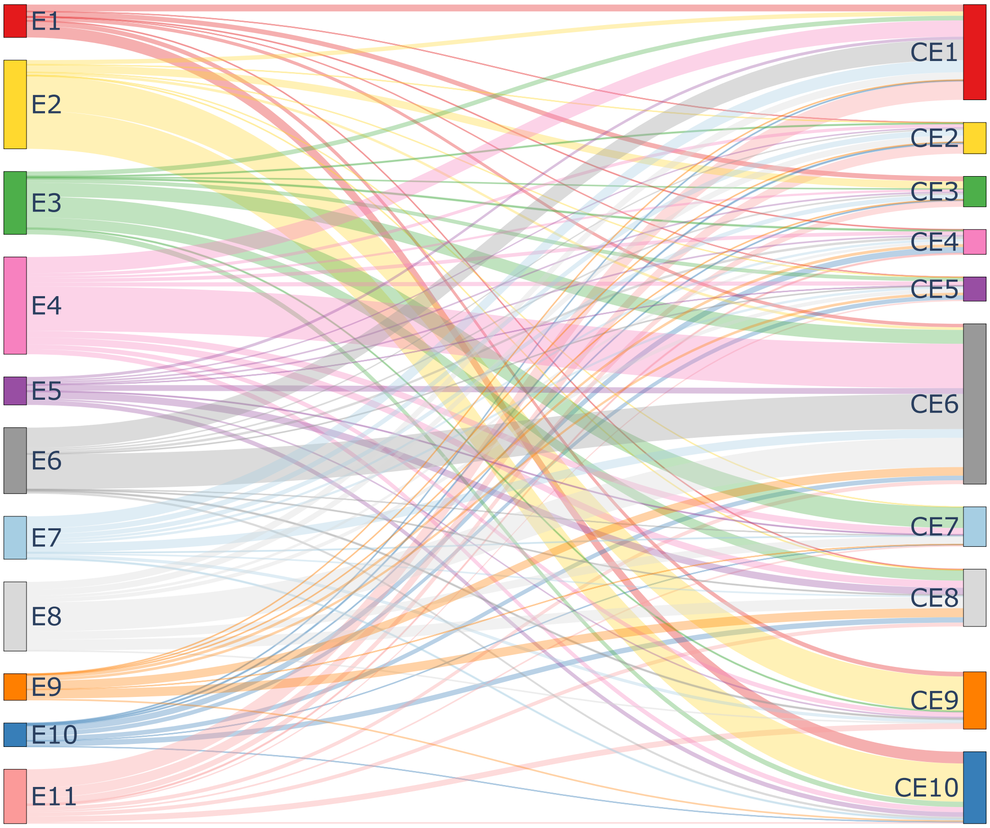

Supplementary Figure S36: **Overlap of marker genes between ccRCC ecotypes and pan-cancer ecotypes.** Each node on the left represents a ccRCC ecotype, whereas each node on the right represents a pan-cancer ecotype [2]. The width of each link corresponds to the number of shared marker genes.

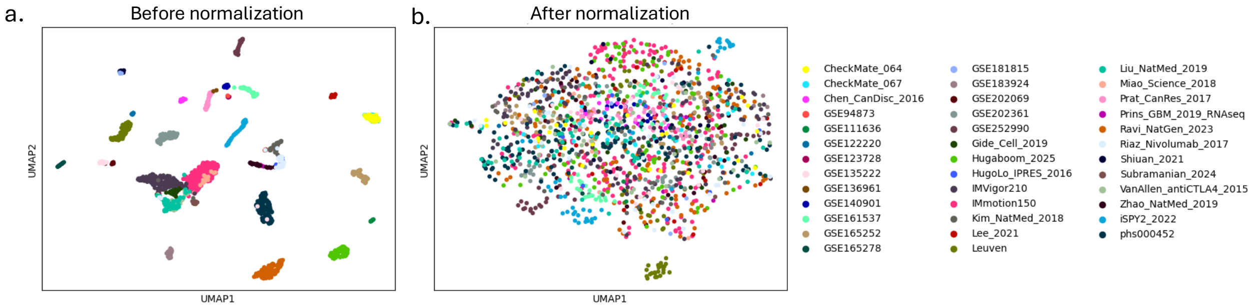

Supplementary Figure S37: **UMAP plots of pan-cancer immunotherapy cohorts before and after $z$-score normalization.** UMAP embeddings are shown for **(a)** unnormalized data and **(b)** data after $z$-scoring. The normalization harmonizes expression distributions across cohorts.

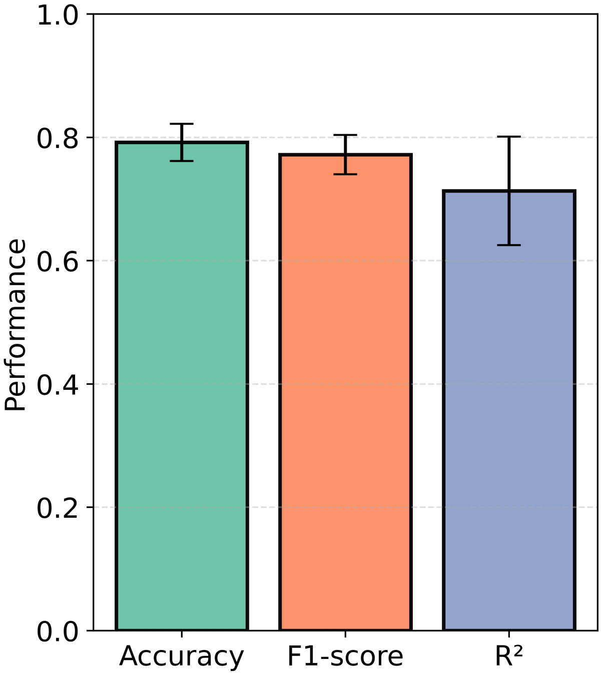

Supplementary Figure S38: **Five-fold cross-validation performance of the representation learning module.** Bars show mean ± standard deviation of Accuracy, F1-score, and $R^2$ across folds.

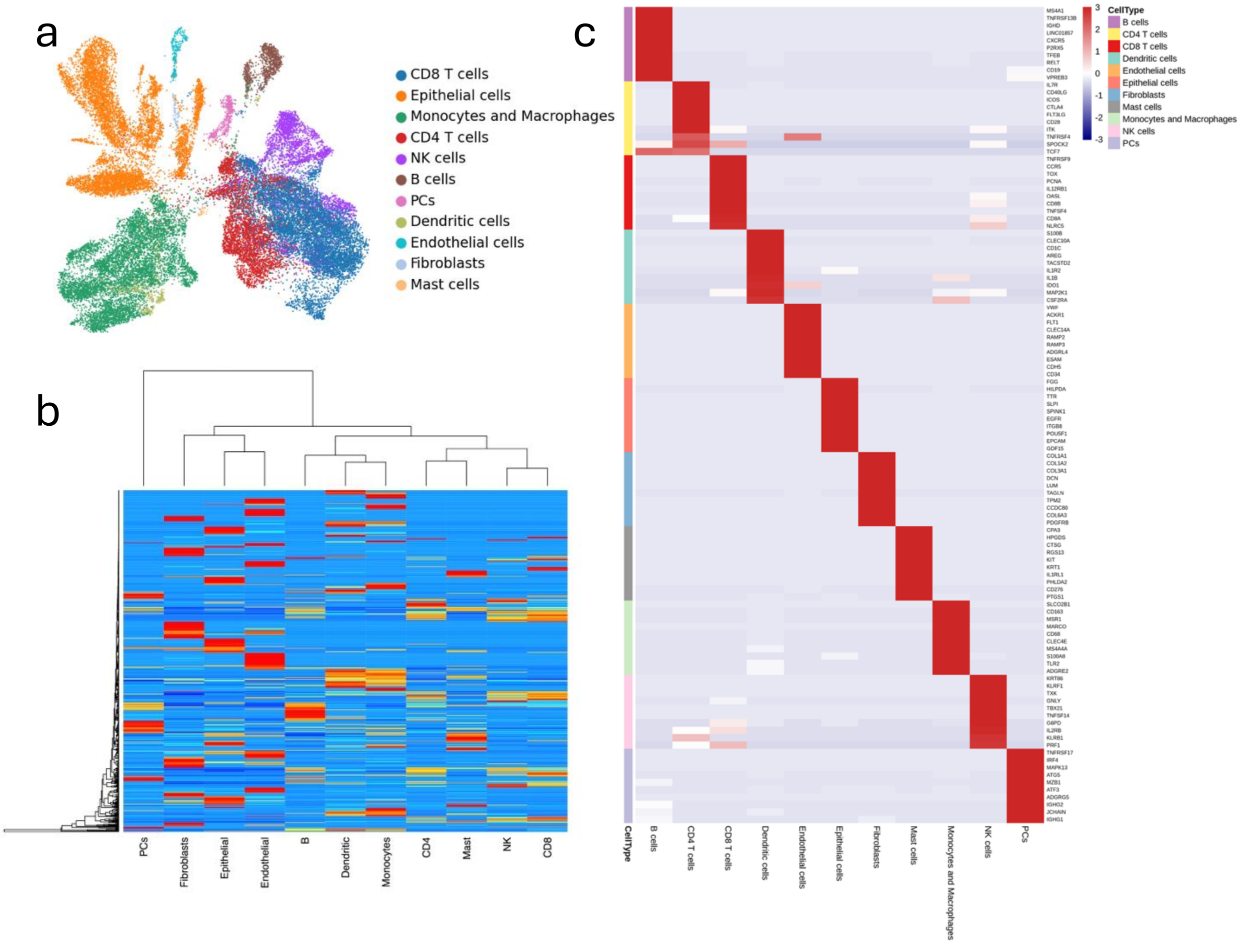

Supplementary Figure S39: **CIBERSORTx-derived ccRCC signature matrix.** **(a)** UMAP of SCP1288 cells computed using genes on the CosMx panel; points are colored by cell type. **(b)** Heatmap of the reference signature matrix used for supervised cell typing, with 11 cell types as columns and 583 genes as rows. **(c)** Heatmap of $z$-scores for the top ten marker genes per cell type derived from the signature matrix, ranked by $\log_2$fold-change. Rows represent genes, columns represent cell types, and the left bar denotes the assigned cell type for each gene.
